# I/σI *vs* {Rmerg, Rmeas, Rpim, CC1/2} for Crystal Diffraction Data Quality Evaluation

**DOI:** 10.1101/2024.12.10.627855

**Authors:** Zheng-Qing Fu, Brian V. Geisbrecht, Samuel Bouyain, Fred Dyda, John Chrzas, Palani Kandavelu, Darcie J. Miller, Bi-Cheng Wang

## Abstract

X-ray crystal diffraction has provided atomic-level structural information on biological macromolecules. Data quality determines the reliability of structural models. In most cases, multiple data sets are available from different crystals and/or collected with different experimental settings. Reliable metrics are critical to rank and select the data set with the highest quality. Many measures have been created or modified for data quality evaluation. However, some are duplicate in functionality, and some are likely misused due to misunderstanding, which causes confusion or problems, especially at synchrotron beamlines where experiments proceed quickly. In this work, these measures are studied through both theoretical analysis and experimental data with various characteristics, which demonstrated that: 1). {Rmerg, Rmeas, Rpim, CC1/2} all measure the equivalence of reflections, and the low-shell values of these metrics can be used as reliable indicators for correctness (or trueness) of Laue symmetry; 2). High-shell I/*σ*I is a reliable and better indicator to select resolution cutoff while the overall value measures the overall strength of the data.

## Introduction

Macromolecular structures have provided a powerful tool for understanding biological function and have played a pivotal role in design of pharmaceuticals. X-ray diffraction, with a century-long and continuous development, has become a primary method to solve 3D structures of macromolecules in terms of number of structures, apparent resolution, and quality. It will continue adding structures into PDB database (Berman et al., 2000; Berman et al., 2003, Burley et al., 2023), which in turn provides more reliable templates for molecular replacement or docking into maps of Cryo-EM (Dubochet et al., 2018; Frank, 2009; Henderson et al., 1990; Cheng et al., 2015). As in any experiment, an effective and reliable metric is critical to evaluate and select the best X-ray diffraction data set for structural solution. Historically, I/*σ*I and/or Rmerg (a.k.a Rsym) were used to evaluate data quality. Rmeas (Diederichs et al., 1997) and Rpim (Weiss et al., 1997) were later introduced to try improving the multiplicity-dependency of Rmerg. Traditionally, resolution cutoffs are selected by I/*σ*I≥2.0 and/or {Rmerg, Rmeas, Rpim}≤40%, 50%, 60% etc. in the highest resolution shell. More recent studies showed that this may be conservative as it could exclude weak but still useful reflections within the higher resolution shells; this resulted in the introduction of the index CC1/2 (Karplus et al., 2012; Karplus et al., 2015). However, CC1/2 does not have a single value as the threshold that is generally applicable to different cases. Different values (such as 0.1, 0.2, 0.3, 0.4, etc.) have been suggested to select data and used as data processing programs’ defaults, or defaults customized by users at different sites, which causes confusion in practice during data collection and processing. Although these different measures have been developed and used to evaluate and select data from X-ray crystal diffraction experiments, what is the best and how to select the best data set remains debatable. This situation is magnified when using synchrotron beamlines, where experiments proceed at a fast pace. In this work, their meanings, capabilities, correlations, and effectiveness are studied through theoretical analysis and experimental data with various characteristics to clarify the confusion and potential misuses, and thus to identify the most reliable measures for evaluating data quality from different aspects.

## Methods

Some background knowledge is summarized here first for easier reading and for helping understand the subjects involved. Data from diffraction experiments are reduced to a set of unique reflections by minimizing differences among the equivalent reflections through target function (1) during the last stage of data processing (Hamilton et al., 1965; Otwinowski et al., 1997; Kabsch, 2010; Leslie et al., 2007; Evans, 2011; Fu, 2005; Winter et al., 2018; Giacovazzo, 1992),

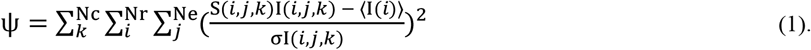

The discrepancy index associated with the minimization process can be calculated by

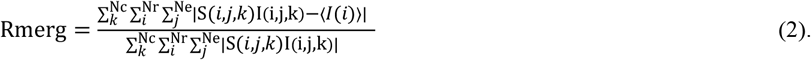

Here *j* runs through the equivalent reflections (denoted as including both repeated measurements of same reflection and those related by Laue group symmetry of the crystal, hereafter in this article for simplicity), *i* runs through the unique reflections, and *k* runs through the number of data sets if multiple data sets are involved. I(*i,j,k*), *σ*I(*i,j,k*), are the observed intensity and estimated error, and S(*i,j,k*) are scaling factors. <I(*i*)> (denoted as I(*i*) for simplicity hereafter) and *σ*I(*i*) are the merged intensity from equivalents and the estimated error or noise for each unique reflection **H**(*i*). The result of data processing is the reduced data set {**H**(*i*), I(*i*), *σ*I(*i*) | *i*=1, N_r_} consisting of the unique reflections up to a selected resolution (a.k.a. resolution cutoff). The number of unique reflections N_r_ is determined by the resolution cutoff selected.

As shown in equation (1), the fundamental of raw data reduction from a diffraction experiment is to minimize the differences among intensities of equivalent reflections. Rmerg, as the standard discrepancy index associated with the minimization, provides a measure of quality of the data set in terms of equivalency or internal consistency (Evans et al., 2013).

The second quality indicator of the reduced data set is the merged <I/*σ*I> (denoted as I/*σ*I hereafter for simplicity). I/*σ*I is necessary as it provides quality evaluation from a different aspect, namely the signal-to-noise ratio (SNR as commonly denoted). SNR is fundamental to all experiments as it compares the level of a desired signal to the level of background noise, which would provide a measure of data quality or reliability.

The third fundamental quality indicator is the completeness (denoted as *Compl* hereafter for simplicity), because the electron density map (commonly calculated as a weighted 2Fo-Fc map during structure refinement) is Fourier transform of the reflections (Giacovazzo, 1992),

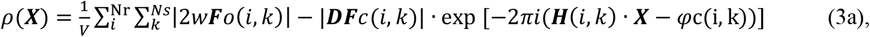

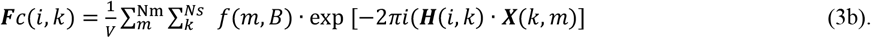

Here *i* runs through the unique reflections {**H**(*i*), I(*i*), | *i*=1, N_r_}, and *k* runs through symmetry operations. *φ*_c_(i,k) is the phase of reflection structure factor. 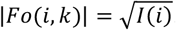 runs through all atoms in the asymmetric unit. 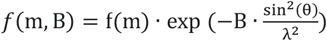 with f(m) as the atomic form factor and θ the diffraction angle. B is the temperature factor that is evaluated during structure refinement (Afonine et al., 2012; Murshudov et al., 2011). **X**(k,m) are the coordinates of atom (k,m). **F**c(*i,k*) is the calculated structure factor using coordinates and B factors of all atoms in the structure. In most diffraction experiments, not all reflections are measurable, leading to N_r_ being less than N_rc_ (the theoretically expected number of unique reflections to a given resolution). If some reflections are missing, it could affect the quality of the electron density map and, thus, the structure model built. The ratio N_r_/N_rc_ directly measures the completeness of the reduced data set. In practice, 100% completeness is not required to calculate a tracible (or solvable) electron density map. But the lower it is, the less tracible the electron density would be. It is worth noting that missing wedge, anisotropy, or overloads can introduce systematic incompleteness and have different impacts than random incompleteness, which is not the subject of this study.

Theoretically, Rmerg, I/*σ*I, and *Compl* are the fundamental measures needed to evaluate the quality of a reduced data set, which would, in turn, determine the quality of the structure. However, since I(*i*) and *σ*I(*i*) are merged from equivalent reflections, the number *Ne* as shown in equation (1) will affect I(i) and *σ*I(i), leading to another measure ‘multiplicity’ (denoted as *Multi* hereafter) calculated as the average of *Ne*. Rmerg evaluated from equation (2) will be affected by *Multi*, such that the higher *Multi* the worse the Rmerg. As merging reduces the random noise, more measurements will typically lead to better quality of reduced data if no severe crystal decay occurred during the experiment. To try compensating for this effect, Rmeas (Diederichs et al., 1997) was introduced by adding 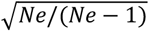 to equation (2). Intended to indicate the precision of the measurements, another alternative, Rpim was defined with a different factor 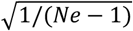 (Weiss et al., 1997).

CC1/2 is an indicator more recently introduced for selecting resolution cutoff (Karplus et al., 2012; Karplus et al., 2015). It is a Pearson correlation coefficient (Pearson, 1895) calculated from comparing the intensities of equivalents by randomly dividing them into two subsets, each containing a half. Since there is a lack of a generally applicable threshold, different values of CC1/2 have been suggested. In addition, when multiplicity is odd, this division excludes a set of reflections, leading to a CC1/2 evaluated with only a portion of the data. This partiality may not be an issue as it is the average for a resolution shell but could affect its estimation at high resolution shells when the in-shell multiplicity dramatically decreases as fewer reflections become measurable.

It is worth pointing out that, in addition to decay, there exist systematic errors from various sources in a diffraction experiment which can’t be cancelled out through merging. Some of them can be reduced through system calibration, whereas others may arise within the experiment such as radiation damage, crystal centering etc. During data processing, errors corrections are critical and applied typically through error models to evaluate both I(*i*) and *σ*I(*i*) in the reduced data (Hamilton et al., 1965; Otwinowski et al., 1997; Kabsch, 2010; Leslie et al., 2007; Evans et al., 2013; Fu, 2005; Winter et al., 2018; Giacovazzo, 1992). While the quality of a reduced data set typically improves as *Multi* increases, the magnitude of such improvement is conditional and can only be evaluated on a case-by-case basis. Details on how data quality improves with *Multi* in different cases is not the subject of this study.

Rmerg, Rmeas, Rpim derived from the minimization process of target function (1) are direct indicators of the differences among intensities of equivalent reflections. CC1/2 is also calculated from comparing the intensities of the equivalents, but in a different form. Pearson statistics (Pearson, 1895; Boslaugh, 2012) could provide a tool to shed light on the relationships, the effectiveness, and capabilities of these indicators, and may help identify the most reliable metrics to evaluate and select the best data set for structure solution. Pearson Correlation Coefficient (denotated as pcc hereafter) is a coefficient that measures linear correlation between two sets of data, U and W. It is a normalized ratio between the covariance of two variables and the product of their standard deviations, ranging between -1 and 1 inclusive. Its sign is determined by the regression slope: positive implies that W increases while U increases; negative means W increases as U decreases. The pcc is widely used as a powerful tool to investigate on the unknown covariance of two variables. The absolute value |pcc| measures the strength how closely the two variables are linearly correlated, which can be interpreted on a reasonable scale e.g.: A |pcc| = 1 implies that U and W are highly correlated with a perfect linear correlation; 1 > |pcc| ≥ 0.8 implies a strong linear correlation; 0.8 > |pcc| ≥ 0.4 indicates a moderate linear correlation; 0.4 > |pcc| > 0 suggests a weak linear correlation, while a |pcc| = 0 means no linear dependency. It is worth pointing out that pcc measures the relationship of two variables, or how one changes responding to the changes of the other, not how each of them changes by itself.

I/*σ*I, Rmerg, Rmeas, Rpim, and CC1/2 are evaluated in different resolution shells at the end of data processing. Each of them can be treated as a variable with the values indexed by resolution shell, and a pcc between each pair (U, W) of these measures can be calculated by,

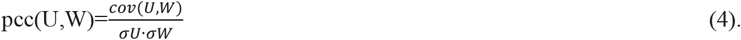

Here *cov(U,W)* is the covariance of U, W. *σU, σW* are the standard deviations of U and W respectively.

### Test and Results

#### 1. How does CC1/2 correlate to I/*σ*I?

To answer this question, we surveyed 815 data sets recently collected for a variety of projects by different research groups, with low-shell Rmerg better than 0.080 (to reduce potential disturbance from possible mis-indexing (Evans et al., 2013)), CC1/2 above 0.0 and resolution better than 6.0Å. All data sets were processed by XDS (Kabsch, 2010) and AIMLESS (Evans, 2011) (see Supplementary 1 for the detailed statistics). The plot of CC1/2 vs I/*σ*I in the highest-resolution-shell (Figure 1A) shows a wide distribution, which suggests that CC1/2 does not have a close correlation with I/*σ*I or signal-to-noise ratio. As different values of 0.1 to 0.4 have been suggested to select resolution cutoffs, we took a closer look of those 478 data sets with CC1/2 between 0.1 and 0.4. Figure 1B shows the breakdown of the number of these 478 data sets vs I/*σ*I. A majority (84%) of them have I/*σ*I below 1.1, and 69% have an I/*σ*I below 0.9.

**Figure 1A.**
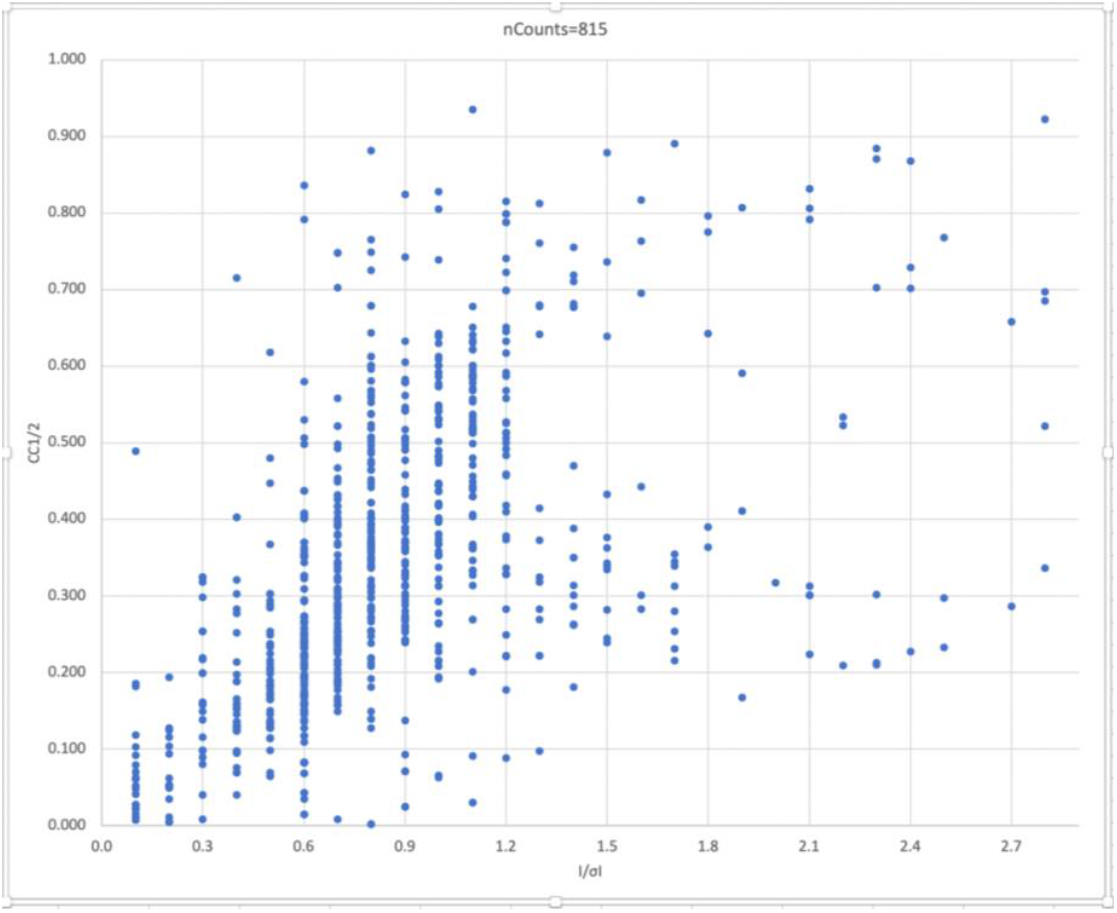
Plot of high-resolution-shell CC1/2 vs high-resolution-shell I/*σ*I of all the 815 data sets.

**Figure 1B.**
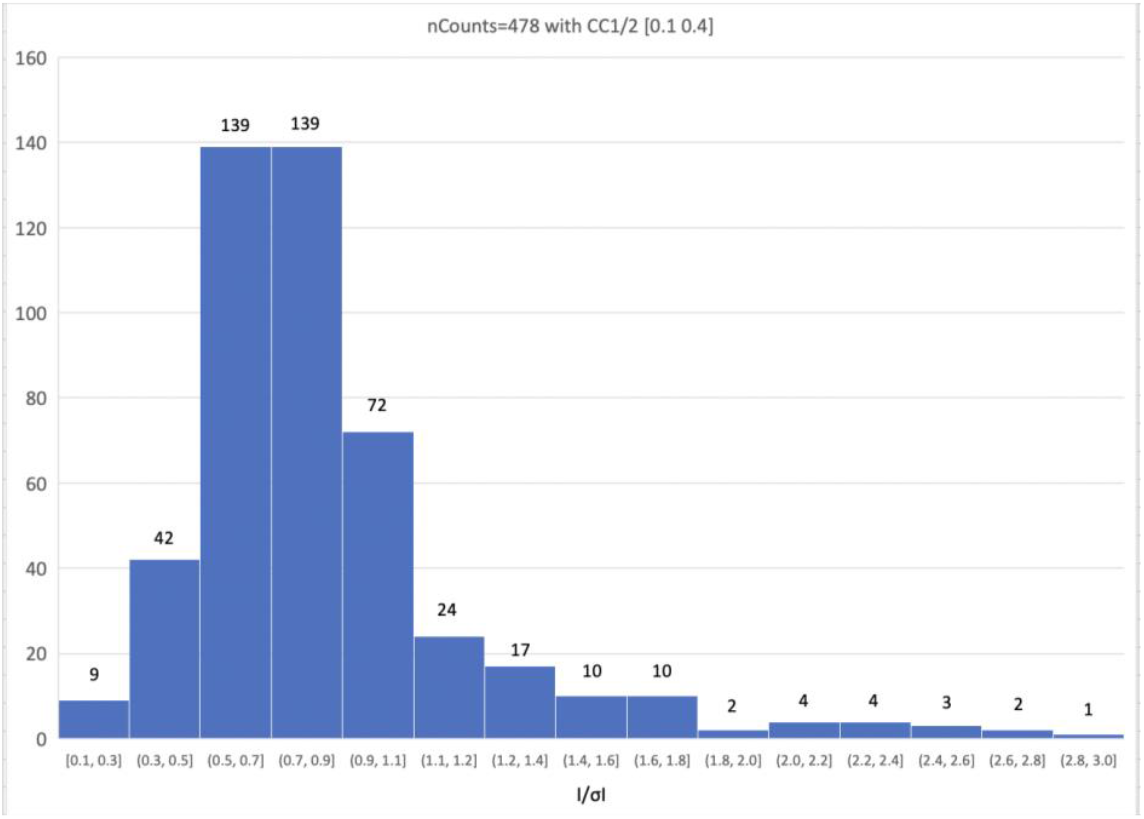
Breakdown of number with I/*σ*I out of the 478 data sets with high-resolution-shell CC1/2 in the range [0.1, 0.4].

The overall *Multi* covers a large range of [1.1, 32.7] with a median at 10.1 and an average of 10.4. The CC1/2’s low correlation to I/ *σ* I in the high-resolution shell seems intrinsic and is not attributable to *Multi*. For strong reflections, when *Multi* increases, the I/*σ*I could improve by a factor up to 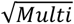 as merging cancels out the random errors while systematic errors are negligible. However, as the reflection intensity decreases, systematic errors will become more and more significant compared to the intensity, the magnitude of improvement will get less than 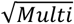. In the high-resolution shells, the *Multi* will also be significantly smaller than the overall as fewer reflections become measurable. Therefore, using the *Multi*, especially the overall *Multi*, to describe I/*σ*I’s dependency within the high-resolution shell is conceptually inappropriate. It’s also worth pointing out that these 815 data sets cover a wide range with various characteristics, 64 of them exhibiting noticeable anisotropy detected by AIMLESS with ANSdd bigger than 5%, and in one very severe case, with an ANSdd value of 37.4%. ANSdd denotes the max{2.0(r*i* – r*j*)/(r*i* + r*j*) | *i,j*=1,2,3; r*i* is the estimated resolution along d*i*} hereafter for simplicity.

CC1/2 had been predicted to be closely correlated with I/*σ*I (Karplus et al., 2015). This is quite contradictory to the experimental data shown above. The disagreement may reflect fact that the assumed model used in the prediction is too simple to represent the variety of real-world data.

#### 2. How are Rmerg, Rmeas, Rpim, CC1/2 correlated to each other?

As mentioned in the Methods section, Rmerg, Rmeas, Rpim and CC1/2 are all derived from comparing the intensities of equivalent reflections. To understand how these metrics are correlated, pcc values (see equation 4) among them were calculated using experimental data with various characteristics (see Table 1). The trypA data set was collected from one standard crystal of bovine trypsin (223 amino acid residues) of P212121 symmetry by one sweep of 120^°^. The trypB data set was collected from a separate bovine trypsin crystal with one 720° sweep but was processed into two different reduced sets using the first 30° and all 720° of data (denoted as trypB30 and trypB720 hereafter) respectively. Data6OPM, Data7MRQ, Data8D7K and Data9ATU are data for solving the structures deposited in PDB with the corresponding IDs. Data ANSdd125, ANSdd227, ANSdd313 and ANSdd374 are 4 of those 64 data sets (described in section 1) with anisotropic ANSdd of 12.5%, 22.7%, 31.3% and 37.4% respectively. MERG10 is merged from 10 data sets each from a different bovine trypsin crystal with a 20^°^ sweep. trypA, trypB, Data8D7K, ANSdt125,227,313,374 were all indexed and integrated with XDS and scaled by AIMLESS. Data6OPM were indexed and integrated with XDS and scaled with XSCALE (Kabsch, 2010). Data7MRQ and Data9ATU were processed with HKL2000 (Otwinowski et al., 1997). MERG10 were indexed and integrated with DIALS (Winter et al., 2018) and merged by XIA2.MULTIPLEX (Gildea et al., 2022). trypADLS is trypA processed by DIALS. Values of all the measures by resolution shell were harvested from data processing (see Supplementary 2 for details). trypB30 with *Compl* 50% and *Multi* of 2 is used as a test on the incomplete data. All other data are complete with *Compl* above 90% and *Multi* between 3 to 26.

**Table 1.** The pcc calculated using equation 4 among Rmerg, Rmeas, Rpim and CC1/2: MgMs=pcc(Rmerg,Rmeas), MgPm=pcc(Rmerg,Rpim), MsPm=pcc(Rmeas,Rpim), MgCc=pcc(Rmerg,CC1/2), MsCc=pcc(Rmeas,CC1/2), PmCc=pcc(Rpim,CC1/2).

| pcc | MgMs | MgPm | MsPm | MgCc | MsCc | PmCc |
| --- | --- | --- | --- | --- | --- | --- |
| trypA | 0.998 | 0.988 | 0.996 | -0.979 | -0.979 | -0.971 |
| trypADLS | 1.000 | 0.999 | 0.999 | -0.984 | -0.984 | -0.984 |
| trypB30 | 1.000 | 1.000 | 1.000 | -0.918 | -0.916 | -0.918 |
| trypB720 | 1.000 | 0.986 | 0.989 | -0.892 | -0.900 | -0.951 |
| Data6OPM | 1.000 | 1.000 | 1.000 | -0.959 | -0.959 | -0.958 |
| Data7MRQ | 1.000 | 0.994 | 0.996 | -0.959 | -0.964 | -0.978 |
| Data8D7K | 1.000 | 1.000 | 1.000 | -0.984 | -0.984 | -0.984 |
| Data9ATU | 1.000 | 0.989 | 0.992 | -0.956 | -0.960 | -0.972 |
| ANSdt125 | 0.999 | 0.995 | 0.998 | -0.939 | -0.950 | -0.967 |
| ANSdt227 | 1.000 | 0.999 | 1.000 | -0.967 | -0.971 | -0.976 |
| ANSdt313 | 1.000 | 1.000 | 1.000 | -0.992 | -0.992 | -0.992 |
| ANSdt374 | 1.000 | 1.000 | 1.000 | -0.900 | -0.902 | -0.903 |
| MERG10 | 0.999 | 0.972 | 0.982 | -0.914 | -0.904 | -0.982 |

Table 1 shows that pcc among Rmerg, Rmeas and Rpim are almost 1.0, which is expected as they are linearly related by defined formulas. Surprisingly, |pcc| between CC1/2 and Rmerg, Rmeas or Rpim are also very high approaching the perfect value of 1.0. This suggests that CC1/2, Rmerg, Rmeas and Rpim are not only strongly correlated but also with a near perfect linear correlation, although CC1/2 takes a different definition. The negative sign only means CC1/2 decreases while the other three increase. Considering the variety of the data and different processing programs used, the high near perfect linear correlations among {Rmerg, Rmeas, Rpim, CC1/2} is intrinsic, regardless of data sets or data processing programs.

#### 3. Correlations of Rmerg, Rmeas, Rpim, CC1/2, and I/*σ*I with Resolution

Each of Rmerg, Rmeas, Rpim, CC1/2 and I/*σ*I has been used to select resolution cutoffs. To compare how these indicators track resolution changes, their pcc with resolution were calculated for the 13 data sets listed in Table 1, and the results are summarized in Table 2. From Table 2, all these pcc can be different for different data sets. But for the same data set, the pcc of Rmerg, Rmeas and Rpim with resolution are very close or almost the same. The pcc of CC1/2 are either close to or lower than those of Rmerg, Rmeas and Rpim for 12 out of the 13 data sets tested and only larger but still comparable for data ANSdd374 which shows severe anisotropy. The results demonstrated that CC1/2 is even less sensitive than Rmerg, Rmeas and Rpim in terms of tracking resolution changes or as an indicator for selecting resolution cutoffs, which agrees well with the fact that CC1/2 does not have a single value as the threshold generally applicable for different cases, similar to Rmerg, Rmeas and Rpim.

**Table 2.** The pcc calculated using equation 4 between Rmerg, Rmeas, Rpim, CC1/2, I/*σ*I and resolution.

| pcc | Rmerg | Rmeas | Rpim | CC1/2 | $I/\sigma I$ |
| --- | --- | --- | --- | --- | --- |
| trypA | -0.552 | -0.535 | -0.508 | 0.535 | 0.876 |
| trypADLS | -0.842 | -0.841 | -0.838 | 0.767 | 0.997 |
| trypB30 | -0.605 | -0.601 | -0.605 | 0.482 | 0.824 |
| trypB720 | -0.560 | -0.553 | -0.503 | 0.367 | 0.774 |
| Data6OPM | -0.469 | -0.469 | -0.466 | 0.349 | 0.888 |
| Data7MRQ | -0.686 | -0.678 | -0.643 | 0.541 | 0.879 |
| Data8D7K | -0.467 | -0.468 | -0.470 | 0.510 | 0.853 |
| Data9ATU | -0.771 | -0.765 | -0.718 | 0.626 | 0.904 |
| ANSdd125 | -0.455 | -0.447 | -0.431 | 0.318 | 0.761 |
| ANSdd227 | -0.541 | -0.548 | -0.559 | 0.569 | 0.800 |
| ANSdd313 | -0.486 | -0.486 | -0.485 | 0.439 | 0.773 |
| ANSdd374 | -0.559 | -0.560 | -0.561 | 0.677 | 0.865 |
| MERG10 | -0.868 | -0.857 | -0.796 | 0.710 | 0.986 |

More interestingly, the pcc between I/*σ*I and resolution are significantly greater than those of Rmerg, Rmeas, Rpim and CC1/2 across all 13 data sets, suggesting that I/*σ*I has a strong linear correlation (pcc close to or well above 0.8) with resolution, and thus can track the resolution changes more closely. Therefore, I/*σ*I is much more sensitive as an indicator to select resolution cutoffs than other metrics. This can be theoretically explained by the Debye-Waller theory (Debye, 1913; Waller, 1923). Reflection intensity decreases as the diffraction angle increases due to attenuation by atomic dynamics. Defects of the crystal could also contribute to observed attenuation. In other words, intensity I and I/*σ*I will decrease with resolution, which provides another theoretical base for using I/*σ*I to judge data significance in addition to its defined capacity as a signal-to-noise ratio.

#### 4. What do Rmerg, Rmeas, Rpim, CC1/2, and I/*σ*I really measure? How to use them?

Both the high near-perfect linear correlations among {Rmerg, Rmeas, Rpim, CC1/2} and their similar but low correlations with resolution pointed to the fact that these four indicators are all evaluated from comparing intensities of equivalent reflections (see Methods) and theoretically what they measure is the same, namely the equivalency or internal consistency among reflections with Laue-symmetry as the underlying fundamental. In addition, this functional similarity agrees well with the fact that neither of {Rmerg, Rmeas, Rpim, CC1/2} has a single value as the threshold generally applicable for different cases in selecting resolution cutoffs or assessing data quality. Furthermore, CC1/2 does not have close correlation with I/ *σ* I (signal-to-noise ratio) as demonstrated by experimental data with various characteristics. Therefore, it is highly debatable to differentiate CC1/2 from {Rmerg, Rmeas, Rpim} especially by claiming that CC1/2 is particularly capable to assess the precision or quality of the reduced data (Karplus et al., 2015; Diederichs, 2015). In a contemporary diffraction experiment with an area detector, a majority of reflections are symmetry-related, and thus {Rmerg (as its previous name Rsym suggested), Rmeas, Rpim, CC1/2} in the low-resolution-shell can serve as a reliable indicator for the correctness (or trueness) of Laue-symmetry.

I/*σ*I, defined as a signal-to-noise ratio, is the indicator measuring the signal strength compared with estimated error. Compared with {Rmerg, Rmeas, Rpim, CC1/2}, I/*σ*I is arguably the best measure for assessing data quality in terms of signal-to-noise ratio. In addition, it has strong linear correlation with resolution, with both the Debye-Waller theory and Laue-symmetry as the underlying fundamentals. Therefore, the high-shell I/*σ*I is more capable and sensitive for selecting resolution cutoffs, or in other words the extent to which the data is still reliable. Furthermore, the overall I/*σ*I is meaningful and capable to measure the overall strength of a data set, while neither that of {Rmerg, Rmeas, Rpim, CC1/2} has the capacity to do so, which can be very helpful to compare different data sets.

#### 5. Resolution extension beyond I/*σ*I of 2.0

The research papers on CC1/2 (Karplus et al., 2012; Karplus et al., 2015) had a great contribution to the field by demonstrating that resolution cutoffs traditionally selected by {Rmerg, Rmeas, Rpim} may be conservative for some cases when weak and noisy data in extended higher resolutions were excluded. Nevertheless, similar to {Rmerg, Rmeas, Rpim}, CC1/2 does not have a single value as the threshold generally applicable to different cases. Therefore, the resolutions selected by some to somewhat arbitrary CC1/2 values need to be validated through further analysis. The paired-refinement procedure was proposed for the purpose (Karplus et al., 2015; Maly, et al., 2020), which compares the R/Rfree/Rgap (Rgap is the difference of Rfree and R) at the lower (reference) resolution instead of the actual target resolution extended.

To understand the model changes by including weak and noisy data beyond the resolution selected by I/*σ*I=2.0, we studied three cases as examples. The first one is bovine trypsin. Data trypA (processed to 1.14Å, described in section 2) of resolution 1.53Å, where {I/*σ*I=2.0, CC1/2=0.786}, was used to solve the structure by molecular replacement with PHASER (McCoy et al., 2007) inside PHENIX package (Liebschner et al., 2019). The model was completed with COOT (Emsley et al., 2010) and refined with PHENIX.REFINE (Afonine et al., 2012). The second is the NE/Eap4/NE complex with 6 chains of two different proteins, whose structure was originally solved at 2.05Å (PDB ID 9ATU) (Mishra et al., 2024). The third is a fragment of chicken CNTN4 that includes four point mutations in its binding site for amyloid precursor protein originally solved at 3.20Å (PDB ID 7MRQ) (Karuppan et al., 2021). Step-by-step paired-refinements following the published protocols (Karplus et al., 2015; Maly, et al., 2020; Winter et al., 2018) were carried out to extend to higher resolutions (see Table 3). PHENIX.REFINE was used for each refinement. For the trypsin and NE/Eap4/NE cases with the resolutions to be checked all within the range where auto-tracing works well, we slightly modified the protocol by auto-tracing the refined electron density maps with PHENIX.AUTOBUILD (Terwilliger et al., 2008) and compared the resulting R/Rfree/Rgap at the lower (reference) resolution, which would add a quantitative evaluation of the qualities of electron density maps.

**Table 3.** Paired-Refinements to test extended resolutions. (A). Tryspin; (B). NE/Eap4/NE; (C). CNTN4. The data used are ‘trypA’, ‘Data9ATU’, and ‘Data7MRQ’ respectively (see section 2 and Supplementary 2 for details of reduction statistics). Here, R.I.C. is the target with {Resolution, in-shell I/*σ*I, in-shell CC1/2} to check, which will also serve as the reference of next step; R.L.={R, Rfree at lower or reference resolution}, R.H.={R, Rfree at extended higher or target resolution to check}. The test is considered true if two of {R, Rfree, Rga} are better or Rfree is close and Rgap is better.

| Trial | R.I.C. | R.L. | R.H. | Test |
| --- | --- | --- | --- | --- |
| start | {1.53 2.0 0.786} | {----- -----} | {----- -----} | ---- |
| step1 | {1.44 1.3 0.722} | {0.1984 0.2234} | {0.2020 0.2273} | true |
| step2 | {1.40 1.1 0.657} | {0.2051 0.2282} | {0.2038 0.2262} | true |
| step3 | {1.34 0.8 0.591} | {0.2061 0.2241} | {0.2036 0.2211} | true |
| step4 | {1.25 0.5 0.464} | {0.2157 0.2311} | {0.2285 0.2448} | false |

**Table 3(B).**
| Trial | R.I.C. | R.L. | R.H. | Test |
| --- | --- | --- | --- | --- |
| start | {2.10 2.2 0.757} | {----- -----} | {----- -----} | ---- |
| step1 | {2.03 1.4 0.476} | {0.2205 0.2545} | {0.2167 0.2488} | true |
| step2 | {1.95 0.7 0.275} | {0.2210 0.2596} | {0.2245 0.2680} | false |

**Table 3(C).**
| Trial | R.I.C. | R.L. | R.H. | Test |
| --- | --- | --- | --- | --- |
| start | {3.18 2.0 0.845} | {----- -----} | {----- -----} | ---- |
| step1 | {3.10 1.5 0.853} | {0.2026 0.2309} | {0.2052 0.2310} | true |
| step2 | {3.04 1.2 0.702} | {0.1994 0.2308} | {0.2071 0.2226} | true |
| step3 | {2.98 0.9 0.491} | {0.1985 0.2249} | {0.2022 0.2272} | true |
| step4 | {2.92 0.7 0.304} | {0.2014 0.2396} | {0.2021 0.2447} | false |

For trypsin data, 1.53Å is selected by a traditional I/*σ*I=2.0 cutoff where {I/*σ*I=2.0, CC1/2=0.786} to solve the structure. According to the paired-refinements, the resolution could be extended to 1.34Å (a gain of 0.19Å). Wilson plot in Figure 2(A) starts to deviate around 1.4Å and clearly gets abnormal or suspicious beyond resolution of 1.3Å. From curiosity on how noisy or weak data would affect the electron density map, we refined the model using all the data up to 1.14 Å at which CC1/2=0.202 and I/*σ*I=0.1, followed by a PHENIX.AUTOBUILD. If the contribution from the noisy data in the high-resolution shell is positive, a better electron density map and thus a better traced model would be expected. But the result shows otherwise. Figure 2(B) is part of the electron density maps around SER119 superposed with the traced models from using resolutions 1.14Å (left) and 1.40Å (Right), respectively. The electron density map from 1.14Å cutoff looks significantly noisier. For example, its auto-tracing failed building GLY120 after SER119 due to too much erroneous extra density, leading to one of 2 main-chain gaps. The erroneous density also affected the building of side chains (such as a TYR was built instead of the correct TRP121).

**Figure 2.**
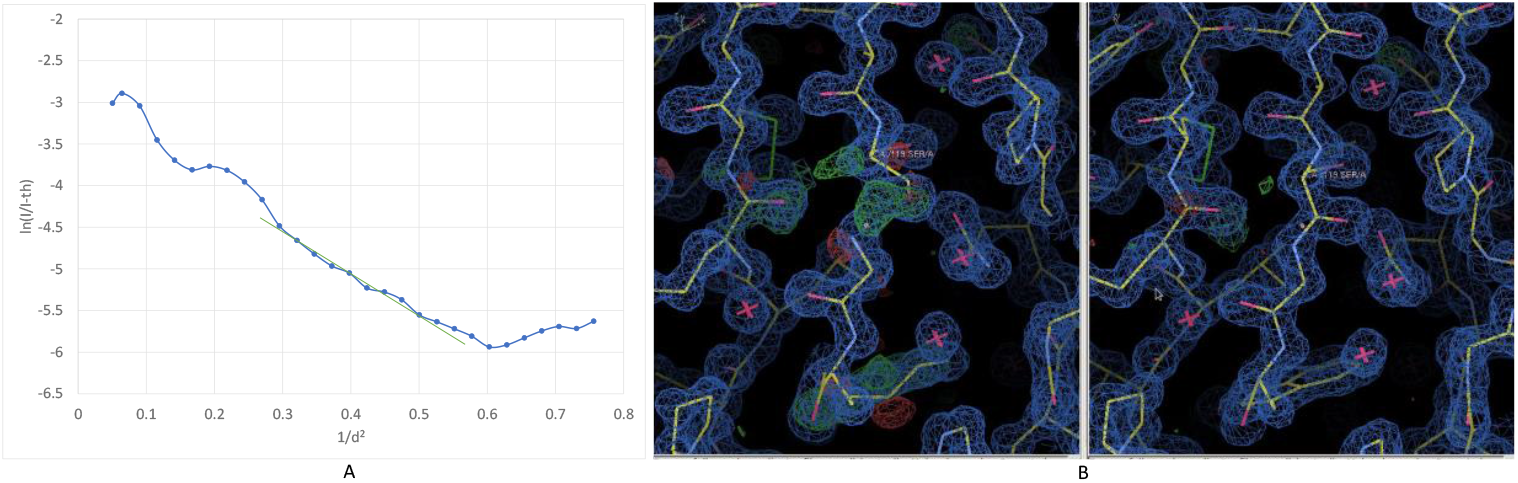
(A) Wilson plot calculated by WILSON in CCP4 (Winn et al., 2011) of trypA, where d is resolution. (B) part of the electron density maps around SER119 superposed with the traced models from PHENIX.AUTOBUILD for resolution cutoff of 1.14Å (Left) and 1.40Å (Right), respectively, displayed by COOT defaults.

For the NE/Eap4/NE data, we took the deposited structure in the PDB and refined at 2.10Å where {I/ *σ* I=2.2, CC1/2=0.757} as the starting model. According to the paired-refinements, the resolution could be extended to 2.03Å (a gain of 0.07Å). For the CNTN4 data, the PDB deposition was first refined at 3.18Å where {I/*σ*I=2.0, CC1/2=0.845} and then served as the starting model. The resolution could be extended to 2.98Å based on the paired-refinements (a gain of 0.20Å). Figure 3 shows that the Wilson plots start to deviate around 2.1Å and 3.0Å for NE/Eap4/NE and CNTN4 data, respectively.

**Figure 3.**
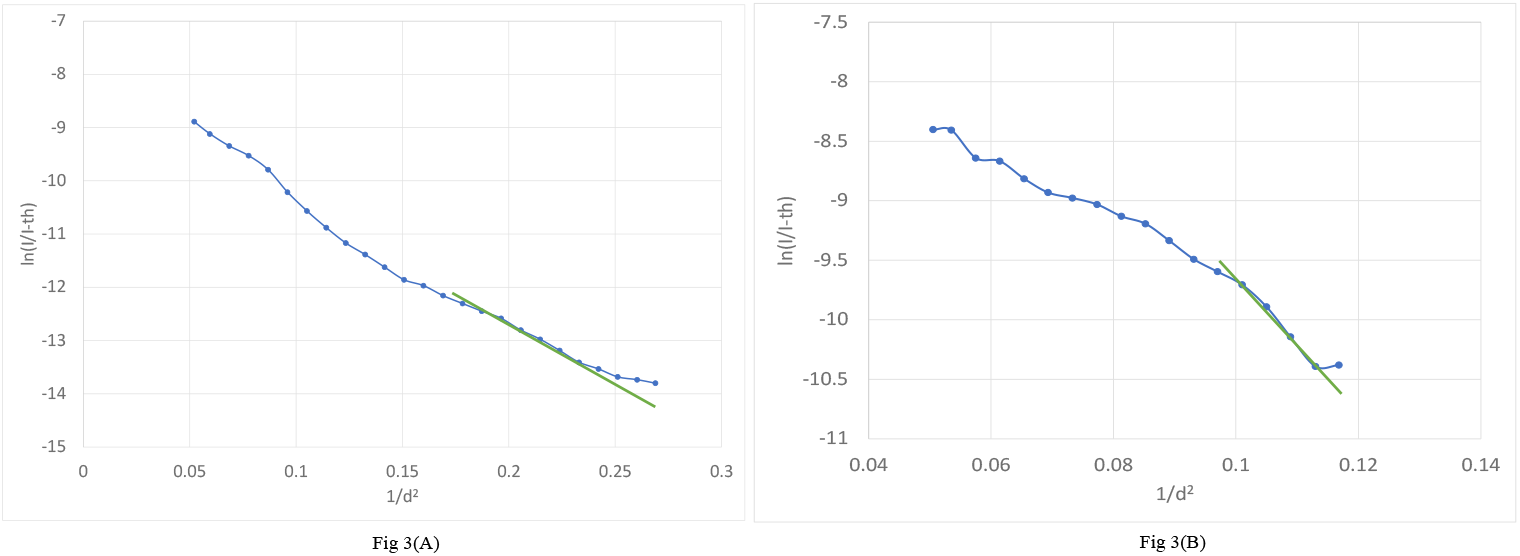
Wilson plot calculated with WILSON in CCP4. (A) NE/Eap4/NE data. (B) CNTN4 data.

It’s worth noticing that, since I/*σ*I and CC1/2 drop suddenly and quickly at high-resolution shells beyond the resolution cutoffs selected by a I/*σ*I of 2.0, the extensions are typically very limited (Evans et al., 2013; Karplus et al., 2015; Maly, et al., 2020; Winter et al., 2018). The paired-refinements demonstrated that the weak and noisy reflections in the extended higher resolution shell may contribute as a restrain to the refinement, leading to a lower R/Rfree at the lower (reference) resolution.

However, since electron density (as a Fourier transform) depends on the reflections collectively, all the reflections to the target resolution must be included to evaluate the quality of the refined structure, including but not limited to R/Rfree, overall quality of the electron density and model geometry. Table 4 lists statistics from the refinements using all reflections to the different resolutions selected by paired-refinements, which showed that adding the noisy and weak data in the extended higher resolution shell didn’t change the geometry of the model or did so in a very limited fashion. On the other hand, both R/Rfree and B-factors (the average representing the overall and the minimum indicating the bottom-line) all became noticeably worse, which would suggest an overall deterioration or no improvement of electron density quality. Therefore, a better R/Rfree to the lower (reference) resolution may not be a proof of better quality of the refined structure at the higher resolution extended, and thus not a strict or reliable validation of the higher resolution extended.

**Table 4.** Final refinements using all data to the extended resolutions selected by paired-refinements. (A). Trypsin; (B). NE/Eap4/NE; (C). CNTN4. R/Rfree is the overall from final refinements. <B>(min) is the average and the minimum B-factors. RMSD is from the pair-wise 3D-structure alignment of the models using USALIGN (Zhang et al., 2022).

| Reso | R/Rfree |  | <B> (min) | RMSD{models (A, B) } |  |  |
| --- | --- | --- | --- | --- | --- | --- |
| 1.53 | 0.1766 | 0.1953 | 11.8 (5.8) | ---- |  |  |
| 1.44 | 0.1871 | 0.2098 | 12.0 (6.1) | 0.02 | ---- |  |
| 1.40 | 0.1901 | 0.2031 | 13.0 (6.7) | 0.02 | 0.02 | ---- |
| 1.34 | 0.1963 | 0.2158 | 13.1 (7.4) | 0.02 | 0.02 | 0.01 |

**Table 4(B).**
| Reso | R/Rfree |  | <B> (min) | RMSD{models (A, B) } |
| --- | --- | --- | --- | --- |
| 2.10 | 0.1802 | 0.2148 | 58.9 (30.4) | ---- |
| 2.03 | 0.1833 | 0.2235 | 60.8 (32.3) | 0.03 |

**Table 4(C).**
| Reso | R/Rfree |  | <B> (min) | RMSD{models (A, B) } |  |  |
| --- | --- | --- | --- | --- | --- | --- |
| 3.18 | 0.2041 | 0.2306 | 86.6 (51.6) | ---- |  |  |
| 3.10 | 0.2065 | 0.2313 | 93.3 (54.5) | 0.06 | ---- |  |
| 3.04 | 0.2079 | 0.2335 | 95.5 (55.8) | 0.07 | 0.07 | ---- |
| 2.98 | 0.2183 | 0.2409 | 89.3 (57.6) | 0.06 | 0.06 | 0.06 |

#### 6. Be careful of both geometry and B-factor changes when trying to extend resolution cutoffs

The most critical factors that determine structure quality include resolution, quality of the electron density map, model geometry, R/Rfree, and B-factors. A valid resolution cutoff extension should be, at least: (1) the model fits better into the electron density that shows structural features matching the resolution; (2) the electron density map with fewer or no change of disordered regions and gets cleaner with less erroneous extra density; (3). better or no change in Ramachandran plot and Clashscore (as modern refinement programs usually do not allow large deviations of bond lengths and angles from ideal values, Ramachandran plot and Clashscore may not show much difference during paired-refinements); (4). better or no significant increase in R/Rfree; (5). smaller or no significant increase of B-factors which may not be a proof of no deterioration of electron density quality, but a dramatic increase could be a sign of deterioration.

As shown in equation 3, changing B-factors have a large impact on the electron density. It is well known that after the molecules are correctly built, B-factors become the most efficient means of reducing R/Rfree during the refinement. However, aggressively extending the resolution to a higher cutoff would likely make the quality worse if the geometry doesn’t change much but B-factors are pushed significantly higher to account for the errors. Indeed, out of the latest 100,000 structures in PDB solved by X-ray diffraction, the average B-factor of macro molecules in the 14,283 structures with reported high-shell I/*σ*I between 1.0 and 2.0 is 49.9, while the average B-factor of all the 4,326 structures with reported high-shell I/*σ*I below 1.0 is significantly worse at 59.4.

#### 7. Neither {Rmerg, Rmeas, Rpim, CC1/2} nor I/*σ*I is Error-Model Free

As pointed out in the Methods, error corrections are critical and applied to evaluate both I(*i*) and *σ*I(*i*) of the reduced data typically through error models during data processing. Thus all {Rmerg, Rmeas, Rpim, CC1/2, I/*σ*I} calculated from {I(*i*), *σ*I(*i*)} are affected by the actual error models used in a data processing program. Different programs may use different ways to model errors, leading to different values of these metrics for the same data. The validity of the error models used by a data processing program is justified by the reduced data {**H**(*i*), I(*i*), *σ*I(*i*)}} capable in solving structures. Therefore, neither {Rmerg, Rmeas, Rpim, CC1/2} nor I/*σ*I is error-model free. It is theoretically unfounded and highly debatable to single out CC1/2 as model-free which was the major argument to favor CC1/2 over I/*σ*I (Karplus et al. 2012, Karplus et al. 2015). Furthermore, the fundamental difference between CC1/2 and I/*σ*I is that CC1/2 measures internal consistence in terms of Laue symmetry while I/*σ*I measures the signal-to-noise ratio by definitions, which represent different aspects of data quality. As an example, we processed the trypA with different programs (see Table 5). Table 5 didn’t show more differences in I/*σ*I than in CC1/2 from different data processing programs except DIALS that came into use recently compared to others. It’s not clear why the I/*σ*I from DIALS is significantly different from others (not sure, seems more likely due to some formulation than errors model itself). <I/*σ*I> and <I>/<*σ*I> can be different if I(*i*)/*σ*I(*i*) are significantly different among the reflections summed within a resolution shell. HKL2000 does not list <I/*σ*I> in its output log file, but instead <I>, <*σ*I> separately, which may be changed in its future release.

**Table 5.** High-resolution-shell I/*σ*I, and CC1/2 from trypA processed by different data processing programs. IMOSFLM (Leslie et al., 2007) in the CCP4 package automatically cut the resolution to 1.28Å. To compare apples to apples, we applied the same resolution cutoffs for DIALS, XDS (XSCALE scaling), and AIMLESS (XDS integration + AIMLESS scaling). HKL2000 didn’t integrate the data beyond 1.54Å, at which the high-resolution-shell <I>, <*σ*I> were evaluated to be 0.2, 0.3 respectively, leading to <I>/<*σ*I>=0.67.

| Program | Resolution | I/σI | CC1/2 |
| --- | --- | --- | --- |
| IMOSFLM | 1.28 | 0.7 | 0.304 |
| XDS/XSCALE | 1.28 | 0.6 | 0.526 |
| XDS/AIMLESS | 1.28 | 0.5 | 0.464 |
| DIALS | 1.28 | 1.3 | 0.559 |
| HKL2000 | 1.54 | 0.7 | 0.566 |

#### 8. Summary

Based on theoretical and pcc analysis, {Rmerg, Rmeas, Rpim, CC1/2} all measure the equivalence of reflections with Laue-symmetry as the underlying fundamental, and the low-shell values of these metrics are reliable indicators for correctness (or trueness) of symmetry. Once symmetry correctness is determined, high-shell I/*σ*I may arguably be the best indicator to select resolution cutoffs for the following reasons: 1) I/*σ*I is the indicator that measures the signal strength compared with error levels in the data; 2) CC1/2 does not have close correlation with the signal-to-noise ratio, thus may not be capable to make reliable judgement on the extent to which the data are still reliable; 3) I/*σ*I has significantly higher correlation with resolution thus better sensitivity as an indicator; 4) The overall I/*σ*I provides a measure of overall strength of the data, while neither that of {Rmerg, Rmeas, Rpim, CC1/2} does. Hence, it is highly debatable and could be misleading to single out CC1/2 as particular useful for detecting weak signals in high resolution shells (Karplus et al. 2012, Karplus et al. 2015).

Extended resolution cutoffs selected by CC1/2 need further verification because CC1/2 does not have a single value as the threshold generally applicable for different cases. The paired-refinement protocol provides a way for this purpose (Karplus et al., 2015; Maly, et al., 2020). However, a better R/Rfree to the lower (reference) resolution is not a proof of better quality of the refined structure at the extended higher or target resolution. Since electron density (as a Fourier transform) depends on the reflections collectively, all the reflections to the extended higher or target resolution cutoff must be included to evaluate the final structure quality and R/Rfree. Comparing the statistics from refinements using all reflections at different resolutions plus checking the quality of electron density map is a strict and traditional way for the more comprehensive verification, which is commonly used to test resolution cutoffs at the last stage of the structure solution process.

Structures solved from data with traditional resolution cutoffs selected by I/*σ*I of 2.0 have been demonstrated to be reliable and can serve as the starting models when resolution extensions are to be tried. Otherwise, a resolution cutoff corresponding to a lower I/*σ*I of ∼1.5 may represent a better starting point to save time from trial-and-errors during the whole process of structural solution.

## Discussion

The quality of the final structural model from X-ray crystallography relies on the quality of the data selected at the data acquisition step. In principle, this selection could be delayed to the stage of structural refinement for a final evaluation by electron density map quality, geometry, R/Rfree, B-factors, etc. However, practically speaking, structure solution still requires significant time to get to the final stage, while data acquisition is comparatively rapid. In most cases, to get the best reduced data, you may need data quality evaluation to make changes or adjust parameters at data collection, especially transmission and more importantly detector distance that is critical for optimizing the spot-separations (spatial resolution) while maximizing the samples’ diffraction capability, which in turn will affect the background and errors estimation in data processing and thus the quality of the reduced data. {Rmerg, Rmeas, Rpim, CC1/2} have been used to measure data quality and select resolution cutoffs, but none of them seems as sensitive, meaningful, or accurate as I/*σ*I.

Nowadays, most structures are solved by using existing structures as templates through molecular replacements or docking into Cryo-EM maps. In the long run, it will be beneficial for X-ray crystallography to keep producing more high-quality structures by holding up its data selection standards. Furthermore, prediction accuracy will be improved when more high-quality structures are available for AI-based modeling programs such as AlphaFold (Jumper et al., 2021; Abramson et al., 2024), RoseTTAfold (Baek et al., 2021) etc. to train their models. It is worth pointing out that crystal diffraction methods refine the B-factor in addition to the position for each atom in the structure model. High-quality structures will have lower errors in B-factors, and thus, will be more suitable for studying molecular dynamics (Ringe et al., 1986; Carugo et al., 1997; Winn et al., 2001; Sun et al., 2019; Pearce et al., 2021) or for quantifying and detecting radiation damage (Kathryn et al., 2022). That said, it would be beneficial for journals and grant reviewers to add some credit to researchers that report structures already solved but repeated with significantly higher quality. It would also be beneficial to include the key data selection standard in publications and PDB depositions, which would provide a tracking record for quick reference in the future.

## Acknowledgement

This research is funded and supported by the Southeast Regional Collaborative Access Team (SER-CAT) Member Institutions (a list may be found on the SER-CAT website), the University of Georgia Research Foundation, and the Georgia Research Alliance, and is also supported in part by grant R35GM140852 to BVG and by award number R15NS108371 to SB. We thank Dr. Yifan Cheng of HHMI & University of California San Francisco for discussion on the current developments of Cryo-EM. We also thank Dr. Stephen K. Burley, Director of the RCSB Protein Data Bank for critically reviewing the manuscript and providing feedback. X-ray diffraction data were collected at SER-CAT beamline 22-ID at APS (Advanced Photon Source), Argonne National Laboratory and at NYX beamline 19-ID at NSLS-II (National Synchrotron Light Source II), Brookhaven National Laboratory. We thank Dr. Kevin Battaile, Dr. Unmesh Chinte and Dr. Zhongmin Jin for helping SER-CAT using NYX beamline 19-ID. Use of APS and NSLS-II was supported by the U.S. Department of Energy, Office of Science, Office of Basic Energy Sciences.

## Author Contributions

ZQF conceived the project and performed the analysis. ZQF and BVG prepared the draft. BVG, SB and FD assisted in the analysis. JC organized the beamlines for data collection. PK checked and formatted the References. PK, DJM, BCW helped in works related to structural biology. All coauthors participated in critical reading of the paper.

## Data Availability Statement

All data generated or analyzed during the study are included in this published article and its supplementary information files.

## References

Abramson, J., Adler, J., Dunger, J., Evans, R., Green, T., Pritzel, A., Ronneberger, O., Willmore, L., Andrew J., Ballard, A.J., Bambrick, J., Bodenstein, S.W., Evans, D. A., Hung, C., O’Neill, M., Reiman, D., Tunyasuvunakool, K., Wu, Z., Žemgulytė, A., Arvaniti, E., Beattie, C., Bertolli, O., Bridgland, A., Cherepanov, A., Congreve, M., Cowen-Rivers, A.I., Cowie, A., Figurnov, M., Fuchs, F.B., Gladman, H., Jain, R., Khan, Y.A., Low, C.M.R., Perlin, K., Potapenko, A., Savy, P., Singh, S., Stecula, A., Thillaisundaram, A., Tong, C., Yakneen, S., Zhong, E.D., Zielinski, M., Žídek, A., Bapst, V., Kohli, P., Jaderberg, M., Hassabis, D. & Jumper, J.M. (2024). Nature. 630, 493–500.

Afonine, P.V., Grosse-Kunstleve, R.W., Echols, N., Headd, J.J., Moriarty, N.W., Mustyakimov, M., Terwilliger, T.C., Urzhumtsev, A., Zwart, P.H. & Adams, P.D. (2012). Acta Cryst. D68, 352–367.

Agirre, J., Atanasova, M., Bagdonas, H., Ballard, C.B., Baslé, A., Beilsten-Edmands, J., Borges, R.F., Brown, D.G., Burgos-Marrmol, J.J., Berrisford, J.M., Bond, P.S., Caballero, I., Catapano, L., Chojnowski, G., Cook, A.G., Cowtan, K.D., Croll, T.I., Debreczeni, J.E., Devenish, N.E., Dodson, E.J, Drevon, T.R., Emsley, P., Evans, G., Evans, P.R., Fando, M., Foadi, J., Fuentes-Montero, L., Garman, E.F., Gerstel, M., Gildea, R.J. Hatti, K., Hekkelman, M.L., Heuser, P.,Hoh, S.W., Hough, M.A., Jenkins, H.T., Jiménez, E., Joosten, R.P., Keegan, R.M., Keep, N., Krissinel, E.B., Kolenko, P., Kovalevskiy, O., Lamzin, V.S., Lawson, D.M., Lebedev, A.A., Leslie, A.G.W., Lohkamp, B., Long, F., Maly, M., McCoy, A.J., McNicholas, S.J., Medina, A., Millán, C., Murray, J.W., Murshudov, G.N., Nicholls, R.A.,Noble, M.E.M., Oeffner, R., Pannu, N.S., Parkhurst, J.M., Pearce, N., Pereira, J., Perrakis, A., Powell, H.R., Read, R.J., Rigden, D.J., Rochira, W., Sammito, M., Rodríguez, F.S., Sheldrick, G.M., Shelley, K.L., Simkovic, F., Simpkin, A.J., Skubak, P., Sobolev, E., Steiner, R.A., Stevenson, K., Tews, I., Thomas, J.M.H., Thorn, A., Valls, J.T., Uski, V., Usón, I., Vagin, A., Velankar, S., Vollmar, M., Walden, H., Waterman, D., Wilson, K.S., Winn, M.D., Winter, G.,Wojdyr, M. & Yamashita, K. (2023). Acta Cryst. D79, 449–461.

Baek, M., DiMaio, F., Anishchenko, I., Dauparas, J., Ovchinnikov, S., Lee, G.R., Wang, J., Cong, Q., Kinch, L.N., Schaeffer, D., Millán, C., Park, H., Adams, C., Glassman, C.R., DeGiovanni, A., Pereira, J.H., Rodrigues, A.V., Van Dijk, A.A., Ebrecht, A.C., Opperman, D.J., Sagmeister, T., Buhlheller, C., Pavkov-Keller, T., Rathinaswamy, M.K., Dalwadi, U., Yip, C.K., Burke, J.E., Garcia, K.C., Grishin, N.V., Adams, P.D., Read, R.J. & Baker, D. (2021). Science. 373, 871–876.

Berman, H.M., Westbrook, J., Feng, Z., Gilliland, G., Bhat, T.N., Weissig, H., Shindyalov, I.N. & Bourne, P.E. (2000). Nucleic Acids Res. 28, 235–242.

Berman, H.M., Henrick, K. & Nakamura, H. (2003). Nature Structural & Molecular Biology 10, 980. 10.1038/nsb1203-980.

Boslaugh, S. (2012). Statistics in a Nutshell, 2nd Edition. O’Reilly Media, Inc.

Brünger, A.T. (1992). Nature. 355, 472–475.

Burley, S.K., Bhikadiya, C., Bi C., Bittrich, S., Chao, H., Chen, L., Craig, P.A., Crichlow, G.V., Dalenberg, K., Duarte, J.M., Dutta, S., Fayazi, M., Feng, Z., Flatt, J.W., Ganesan, S., Ghosh, S., Goodsell, D.S., Green, R.K., Guranovic, V., Henry, J., Hudson, B.P., Khokhriakov, I., Lawson, C.L., Liang, Y., Lowe, R., Peisach, E., Persikova, I., Piehl, D.W., Rose, Y., Sali, A., Segura, A., Sekharan, M., Shao, C., Vallat, B., Voigt, M., Webb, B., Westbrook, J.D., Whetstone, S., Young, J.Y., Zalevsky, A. & Zardecki, C. (2023). Nucleic Acids Res. 51, D488–D508.

Carugo, O. & Argos, P. (1997). Protein Eng. Des. Sel. 10, 777–787.

Cheng, Y., Grigorieff, N., Penczek, P.A. & Walz, T. (2015). Cell. 161, 438–449.

Debye, P. (1913). Annalen der Physik. 348, 49–92.

Diederichs, K. & Karplus, P.A. (1997). Nat. Struct. Mol. Biol. 4, 269–275.

Diederichs, K. (2016). Nucleic Acids Crystallography: Methods and Protocols. 1320. Edited by Eric Ennifar, New York: Springer.

Dubochet, J. & Knapek, E. (2018). PLoS Biol. 16, e2005550.

Emsley, P., Lohkamp,b Scott, W. & Cowtan, K. (2010). Acta Cryst. D66, 486–501.

Evans, P.R. (2011). Acta Cryst. D67, 282–292.

Evans, P.R. and Murshudov, G.N. (2013). Acta Cryst. D69, 1204–1214.

Frank, J. (2009). Q Rev Biophys. 42(3), 139–158.

Fu, Z.-Q. (2005). Acta Cryst. D61, 1643–1648.

Fu, Z.Q.: Kylin a program for crystal diffraction data reduction and data collection monitoring. (To be published).

Giacovazzo, C., Monaco, H.L., Artioli, G., Viterbo, D., Milanesio, M., Gilli, G., Gilli, P., Zanotti, G., Ferraris, G. & Catti, M. (1992). Fundamentals of Crystallography. Oxford University.

Gildea, R.J., Beilsten-Edmands, J., Axford, D., Horrell, S., Aller, P., Sandy, J., Sanchez-Weatherby, J., Owen, D., Lukacik, P., Strain-Damerell, C., Owen, R.L., Walsha, M.A. Winter, G. (2022). Acta Cryst. D78, 752–769.

Hamilton, W.C., Rollett, J.S. & Sparks, R.A. (1965). Acta Cryst. 18, 129–130.

Henderson, R., Baldwin, J.M., Ceska, T.A., Zemlin, F., Beckmann, E. & Downing, K.H. (1990). J. Mol. Biol. 213, 899–929.

Jumper, J., Evans, R., Pritzel A., Green, T., Figurnov, M., Ronneberger, O., Tunyasuvunakoo, K., Bates, R., Žídek, A., Potapenko, A., Bridgland, A., Meyer, C., A. Kohl, S.A.A., Ballard, A.J., Cowie, A., Romera-Paredes, B., Nikolov, S., Jain, R., Adler, J., Back, T., Petersen, S., Reiman, D., Clancy, E., Zielinski, M., Steinegger, M., Pacholska, M., Berghammer, T., Bodenstein, S., Silver, D., Vinyals, O., Senior, A.W., Kavukcuoglu, K., Kohli, P. & Hassabis, D. (2021). Nature. 596, 583–589.

Kabsch, W. (2010). Acta Cryst. D66, 125–132.

Karplus, P.A. & Diederichs, K. (2012). Science. 336, 1030–1033.

Karplus, P.A. & Diederichs, K. (2015). Curr. Opin. Struct. Biol. 34, 60–68.

Karuppan, S.J., Vogt, A., Fischer, Z., Ladutska, A., Swiastyn, J., McGraw, H.F., Bouyain, S. (2021). J.BiolChem. 298: 01541.

Shelley, K.L. & Garman, E.F. (2022). Nat. Commun. 13, 1314.

Leslie, A.G.W. & Powell, H.R. (2007). NATO Science Series, 245, Read, R.J., Sussman, J.L. (eds) Springer, Dordrecht.

Liebschner, D., Afonine, P.V., Baker, M.L., Bunkoczi, G., Chen, V.B., Croll, T.I., Hintze, B., Hung, L. W., Jain, S., McCoy, A.J., Moriarty, N.W., Oeffner, R.D., Poon, B.K., Prisant, M.J., Read, R.J., Richardson, J.R., Richardson, D.C., Sammito, M.D., Sobolev, O.V., Stockwell, D.H., Terwilliger, T.C, Urzhumtsev, A.G., Videau, L.L., Williams, C.J & Adams, P.D. (2019). Acta Cryst. D75, 861–877.

Maly, M., Diederichs, K., Dohnalek, J. & Kolenko, P. (2020). IUCrJ. 7, 681–692.

McCoy, A.J., Grosse-Kunstleve, R.W., Adams, P.D., Winn, M.D., Storoni, L.C. & Read, R.J. (2007). J. Appl. Cryst. 40, 658–674.

Mishra, N., Gido, C.D., Herdendorf, T.J., Hamme, M., Hura, G.L., Fu, Z.Q. & Geisbrecht, B.V. (2024). J. Biol. Chem. 300, 107627.

Murshudov, G.N., Skubak, P., Lebedev, A.A., Pannu, N.S., Steiner, R.A., Nicholls, R.A., Winn, M.D., Long, F. & Vagin, A.A. (2011). Acta Cryst. D67, 355–367.

Otwinowski, Z. & Minor, W. (1997). Methods Enzymol. 276, 307–326.

Pearce, N. M. & Gros, P. (2021). Nat. Commun. 12, 5493.

Pearson, K. (1895). Proc. Royal Soc. of London. 58, 240–242.

Ringe, D. & Petsko, G.A. (1986). Methods Enzymol. 131, 389–433.

Sun, Z., Liu, Q., Feng, Y. & Reetz, M.T. (2019). Chem. Rev. 119, 1626–1665.

Terwilliger, T.C, Grosse-Kunstleve, R.W., Afonine, P.V., Moriarty, N.W., Zwart, P.H., Hung, L.-W., Read, R.J. & Adams, P.D. (2008). Acta Cryst. D64, 61–69.

Waller, I. Zur Frage der Einwirkung der Wärmebewegung auf die Interferenz von Röntgenstrahlen. (1923). Z. Physik. 17, 398–408.

Weiss, M. & Hilgenfeld, R. (1997). J. Appl. Cryst. 30, 203–205.

Winn, M. D., Isupov, M. N. & Murshudov, G. N. (2001). Acta Cryst. D57, 122–133.

Winn, M.D, Ballard, C.C., Cowtan, K.D., Dodson, E.J., Emsley, P., Evans, P.R., Keegan, R.M., Krissinel, E.B., Leslie, A.G.W, McCoy, A., McNicholas, S.J., Murshudov, G.N., Pannu, N.S., Potterton, E.A., Powell, H.R., Read, R.J., Vaginc, A. and Wilson, K.S. (2011). Acta Cryst. D67, 235–242.

Winter, G., Waterman, D.G., Parkhurst, J.M., Brewster, A.S., Gildea, R.J., Gerstel, M., Fuentes-Montero, L., Vollmar, M., Michels-Clark, T., Young, I.D., Sauter, N.K. & Evans, G. (2018). Acta Cryst. D74, 85–97.

Zhang, C., Shine, M., Pyle, A.M., Zhang, Y. (2022). Nat Methods. 19(9),1109–1115.

