## supplementary1 for "I/σI *vs* {Rmerg, Rmeas, Rpim, CC1/2} for Crystal Diffraction Data Quality Evaluation"

<<< Supplementary_1 >>> Survey of 815 Data Sets with Various Characteristics
Article Title: Effective and Reliable Metrics to Select Crystal Diffraction Data

Listed below is a summary (unit cell removed to protect privacy) of statistics harvested during routine quality control from output of AIMLESS scaling after XDS indexing and integration of 815 data sets most recently collected at NYX beamline of NSLSII (SER-CATΓÇÖs partner beamline during APS dark period), with low-shell Rmerg<=0.08, high-shell CC1/2>=0.0 and Resolution better than 6Å. Multi, Compl, I/σIall are for the overall. The I/σIhigh and CC1/2high are for the highest-resolution shell.

| Reso | RmergAll | (Low | High) | Multi | Compl | I/σI(all | high) | CC1/2high | SpaceGroup |
|---|---|---|---|---|---|---|---|---|---|
| 1.47 | 0.028 | 0.015 | 0.853 | 3.2 | 93.1 | 17.0 | 1.0 | 0.482 | P121 |
| 1.29 | 0.027 | 0.016 | 1.039 | 1.8 | 79.8 | 11.8 | 0.9 | 0.345 | P1 |
| 1.48 | 0.029 | 0.016 | 1.203 | 3.4 | 99.8 | 17.0 | 0.9 | 0.413 | P2 |
| 1.49 | 0.035 | 0.017 | 2.246 | 3.8 | 98.8 | 14.9 | 0.7 | 0.320 | C2 |
| 3.44 | 0.074 | 0.017 | 1.184 | 6.4 | 95.6 | 15.5 | 1.7 | 0.216 | P622 |
| 1.43 | 0.033 | 0.017 | 1.141 | 3.2 | 91.2 | 16.1 | 0.9 | 0.398 | P2 |
| 1.61 | 0.064 | 0.018 | 0.596 | 6.1 | 98.5 | 11.8 | 0.1 | 0.007 | P212121 |
| 1.72 | 0.053 | 0.018 | 0.552 | 5.7 | 99.9 | 12.1 | 0.1 | 0.063 | P212121 |
| 1.95 | 0.049 | 0.018 | 0.937 | 3.2 | 96.0 | 13.8 | 1.0 | 0.437 | P2 |
| 3.48 | 0.109 | 0.018 | 2.624 | 8.7 | 98.4 | 15.6 | 1.5 | 0.363 | P622 |
| 1.38 | 0.035 | 0.018 | 1.311 | 6.5 | 100.0 | 20.0 | 1.0 | 0.446 | P212121 |
| 2.64 | 0.051 | 0.019 | 4.780 | 3.4 | 99.3 | 10.8 | 0.2 | 0.035 | P2 |
| 2.90 | 0.045 | 0.019 | 2.386 | 3.3 | 98.9 | 12.9 | 0.6 | 0.231 | P2 |
| 3.62 | 0.096 | 0.019 | 7.091 | 8.6 | 99.7 | 17.8 | 0.6 | 0.580 | P6122 |
| 1.34 | 0.038 | 0.019 | 1.599 | 6.4 | 99.9 | 18.2 | 0.9 | 0.327 | P222 |
| 1.48 | 0.057 | 0.019 | 2.161 | 6.6 | 99.9 | 19.3 | 1.1 | 0.499 | P212121 |
| 1.29 | 0.039 | 0.020 | 0.498 | 1.8 | 79.8 | 9.9 | 1.4 | 0.719 | P1 |
| 2.33 | 0.105 | 0.020 | 2.316 | 3.4 | 99.4 | 10.1 | 0.7 | 0.279 | P2 |
| 3.99 | 0.132 | 0.020 | 2.488 | 7.3 | 93.8 | 10.5 | 0.7 | 0.283 | P321 |
| 4.13 | 0.118 | 0.020 | 1.820 | 7.2 | 94.3 | 11.6 | 1.0 | 0.421 | P321 |
| 2.76 | 0.061 | 0.020 | 1.337 | 3.4 | 99.4 | 12.6 | 0.9 | 0.363 | P1211 |
| 1.53 | 0.042 | 0.020 | 0.628 | 6.2 | 100.0 | 14.1 | 0.1 | 0.023 | C121 |
| 1.46 | 0.033 | 0.020 | 0.876 | 3.3 | 96.9 | 14.5 | 1.0 | 0.483 | P121 |
| 2.33 | 0.064 | 0.021 | 1.663 | 3.7 | 99.0 | 12.0 | 0.8 | 0.255 | C2 |
| 3.04 | 0.050 | 0.021 | 2.246 | 6.5 | 99.9 | 17.1 | 0.8 | 0.403 | C222 |
| 1.40 | 0.043 | 0.021 | 2.924 | 7.1 | 99.4 | 17.5 | 0.6 | 0.189 | I222 |
| 2.29 | 0.062 | 0.021 | 1.718 | 7.3 | 100.0 | 18.9 | 1.1 | 0.367 | I222 |
| 1.49 | 0.060 | 0.021 | 4.494 | 6.6 | 99.6 | 19.3 | 1.5 | 0.639 | P222 |
| 1.48 | 0.039 | 0.021 | 1.435 | 7.3 | 99.9 | 21.0 | 1.2 | 0.513 | I222 |
| 1.69 | 0.034 | 0.021 | 0.578 | 15.7 | 100.0 | 31.9 | 0.3 | 0.325 | P41212 |
| 1.48 | 0.045 | 0.022 | 1.133 | 2.0 | 95.5 | 9.3 | 1.5 | 0.239 | P1 |
| 2.63 | 0.077 | 0.022 | 2.333 | 3.5 | 99.3 | 10.4 | 0.6 | 0.222 | P2 |
| 2.15 | 0.055 | 0.022 | 0.577 | 3.1 | 99.6 | 10.7 | 0.2 | 0.049 | I121 |
| 4.15 | 0.146 | 0.022 | 2.472 | 6.8 | 99.8 | 10.7 | 0.9 | 0.324 | P3121 |
| 1.62 | 0.057 | 0.022 | 0.663 | 5.9 | 99.8 | 12.3 | 0.1 | 0.022 | P212121 |
| 2.10 | 0.080 | 0.022 | 4.388 | 7.4 | 100.0 | 14.2 | 0.5 | 0.114 | I222 |
| 2.26 | 0.082 | 0.022 | 0.575 | 6.1 | 99.1 | 14.8 | 0.4 | 0.197 | P212121 |
| 1.48 | 0.062 | 0.022 | 1.654 | 6.6 | 100.0 | 16.8 | 1.2 | 0.336 | P212121 |
| 3.04 | 0.047 | 0.022 | 2.481 | 6.5 | 99.6 | 17.4 | 0.7 | 0.397 | P222 |
| 1.94 | 0.053 | 0.022 | 1.055 | 5.8 | 98.2 | 18.4 | 0.9 | 0.383 | P212121 |

| | | | | | | | | | |
|---|---|---|---|---|---|---|---|---|---|
| 1.72 | 0.064 | 0.022 | 2.038 | 6.6 | 98.8 | 21.3 | 2.5 | 0.768 | P212121 |
| 1.88 | 0.061 | 0.023 | 2.148 | 3.4 | 99.3 | 9.3 | 0.5 | 0.173 | P2 |
| 1.86 | 0.059 | 0.023 | 2.017 | 3.9 | 99.8 | 11.1 | 0.7 | 0.228 | P2 |
| 2.27 | 0.085 | 0.023 | 0.586 | 6.2 | 99.8 | 11.8 | 0.1 | 0.079 | P212121 |
| 1.61 | 0.042 | 0.023 | 1.394 | 3.4 | 93.8 | 12.8 | 0.7 | 0.298 | P121 |
| 1.74 | 0.067 | 0.023 | 3.089 | 6.7 | 99.8 | 13.4 | 0.6 | 0.293 | C222 |
| 1.42 | 0.039 | 0.023 | 1.116 | 3.2 | 95.4 | 13.4 | 0.8 | 0.401 | P2 |
| 1.57 | 0.048 | 0.023 | 0.629 | 6.1 | 100.0 | 14.1 | 0.2 | 0.052 | P1211 |
| 1.79 | 0.061 | 0.023 | 2.272 | 6.7 | 99.9 | 15.0 | 0.7 | 0.366 | C2221 |
| 2.11 | 0.047 | 0.023 | 2.389 | 5.7 | 99.9 | 15.7 | 0.9 | 0.243 | R3 |
| 1.47 | 0.030 | 0.023 | 0.955 | 3.1 | 86.9 | 17.1 | 1.5 | 0.879 | P121 |
| 2.19 | 0.042 | 0.023 | 1.485 | 5.7 | 99.9 | 18.2 | 1.1 | 0.327 | H3 |
| 1.53 | 0.035 | 0.023 | 1.984 | 7.4 | 99.9 | 21.6 | 0.8 | 0.519 | I222 |
| 1.48 | 0.035 | 0.023 | 1.331 | 7.3 | 100.0 | 22.9 | 1.2 | 0.592 | I222 |
| 1.47 | 0.034 | 0.023 | 1.720 | 6.6 | 100.0 | 23.2 | 1.0 | 0.445 | P212121 |
| 1.69 | 0.096 | 0.024 | 0.851 | 1.8 | 89.1 | 5.2 | 0.9 | 0.411 | P1 |
| 2.46 | 0.090 | 0.024 | 1.429 | 1.8 | 96.9 | 5.4 | 0.9 | 0.137 | P1 |
| 2.45 | 0.085 | 0.024 | 0.812 | 3.1 | 99.7 | 7.2 | 0.1 | 0.103 | C121 |
| 1.74 | 0.040 | 0.024 | 0.607 | 1.9 | 89.3 | 7.5 | 0.2 | 0.062 | P1 |
| 4.01 | 0.166 | 0.024 | 3.643 | 6.8 | 99.9 | 9.6 | 0.7 | 0.239 | P321 |
| 2.34 | 0.091 | 0.024 | 0.642 | 5.9 | 100.0 | 9.8 | 0.1 | 0.027 | P212121 |
| 1.70 | 0.050 | 0.024 | 0.554 | 6.1 | 100.0 | 12.3 | 0.1 | 0.070 | P1211 |
| 1.57 | 0.047 | 0.024 | 1.968 | 3.4 | 92.1 | 12.9 | 0.6 | 0.251 | P2 |
| 2.00 | 0.048 | 0.024 | 1.223 | 4.1 | 98.9 | 13.5 | 1.2 | 0.500 | P1211 |
| 2.09 | 0.092 | 0.024 | 3.932 | 8.9 | 99.9 | 14.3 | 0.6 | 0.183 | I222 |
| 1.95 | 0.067 | 0.024 | 1.996 | 6.6 | 100.0 | 14.8 | 0.8 | 0.337 | P212121 |
| 1.72 | 0.120 | 0.024 | 3.515 | 6.5 | 95.7 | 15.3 | 0.8 | 0.127 | P212121 |
| 1.59 | 0.077 | 0.024 | 4.011 | 6.6 | 99.4 | 16.7 | 1.1 | 0.578 | P212121 |
| 2.30 | 0.072 | 0.024 | 2.133 | 7.2 | 99.9 | 17.1 | 0.9 | 0.294 | I222 |
| 2.20 | 0.079 | 0.024 | 2.413 | 8.9 | 100.0 | 17.2 | 0.9 | 0.288 | I222 |
| 1.44 | 0.041 | 0.024 | 2.218 | 7.3 | 99.7 | 18.2 | 0.9 | 0.401 | I222 |
| 1.40 | 0.038 | 0.024 | 2.412 | 7.1 | 99.4 | 19.1 | 0.6 | 0.295 | I222 |
| 1.49 | 0.038 | 0.024 | 2.684 | 7.4 | 99.8 | 19.2 | 0.7 | 0.426 | I222 |
| 1.48 | 0.038 | 0.024 | 1.639 | 7.4 | 99.9 | 20.2 | 1.0 | 0.524 | I222 |
| 1.50 | 0.042 | 0.024 | 1.008 | 6.0 | 98.9 | 20.3 | 1.2 | 0.587 | P212121 |
| 1.48 | 0.046 | 0.025 | 0.652 | 1.5 | 46.2 | 7.8 | 0.7 | 0.558 | I121 |
| 1.45 | 0.037 | 0.025 | 1.164 | 1.8 | 78.6 | 9.6 | 0.9 | 0.274 | P1 |
| 2.77 | 0.110 | 0.025 | 2.043 | 3.4 | 98.9 | 9.9 | 0.8 | 0.338 | P1211 |
| 1.87 | 0.060 | 0.025 | 2.787 | 4.2 | 98.5 | 10.8 | 0.6 | 0.214 | P2 |
| 1.61 | 0.055 | 0.025 | 0.589 | 5.7 | 100.0 | 11.6 | 0.1 | 0.011 | P212121 |
| 1.79 | 0.090 | 0.025 | 2.185 | 6.6 | 99.8 | 12.0 | 0.7 | 0.381 | P212121 |
| 1.88 | 0.075 | 0.025 | 2.962 | 6.7 | 99.9 | 12.6 | 0.6 | 0.233 | P222 |
| 1.60 | 0.056 | 0.025 | 0.603 | 5.9 | 99.9 | 14.1 | 0.2 | 0.104 | P212121 |
| 1.49 | 0.076 | 0.025 | 4.734 | 6.6 | 99.7 | 15.7 | 1.2 | 0.484 | P212121 |
| 1.92 | 0.065 | 0.025 | 1.974 | 7.4 | 99.8 | 16.3 | 1.1 | 0.443 | P212121 |
| 2.00 | 0.063 | 0.025 | 1.577 | 7.4 | 100.0 | 16.8 | 1.2 | 0.514 | P212121 |
| 1.42 | 0.047 | 0.025 | 2.011 | 5.7 | 94.6 | 18.0 | 0.6 | 0.355 | P222 |
| 1.66 | 0.074 | 0.025 | 70.286 | 6.6 | 99.1 | 18.5 | 1.8 | 0.643 | P222 |
| 1.48 | 0.035 | 0.025 | 1.615 | 8.1 | 99.9 | 23.7 | 1.1 | 0.641 | I222 |
| 2.86 | 0.178 | 0.026 | 3.798 | 1.2 | 15.6 | 3.2 | 0.6 | 0.082 | C2 |
| 2.86 | 0.178 | 0.026 | 3.798 | 1.2 | 15.6 | 3.2 | 0.6 | 0.082 | C2 |
| 1.85 | 0.057 | 0.026 | 1.194 | 3.4 | 98.2 | 9.8 | 1.1 | 0.631 | C121 |
| 1.75 | 0.100 | 0.026 | 3.026 | 6.6 | 99.7 | 11.1 | 0.6 | 0.265 | P222 |
| 1.87 | 0.082 | 0.026 | 3.232 | 7.5 | 99.5 | 12.7 | 0.6 | 0.195 | P222 |

| | | | | | | | | | |
|---|---|---|---|---|---|---|---|---|---|
| 1.87 | 0.074 | 0.026 | 3.240 | 7.5 | 100.0 | 12.9 | 0.6 | 0.248 | P222 |
| 2.50 | 0.065 | 0.026 | 2.895 | 5.6 | 99.9 | 13.1 | 0.8 | 0.192 | R3 |
| 2.77 | 0.057 | 0.026 | 2.651 | 6.5 | 99.8 | 13.8 | 0.7 | 0.258 | P222 |
| 1.86 | 0.068 | 0.026 | 2.481 | 6.7 | 98.8 | 14.1 | 0.7 | 0.254 | P21212 |
| 1.86 | 0.071 | 0.026 | 2.561 | 7.5 | 99.7 | 14.2 | 0.9 | 0.364 | P222 |
| 1.93 | 0.075 | 0.026 | 2.583 | 7.5 | 99.7 | 14.6 | 0.7 | 0.304 | P212121 |
| 1.45 | 0.034 | 0.026 | 0.378 | 3.1 | 87.0 | 17.2 | 1.7 | 0.891 | P2 |
| 1.92 | 0.062 | 0.026 | 1.526 | 7.4 | 100.0 | 17.7 | 1.2 | 0.525 | P212121 |
| 1.50 | 0.043 | 0.026 | 0.582 | 6.1 | 99.9 | 18.0 | 0.4 | 0.214 | P1211 |
| 1.38 | 0.036 | 0.026 | 1.253 | 6.6 | 99.5 | 19.6 | 1.1 | 0.513 | P212121 |
| 2.63 | 0.121 | 0.027 | 0.725 | 1.1 | 16.7 | 3.7 | 0.9 | 0.506 | C2 |
| 2.63 | 0.121 | 0.027 | 0.725 | 1.1 | 16.7 | 3.7 | 0.9 | 0.506 | C2 |
| 2.47 | 0.092 | 0.027 | 0.955 | 1.2 | 16.0 | 5.5 | 1.0 | 0.419 | C2 |
| 2.47 | 0.092 | 0.027 | 0.955 | 1.2 | 16.0 | 5.5 | 1.0 | 0.419 | C2 |
| 2.58 | 0.108 | 0.027 | 0.585 | 5.8 | 100.0 | 8.6 | 0.1 | 0.041 | P212121 |
| 1.53 | 0.048 | 0.027 | 0.510 | 3.1 | 92.1 | 9.8 | 0.9 | 0.743 | P121 |
| 2.01 | 0.081 | 0.027 | 1.802 | 6.7 | 99.9 | 11.5 | 1.1 | 0.523 | P212121 |
| 1.80 | 0.075 | 0.027 | 3.676 | 6.8 | 98.4 | 12.0 | 0.6 | 0.170 | P222 |
| 2.21 | 0.080 | 0.027 | 2.951 | 8.8 | 99.9 | 14.4 | 0.8 | 0.286 | I222 |
| 2.17 | 0.058 | 0.027 | 0.566 | 10.3 | 100.0 | 15.3 | 0.1 | 0.048 | P41212 |
| 2.30 | 0.073 | 0.027 | 2.059 | 8.8 | 100.0 | 16.9 | 1.0 | 0.417 | I222 |
| 1.54 | 0.068 | 0.027 | 0.624 | 6.0 | 98.6 | 17.2 | 0.8 | 0.219 | P1211 |
| 2.00 | 0.058 | 0.027 | 2.661 | 8.9 | 100.0 | 17.5 | 0.8 | 0.369 | I222 |
| 1.55 | 0.065 | 0.027 | 0.571 | 6.1 | 92.3 | 18.7 | 0.9 | 0.583 | P1211 |
| 1.48 | 0.064 | 0.027 | 0.569 | 6.2 | 92.1 | 19.1 | 1.1 | 0.568 | P1211 |
| 1.44 | 0.037 | 0.027 | 1.987 | 8.1 | 99.8 | 21.4 | 0.9 | 0.542 | I222 |
| 2.81 | 0.152 | 0.028 | 1.648 | 1.1 | 16.6 | 3.2 | 0.6 | 0.370 | C2 |
| 2.81 | 0.152 | 0.028 | 1.648 | 1.1 | 16.6 | 3.2 | 0.6 | 0.370 | C2 |
| 2.46 | 0.117 | 0.028 | 0.965 | 1.1 | 16.7 | 3.6 | 0.5 | 0.127 | C2 |
| 2.46 | 0.117 | 0.028 | 0.965 | 1.1 | 16.7 | 3.6 | 0.5 | 0.127 | C2 |
| 1.38 | 0.050 | 0.028 | 1.395 | 3.4 | 98.9 | 10.1 | 0.9 | 0.253 | P2 |
| 1.88 | 0.046 | 0.028 | 0.783 | 2.3 | 68.8 | 12.2 | 1.8 | 0.775 | P121 |
| 1.94 | 0.062 | 0.028 | 3.322 | 9.0 | 100.0 | 15.5 | 0.6 | 0.271 | I222 |
| 1.30 | 0.040 | 0.028 | 1.821 | 6.3 | 97.4 | 17.0 | 0.7 | 0.299 | P222 |
| 3.01 | 0.065 | 0.028 | 3.488 | 12.6 | 99.9 | 24.0 | 0.7 | 0.334 | P41212 |
| 1.54 | 0.073 | 0.029 | 0.800 | 1.8 | 93.6 | 7.1 | 2.1 | 0.313 | P1 |
| 1.74 | 0.069 | 0.029 | 1.967 | 3.4 | 96.4 | 8.0 | 0.6 | 0.408 | C2 |
| 1.46 | 0.054 | 0.029 | 0.517 | 3.0 | 87.0 | 9.3 | 0.8 | 0.725 | P2 |
| 1.94 | 0.091 | 0.029 | 2.593 | 6.7 | 99.9 | 10.0 | 0.9 | 0.433 | P222 |
| 2.57 | 0.065 | 0.029 | 1.225 | 3.4 | 99.5 | 10.1 | 0.9 | 0.370 | P1211 |
| 1.88 | 0.073 | 0.029 | 2.870 | 6.6 | 100.0 | 11.9 | 0.7 | 0.272 | P222 |
| 1.95 | 0.066 | 0.029 | 1.967 | 6.6 | 100.0 | 13.4 | 0.9 | 0.405 | P212121 |
| 1.51 | 0.036 | 0.029 | 0.654 | 3.2 | 95.9 | 13.4 | 1.2 | 0.651 | P121 |
| 2.60 | 0.059 | 0.029 | 1.637 | 5.6 | 99.9 | 15.5 | 1.0 | 0.216 | H3 |
| 2.63 | 0.038 | 0.029 | 0.844 | 30.1 | 100.0 | 24.4 | 0.1 | 0.027 | P6122 |
| 2.32 | 0.100 | 0.030 | 1.899 | 2.4 | 97.3 | 6.2 | 0.6 | 0.127 | P1 |
| 1.59 | 0.037 | 0.030 | 0.671 | 1.9 | 87.3 | 8.1 | 0.4 | 0.129 | P1 |
| 2.46 | 0.076 | 0.030 | 1.969 | 3.4 | 99.4 | 8.6 | 0.6 | 0.242 | P2 |
| 2.07 | 0.111 | 0.030 | 3.436 | 6.6 | 99.9 | 9.3 | 0.5 | 0.165 | P222 |
| 2.67 | 0.121 | 0.030 | 1.110 | 3.3 | 99.2 | 9.8 | 1.5 | 0.335 | P1211 |
| 1.72 | 0.089 | 0.030 | 0.605 | 5.9 | 99.9 | 11.3 | 0.4 | 0.252 | P1211 |
| 2.32 | 0.129 | 0.030 | 2.868 | 7.3 | 99.9 | 11.3 | 0.7 | 0.229 | I222 |
| 2.58 | 0.165 | 0.030 | 3.296 | 6.9 | 99.8 | 11.7 | 0.7 | 0.409 | I121 |
| 2.16 | 0.097 | 0.030 | 2.136 | 6.6 | 99.9 | 11.7 | 0.8 | 0.273 | P212121 |

| | | | | | | | | | |
|---|---|---|---|---|---|---|---|---|---|
| 1.71 | 0.081 | 0.030 | 0.563 | 6.0 | 100.0 | 12.8 | 0.3 | 0.325 | P1211 |
| 2.05 | 0.076 | 0.030 | 1.598 | 6.5 | 100.0 | 13.4 | 1.1 | 0.449 | P212121 |
| 2.18 | 0.083 | 0.030 | 2.705 | 8.6 | 99.9 | 13.8 | 0.8 | 0.315 | I222 |
| 2.79 | 0.068 | 0.030 | 5.033 | 12.6 | 99.9 | 15.2 | 0.5 | 0.249 | P422 |
| 3.02 | 0.060 | 0.030 | 1.912 | 12.3 | 100.0 | 21.0 | 1.2 | 0.741 | P41212 |
| 1.42 | 0.056 | 0.030 | 0.507 | 6.3 | 84.5 | 21.8 | 1.3 | 0.642 | P1211 |
| 2.71 | 0.175 | 0.031 | 0.585 | 2.0 | 89.3 | 2.8 | 0.7 | 0.149 | H32 |
| 1.88 | 0.094 | 0.031 | 4.570 | 6.6 | 99.9 | 9.6 | 0.4 | 0.155 | P222 |
| 1.59 | 0.060 | 0.031 | 0.631 | 3.5 | 91.9 | 10.0 | 0.7 | 0.400 | I121 |
| 2.10 | 0.094 | 0.031 | 3.753 | 8.7 | 99.9 | 10.8 | 0.6 | 0.239 | I222 |
| 1.99 | 0.061 | 0.031 | 0.722 | 6.2 | 98.8 | 10.9 | 0.2 | 0.116 | I121 |
| 2.89 | 0.068 | 0.031 | 4.662 | 6.6 | 99.8 | 11.4 | 0.4 | 0.076 | P4 |
| 1.43 | 0.041 | 0.031 | 1.234 | 3.2 | 91.1 | 12.3 | 0.8 | 0.363 | P2 |
| 2.42 | 0.113 | 0.031 | 2.109 | 7.4 | 100.0 | 13.3 | 0.8 | 0.297 | I222 |
| 1.93 | 0.069 | 0.031 | 1.777 | 6.6 | 99.9 | 13.3 | 1.0 | 0.400 | P212121 |
| 1.63 | 0.077 | 0.031 | 0.580 | 6.0 | 100.0 | 14.3 | 0.5 | 0.211 | P1211 |
| 2.78 | 0.077 | 0.031 | 5.265 | 12.5 | 99.9 | 14.8 | 0.6 | 0.211 | P422 |
| 1.58 | 0.090 | 0.031 | 0.569 | 6.5 | 89.1 | 16.1 | 0.9 | 0.439 | P121 |
| 2.86 | 0.217 | 0.032 | 2.391 | 1.4 | 13.9 | 3.4 | 0.4 | 0.097 | C2 |
| 2.86 | 0.217 | 0.032 | 2.391 | 1.4 | 13.9 | 3.4 | 0.4 | 0.097 | C2 |
| 3.12 | 0.221 | 0.032 | 1.983 | 1.6 | 11.5 | 3.6 | 0.6 | 0.222 | C2 |
| 3.12 | 0.221 | 0.032 | 1.983 | 1.6 | 11.5 | 3.6 | 0.6 | 0.222 | C2 |
| 2.20 | 0.120 | 0.032 | 0.586 | 3.8 | 91.8 | 7.2 | 0.7 | 0.426 | P222 |
| 2.93 | 0.150 | 0.032 | 0.958 | 7.9 | 79.8 | 7.8 | 2.0 | 0.317 | P622 |
| 3.05 | 0.063 | 0.032 | 2.172 | 3.3 | 99.1 | 8.7 | 0.7 | 0.307 | P2 |
| 2.01 | 0.232 | 0.032 | 1.510 | 3.2 | 97.7 | 9.0 | 1.2 | 0.221 | P2 |
| 3.40 | 0.125 | 0.032 | 4.427 | 6.3 | 99.8 | 10.4 | 0.6 | 0.162 | P222 |
| 1.72 | 0.101 | 0.032 | 0.596 | 5.9 | 99.5 | 11.5 | 0.6 | 0.323 | P1211 |
| 1.48 | 0.105 | 0.032 | 7.855 | 6.9 | 97.9 | 11.6 | 0.9 | 0.271 | C2 |
| 1.87 | 0.076 | 0.032 | 2.383 | 6.7 | 99.9 | 11.7 | 0.8 | 0.312 | P222 |
| 2.64 | 0.165 | 0.032 | 8.464 | 6.9 | 99.2 | 11.8 | 1.0 | 0.445 | C2 |
| 1.97 | 0.085 | 0.032 | 2.617 | 6.6 | 99.9 | 11.9 | 0.8 | 0.266 | P222 |
| 2.00 | 0.081 | 0.032 | 1.444 | 7.4 | 100.0 | 12.8 | 1.1 | 0.517 | P212121 |
| 1.94 | 0.076 | 0.032 | 1.915 | 6.6 | 99.9 | 13.0 | 0.9 | 0.385 | P212121 |
| 2.05 | 0.076 | 0.032 | 1.735 | 6.6 | 100.0 | 13.9 | 1.0 | 0.374 | P212121 |
| 1.65 | 0.135 | 0.032 | 0.913 | 6.4 | 93.3 | 14.0 | 2.8 | 0.336 | P212121 |
| 2.09 | 0.066 | 0.032 | 1.794 | 7.5 | 100.0 | 15.2 | 1.0 | 0.396 | I222 |
| 1.74 | 0.059 | 0.032 | 0.982 | 6.6 | 98.7 | 15.2 | 1.2 | 0.699 | P1211 |
| 1.74 | 0.059 | 0.032 | 0.982 | 6.6 | 98.7 | 15.2 | 1.2 | 0.699 | P1211 |
| 2.89 | 0.065 | 0.032 | 5.076 | 12.7 | 99.9 | 17.6 | 0.5 | 0.234 | P422 |
| 2.88 | 0.072 | 0.032 | 3.298 | 12.4 | 100.0 | 18.2 | 0.7 | 0.345 | P41212 |
| 3.00 | 0.063 | 0.032 | 3.409 | 12.6 | 100.0 | 20.6 | 0.7 | 0.498 | P41212 |
| 2.85 | 0.178 | 0.033 | 1.304 | 1.4 | 14.0 | 3.6 | 0.5 | 0.288 | C2 |
| 2.85 | 0.178 | 0.033 | 1.304 | 1.4 | 14.0 | 3.6 | 0.5 | 0.288 | C2 |
| 1.86 | 0.149 | 0.033 | 3.369 | 7.5 | 99.7 | 9.8 | 0.6 | 0.258 | P222 |
| 1.86 | 0.093 | 0.033 | 3.067 | 7.4 | 100.0 | 10.0 | 0.6 | 0.219 | P222 |
| 1.76 | 0.095 | 0.033 | 0.575 | 6.1 | 99.3 | 10.5 | 0.5 | 0.202 | P1211 |
| 1.50 | 0.132 | 0.033 | 2.426 | 6.2 | 97.9 | 10.6 | 0.9 | 0.093 | P222 |
| 1.88 | 0.084 | 0.033 | 2.834 | 6.6 | 99.9 | 10.9 | 0.7 | 0.264 | P222 |
| 1.58 | 0.116 | 0.033 | 1.416 | 6.4 | 99.6 | 12.1 | 1.0 | 0.277 | P212121 |
| 2.08 | 0.103 | 0.033 | 2.009 | 7.3 | 100.0 | 12.5 | 0.9 | 0.341 | P212121 |
| 1.74 | 0.121 | 0.033 | 3.603 | 7.3 | 99.9 | 12.5 | 1.5 | 0.245 | I222 |
| 1.85 | 0.072 | 0.033 | 1.474 | 6.7 | 99.9 | 12.8 | 1.1 | 0.557 | P212121 |
| 2.32 | 0.109 | 0.033 | 2.645 | 8.8 | 100.0 | 13.2 | 0.8 | 0.280 | I222 |

| | | | | | | | | | |
|---|---|---|---|---|---|---|---|---|---|
| 2.29 | 0.123 | 0.033 | 2.652 | 6.6 | 99.9 | 13.8 | 0.8 | 0.395 | P212121 |
| 1.58 | 0.057 | 0.033 | 0.754 | 6.0 | 89.8 | 15.8 | 1.0 | 0.235 | P212121 |
| 1.60 | 0.062 | 0.033 | 1.188 | 10.7 | 99.0 | 16.8 | 1.0 | 0.643 | P422 |
| 3.28 | 0.082 | 0.033 | 4.024 | 12.5 | 100.0 | 17.1 | 0.8 | 0.321 | P21221 |
| 1.65 | 0.060 | 0.033 | 0.971 | 11.1 | 99.5 | 18.0 | 1.0 | 0.739 | P41212 |
| 1.51 | 0.048 | 0.033 | 0.879 | 7.6 | 98.4 | 19.8 | 1.3 | 0.680 | I222 |
| 2.86 | 0.170 | 0.034 | 3.001 | 1.5 | 12.9 | 3.1 | 0.4 | 0.159 | C2 |
| 2.86 | 0.170 | 0.034 | 3.001 | 1.5 | 12.9 | 3.1 | 0.4 | 0.159 | C2 |
| 2.62 | 0.138 | 0.034 | 0.983 | 1.1 | 16.7 | 3.7 | 0.6 | 0.158 | C2 |
| 2.62 | 0.138 | 0.034 | 0.983 | 1.1 | 16.7 | 3.7 | 0.6 | 0.158 | C2 |
| 2.33 | 0.096 | 0.034 | 0.524 | 1.1 | 16.3 | 4.0 | 0.7 | 0.748 | C2 |
| 2.33 | 0.096 | 0.034 | 0.524 | 1.1 | 16.3 | 4.0 | 0.7 | 0.748 | C2 |
| 2.91 | 0.070 | 0.034 | 1.378 | 3.4 | 97.7 | 7.5 | 0.8 | 0.447 | C2 |
| 2.74 | 0.214 | 0.034 | 1.718 | 3.3 | 98.9 | 8.3 | 2.3 | 0.210 | P2 |
| 2.12 | 0.147 | 0.034 | 4.303 | 7.3 | 100.0 | 8.6 | 0.6 | 0.152 | P222 |
| 1.89 | 0.090 | 0.034 | 3.032 | 6.6 | 99.9 | 9.5 | 0.6 | 0.206 | P222 |
| 1.75 | 0.081 | 0.034 | 3.197 | 6.6 | 99.6 | 10.5 | 0.5 | 0.216 | P222 |
| 2.01 | 0.114 | 0.034 | 2.856 | 7.3 | 100.0 | 10.7 | 0.8 | 0.254 | P222 |
| 1.95 | 0.082 | 0.034 | 2.178 | 6.5 | 99.9 | 11.3 | 0.8 | 0.312 | P212121 |
| 1.92 | 0.131 | 0.034 | 2.517 | 7.4 | 99.8 | 11.4 | 0.7 | 0.354 | P212121 |
| 2.21 | 0.159 | 0.034 | 5.450 | 13.5 | 99.9 | 12.1 | 0.6 | 0.401 | P222 |
| 1.95 | 0.073 | 0.034 | 0.911 | 5.8 | 92.0 | 12.4 | 1.2 | 0.788 | C222 |
| 1.95 | 0.073 | 0.034 | 0.911 | 5.8 | 92.0 | 12.4 | 1.2 | 0.788 | C222 |
| 2.21 | 0.151 | 0.034 | 5.170 | 13.4 | 99.9 | 12.7 | 0.7 | 0.454 | P222 |
| 1.65 | 0.066 | 0.034 | 1.631 | 6.4 | 95.5 | 13.1 | 0.8 | 0.507 | P2 |
| 1.65 | 0.066 | 0.034 | 1.631 | 6.4 | 95.5 | 13.1 | 0.8 | 0.507 | P2 |
| 3.12 | 0.090 | 0.034 | 4.830 | 12.5 | 99.9 | 13.8 | 0.5 | 0.225 | P222 |
| 1.38 | 0.055 | 0.034 | 1.597 | 6.4 | 99.5 | 14.4 | 0.9 | 0.315 | P212121 |
| 2.90 | 0.078 | 0.034 | 4.275 | 12.4 | 99.9 | 14.6 | 0.5 | 0.237 | P422 |
| 1.65 | 0.084 | 0.034 | 0.558 | 6.1 | 98.0 | 15.0 | 0.8 | 0.496 | P1211 |
| 2.30 | 0.138 | 0.034 | 3.693 | 13.4 | 99.9 | 15.2 | 0.9 | 0.562 | P212121 |
| 1.51 | 0.077 | 0.034 | 0.589 | 5.9 | 97.6 | 15.6 | 0.8 | 0.375 | P1211 |
| 1.59 | 0.099 | 0.034 | 1.815 | 6.4 | 99.8 | 15.7 | 1.2 | 0.249 | P212121 |
| 2.42 | 0.099 | 0.034 | 1.808 | 8.8 | 100.0 | 15.9 | 1.1 | 0.346 | I222 |
| 3.01 | 0.074 | 0.034 | 2.676 | 12.4 | 100.0 | 17.6 | 0.8 | 0.596 | P41212 |
| 1.43 | 0.051 | 0.034 | 1.884 | 7.3 | 95.4 | 17.7 | 0.8 | 0.391 | I222 |
| 3.00 | 0.068 | 0.034 | 3.238 | 12.4 | 100.0 | 19.6 | 0.8 | 0.356 | P41212 |
| 1.47 | 0.120 | 0.034 | 3.429 | 20.0 | 100.0 | 20.0 | 1.1 | 0.365 | P3121 |
| 2.84 | 0.137 | 0.035 | 1.333 | 1.1 | 16.5 | 3.6 | 0.9 | 0.313 | C2 |
| 2.84 | 0.137 | 0.035 | 1.333 | 1.1 | 16.5 | 3.6 | 0.9 | 0.313 | C2 |
| 3.77 | 0.143 | 0.035 | 0.780 | 1.6 | 11.3 | 3.9 | 0.9 | 0.579 | C2 |
| 3.77 | 0.143 | 0.035 | 0.780 | 1.6 | 11.3 | 3.9 | 0.9 | 0.579 | C2 |
| 2.33 | 0.092 | 0.035 | 0.794 | 1.2 | 15.9 | 4.6 | 0.9 | 0.389 | C2 |
| 2.33 | 0.092 | 0.035 | 0.794 | 1.2 | 15.9 | 4.6 | 0.9 | 0.389 | C2 |
| 2.46 | 0.187 | 0.035 | 0.885 | 6.6 | 72.9 | 5.5 | 1.0 | 0.216 | P622 |
| 1.49 | 0.049 | 0.035 | 0.593 | 1.9 | 85.1 | 6.4 | 0.6 | 0.405 | P1 |
| 1.94 | 0.201 | 0.035 | 4.259 | 6.6 | 99.9 | 7.3 | 0.6 | 0.253 | P222 |
| 2.01 | 0.089 | 0.035 | 1.199 | 3.3 | 99.3 | 7.4 | 1.0 | 0.543 | C121 |
| 1.90 | 0.122 | 0.035 | 0.681 | 6.0 | 100.0 | 8.3 | 0.2 | 0.125 | P1211 |
| 2.09 | 0.158 | 0.035 | 1.918 | 6.4 | 99.9 | 9.5 | 1.1 | 0.443 | P212121 |
| 2.18 | 0.063 | 0.035 | 1.110 | 4.0 | 99.5 | 9.8 | 1.1 | 0.595 | P1211 |
| 2.20 | 0.132 | 0.035 | 2.831 | 7.3 | 100.0 | 10.4 | 0.7 | 0.254 | P212121 |
| 1.63 | 0.080 | 0.035 | 0.630 | 5.7 | 99.7 | 11.4 | 0.3 | 0.089 | P212121 |
| 2.02 | 0.073 | 0.035 | 2.180 | 7.5 | 100.0 | 13.4 | 0.9 | 0.368 | I222 |

| | | | | | | | | | |
|---|---|---|---|---|---|---|---|---|---|
| 2.33 | 0.172 | 0.035 | 4.192 | 13.4 | 99.9 | 13.8 | 0.8 | 0.559 | P222 |
| 1.79 | 0.107 | 0.035 | 1.775 | 7.2 | 99.9 | 14.0 | 1.6 | 0.301 | I222 |
| 2.30 | 0.145 | 0.035 | 4.187 | 13.4 | 100.0 | 14.2 | 0.8 | 0.465 | P212121 |
| 2.01 | 0.079 | 0.036 | 2.704 | 4.1 | 99.5 | 7.5 | 0.6 | 0.246 | P2 |
| 2.13 | 0.124 | 0.036 | 1.313 | 3.2 | 94.4 | 7.6 | 1.2 | 0.222 | P121 |
| 1.90 | 0.087 | 0.036 | 0.560 | 8.3 | 99.9 | 7.8 | 0.4 | 0.283 | P212121 |
| 2.08 | 0.113 | 0.036 | 2.480 | 6.6 | 99.8 | 8.7 | 0.8 | 0.255 | P222 |
| 4.32 | 0.128 | 0.036 | 1.961 | 7.1 | 94.7 | 8.9 | 1.0 | 0.337 | P321 |
| 2.13 | 0.128 | 0.036 | 2.473 | 6.5 | 99.9 | 10.2 | 0.7 | 0.253 | P212121 |
| 2.17 | 0.102 | 0.036 | 1.808 | 6.6 | 99.9 | 10.3 | 1.0 | 0.381 | P212121 |
| 1.75 | 0.093 | 0.036 | 0.577 | 6.1 | 99.7 | 10.5 | 0.3 | 0.219 | P1211 |
| 2.08 | 0.101 | 0.036 | 2.198 | 7.3 | 100.0 | 10.8 | 0.8 | 0.402 | P212121 |
| 2.29 | 0.105 | 0.036 | 2.073 | 6.7 | 99.8 | 11.9 | 1.3 | 0.678 | I121 |
| 1.86 | 0.094 | 0.036 | 6.461 | 13.5 | 99.2 | 12.4 | 0.4 | 0.303 | F222 |
| 4.32 | 0.100 | 0.036 | 2.430 | 8.1 | 100.0 | 13.1 | 1.6 | 0.695 | P6122 |
| 2.19 | 0.073 | 0.036 | 2.124 | 7.4 | 100.0 | 14.3 | 0.9 | 0.332 | I222 |
| 2.02 | 0.083 | 0.036 | 2.711 | 13.4 | 99.5 | 15.8 | 1.1 | 0.601 | F222 |
| 2.09 | 0.131 | 0.036 | 10.371 | 13.2 | 100.0 | 17.1 | 0.6 | 0.836 | P212121 |
| 1.36 | 0.069 | 0.036 | 3.593 | 23.7 | 96.8 | 32.7 | 2.8 | 0.923 | P422 |
| 2.63 | 0.226 | 0.037 | 9.377 | 6.8 | 99.7 | 6.6 | 0.3 | 0.217 | C222 |
| 2.63 | 0.226 | 0.037 | 9.377 | 6.8 | 99.7 | 6.6 | 0.3 | 0.217 | C222 |
| 1.35 | 0.126 | 0.037 | 1.647 | 3.1 | 94.3 | 7.0 | 1.0 | 0.063 | C2 |
| 4.17 | 0.142 | 0.037 | 3.094 | 7.1 | 94.3 | 7.8 | 0.8 | 0.208 | P321 |
| 2.58 | 0.144 | 0.037 | 1.826 | 3.3 | 99.1 | 8.4 | 1.3 | 0.269 | P2 |
| 2.28 | 0.105 | 0.037 | 1.197 | 4.7 | 99.8 | 8.8 | 1.1 | 0.430 | P1211 |
| 2.01 | 0.112 | 0.037 | 2.870 | 7.3 | 99.9 | 8.9 | 0.7 | 0.310 | P222 |
| 1.76 | 0.068 | 0.037 | 0.657 | 6.0 | 99.8 | 9.8 | 0.1 | 0.092 | I41 |
| 1.93 | 0.084 | 0.037 | 2.067 | 7.3 | 100.0 | 11.9 | 0.8 | 0.408 | P212121 |
| 1.57 | 0.059 | 0.037 | 0.571 | 6.1 | 100.0 | 12.3 | 0.2 | 0.127 | P1211 |
| 2.11 | 0.081 | 0.037 | 2.892 | 7.4 | 100.0 | 12.7 | 0.6 | 0.257 | I222 |
| 1.87 | 0.091 | 0.037 | 4.409 | 13.6 | 99.3 | 13.1 | 0.6 | 0.402 | F222 |
| 1.53 | 0.070 | 0.037 | 1.741 | 7.4 | 99.9 | 13.3 | 1.0 | 0.483 | I222 |
| 2.15 | 0.191 | 0.037 | 0.591 | 5.3 | 91.7 | 13.4 | 1.1 | 0.601 | P222 |
| 1.44 | 0.060 | 0.037 | 1.856 | 7.2 | 99.8 | 15.4 | 0.8 | 0.247 | I222 |
| 1.59 | 0.067 | 0.037 | 1.980 | 8.7 | 99.9 | 15.4 | 1.1 | 0.480 | P212121 |
| 2.43 | 0.158 | 0.037 | 2.728 | 13.4 | 100.0 | 16.2 | 1.1 | 0.634 | P212121 |
| 2.11 | 0.123 | 0.037 | 10.919 | 13.2 | 98.8 | 16.9 | 2.3 | 0.884 | P222 |
| 3.02 | 0.078 | 0.037 | 1.939 | 12.3 | 100.0 | 17.6 | 1.2 | 0.568 | P41212 |
| 2.93 | 0.077 | 0.037 | 10.580 | 25.3 | 100.0 | 20.3 | 0.3 | 0.149 | P222 |
| 1.31 | 0.072 | 0.037 | 4.334 | 21.6 | 97.4 | 28.1 | 0.6 | 0.498 | P42212 |
| 2.85 | 0.193 | 0.038 | 1.251 | 1.4 | 13.5 | 2.7 | 0.5 | 0.134 | C2 |
| 2.85 | 0.193 | 0.038 | 1.251 | 1.4 | 13.5 | 2.7 | 0.5 | 0.134 | C2 |
| 2.16 | 0.144 | 0.038 | 1.481 | 3.2 | 93.3 | 8.1 | 1.3 | 0.283 | P2 |
| 1.44 | 0.070 | 0.038 | 0.715 | 3.2 | 98.2 | 8.3 | 1.2 | 0.492 | P1211 |
| 1.87 | 0.092 | 0.038 | 2.806 | 7.3 | 99.9 | 10.0 | 0.7 | 0.308 | P222 |
| 1.67 | 0.090 | 0.038 | 0.690 | 6.1 | 99.3 | 11.0 | 0.5 | 0.294 | P1211 |
| 1.30 | 0.093 | 0.038 | 1.043 | 6.6 | 92.4 | 12.4 | 1.2 | 0.378 | P1211 |
| 2.79 | 0.088 | 0.038 | 4.583 | 12.6 | 100.0 | 12.8 | 0.6 | 0.244 | P422 |
| 1.85 | 0.109 | 0.038 | 2.294 | 8.8 | 100.0 | 12.8 | 0.9 | 0.371 | P212121 |
| 1.40 | 0.064 | 0.038 | 2.832 | 7.2 | 99.4 | 14.3 | 0.8 | 0.218 | I222 |
| 1.93 | 0.086 | 0.038 | 3.886 | 13.6 | 99.5 | 15.1 | 0.8 | 0.553 | F222 |
| 2.02 | 0.074 | 0.038 | 0.264 | 5.6 | 90.1 | 16.8 | 2.1 | 0.806 | P22121 |
| 3.07 | 0.073 | 0.038 | 6.942 | 25.4 | 100.0 | 24.5 | 0.6 | 0.295 | P21221 |
| 3.13 | 0.125 | 0.039 | 1.963 | 1.2 | 15.2 | 4.7 | 1.1 | 0.030 | C2 |

| | | | | | | | | | |
|---|---|---|---|---|---|---|---|---|---|
| 3.13 | 0.125 | 0.039 | 1.963 | 1.2 | 15.2 | 4.7 | 1.1 | 0.030 | C2 |
| 2.64 | 0.202 | 0.039 | 3.148 | 3.6 | 98.2 | 6.2 | 0.6 | 0.234 | P1 |
| 2.74 | 0.097 | 0.039 | 0.535 | 2.4 | 98.1 | 6.5 | 1.4 | 0.314 | P1 |
| 2.20 | 0.181 | 0.039 | 2.352 | 3.7 | 98.7 | 6.6 | 0.6 | 0.235 | P2 |
| 1.94 | 0.105 | 0.039 | 1.689 | 3.4 | 99.1 | 6.7 | 0.9 | 0.477 | C2 |
| 2.33 | 0.083 | 0.039 | 1.549 | 3.3 | 97.9 | 6.9 | 0.8 | 0.486 | P2 |
| 1.36 | 0.087 | 0.039 | 1.414 | 3.1 | 96.1 | 7.4 | 0.8 | 0.212 | P2 |
| 2.43 | 0.074 | 0.039 | 1.206 | 3.3 | 98.0 | 7.8 | 0.8 | 0.524 | P121 |
| 2.05 | 0.150 | 0.039 | 3.305 | 6.6 | 99.9 | 8.6 | 0.5 | 0.172 | P222 |
| 4.57 | 0.170 | 0.039 | 2.007 | 6.6 | 99.6 | 8.6 | 1.2 | 0.457 | P3121 |
| 2.03 | 0.066 | 0.039 | 0.585 | 7.0 | 94.3 | 9.1 | 0.3 | 0.318 | I121 |
| 1.78 | 0.099 | 0.039 | 1.476 | 4.5 | 99.9 | 9.4 | 1.1 | 0.269 | I121 |
| 1.78 | 0.099 | 0.039 | 1.476 | 4.5 | 99.9 | 9.4 | 1.1 | 0.269 | I121 |
| 2.91 | 0.078 | 0.039 | 2.481 | 6.5 | 99.9 | 10.5 | 0.7 | 0.290 | P222 |
| 1.44 | 0.078 | 0.039 | 3.841 | 7.3 | 99.5 | 10.8 | 0.6 | 0.138 | I222 |
| 1.59 | 0.102 | 0.039 | 1.392 | 6.6 | 99.5 | 11.8 | 1.2 | 0.374 | P212121 |
| 1.30 | 0.066 | 0.039 | 2.900 | 4.2 | 91.9 | 12.1 | 0.9 | 0.373 | P121 |
| 1.54 | 0.062 | 0.039 | 1.530 | 7.4 | 100.0 | 13.4 | 0.9 | 0.543 | I222 |
| 2.18 | 0.152 | 0.039 | 0.442 | 5.1 | 91.9 | 13.7 | 1.0 | 0.477 | P212121 |
| 2.33 | 0.084 | 0.039 | 3.367 | 13.2 | 99.9 | 16.7 | 0.8 | 0.422 | I222 |
| 2.33 | 0.084 | 0.039 | 3.367 | 13.2 | 99.9 | 16.7 | 0.8 | 0.422 | I222 |
| 2.43 | 0.079 | 0.039 | 2.311 | 13.3 | 100.0 | 19.4 | 1.1 | 0.589 | I222 |
| 2.43 | 0.079 | 0.039 | 2.311 | 13.3 | 100.0 | 19.4 | 1.1 | 0.589 | I222 |
| 1.53 | 0.045 | 0.039 | 1.777 | 7.3 | 99.7 | 23.1 | 1.1 | 0.588 | I222 |
| 2.45 | 0.098 | 0.040 | 1.116 | 1.1 | 16.7 | 4.3 | 0.6 | 0.068 | C2 |
| 2.45 | 0.098 | 0.040 | 1.116 | 1.1 | 16.7 | 4.3 | 0.6 | 0.068 | C2 |
| 2.20 | 0.209 | 0.040 | 1.659 | 3.7 | 99.6 | 6.9 | 0.9 | 0.417 | P2 |
| 2.09 | 0.122 | 0.040 | 2.500 | 4.5 | 99.8 | 7.3 | 0.7 | 0.247 | P212121 |
| 2.09 | 0.122 | 0.040 | 2.500 | 4.5 | 99.8 | 7.3 | 0.7 | 0.247 | P212121 |
| 1.36 | 0.054 | 0.040 | 0.650 | 3.3 | 91.6 | 9.8 | 1.1 | 0.520 | P1211 |
| 1.71 | 0.107 | 0.040 | 0.561 | 6.1 | 99.1 | 10.6 | 0.6 | 0.197 | P1211 |
| 3.39 | 0.099 | 0.040 | 6.321 | 12.4 | 99.9 | 11.3 | 0.4 | 0.095 | P422 |
| 1.67 | 0.092 | 0.040 | 0.578 | 5.9 | 99.8 | 11.6 | 0.4 | 0.321 | P1211 |
| 1.49 | 0.067 | 0.040 | 2.328 | 7.4 | 99.9 | 11.9 | 0.8 | 0.376 | I222 |
| 1.49 | 0.078 | 0.040 | 4.739 | 8.8 | 99.6 | 12.4 | 0.5 | 0.234 | P222 |
| 1.27 | 0.099 | 0.040 | 1.582 | 6.6 | 91.3 | 12.4 | 1.5 | 0.343 | P2 |
| 1.53 | 0.095 | 0.040 | 7.127 | 8.8 | 99.0 | 13.8 | 0.5 | 0.367 | P212121 |
| 1.53 | 0.083 | 0.040 | 1.934 | 8.8 | 98.5 | 14.0 | 1.1 | 0.584 | P212121 |
| 1.95 | 0.082 | 0.040 | 1.615 | 6.5 | 99.9 | 14.6 | 1.0 | 0.447 | P212121 |
| 3.34 | 0.152 | 0.040 | 5.053 | 12.2 | 99.6 | 14.7 | 0.6 | 0.168 | P41212 |
| 3.33 | 0.098 | 0.040 | 3.194 | 12.3 | 99.5 | 15.5 | 1.0 | 0.436 | P41212 |
| 2.37 | 0.113 | 0.040 | 4.079 | 25.6 | 100.0 | 19.8 | 1.0 | 0.613 | P21221 |
| 3.11 | 0.211 | 0.041 | 2.084 | 1.5 | 12.8 | 3.4 | 0.5 | 0.189 | C2 |
| 3.11 | 0.211 | 0.041 | 2.084 | 1.5 | 12.8 | 3.4 | 0.5 | 0.189 | C2 |
| 2.32 | 0.106 | 0.041 | 1.700 | 1.2 | 16.0 | 3.9 | 0.6 | 0.149 | C2 |
| 2.32 | 0.106 | 0.041 | 1.700 | 1.2 | 16.0 | 3.9 | 0.6 | 0.149 | C2 |
| 2.65 | 0.141 | 0.041 | 1.650 | 1.2 | 16.4 | 4.1 | 0.6 | 0.015 | C2 |
| 2.65 | 0.141 | 0.041 | 1.650 | 1.2 | 16.4 | 4.1 | 0.6 | 0.015 | C2 |
| 1.98 | 0.103 | 0.041 | 2.454 | 3.3 | 98.0 | 6.1 | 0.5 | 0.129 | P2 |
| 2.47 | 0.180 | 0.041 | 2.395 | 3.4 | 99.1 | 6.2 | 0.7 | 0.163 | C2 |
| 2.02 | 0.142 | 0.041 | 3.607 | 4.5 | 99.7 | 6.3 | 0.7 | 0.189 | P222 |
| 2.02 | 0.142 | 0.041 | 3.607 | 4.5 | 99.7 | 6.3 | 0.7 | 0.189 | P222 |
| 2.13 | 0.129 | 0.041 | 1.734 | 4.7 | 99.6 | 7.4 | 0.8 | 0.238 | P2 |
| 2.09 | 0.156 | 0.041 | 4.800 | 6.7 | 98.3 | 8.5 | 0.8 | 0.346 | I121 |

| | | | | | | | | | | |
|---|---|---|---|---|---|---|---|---|---|---|
| 1.73 | 0.117 | 0.041 | 2.642 | 4.5 | 99.6 | 8.8 | 1.0 | 0.194 | C2 | |
| 1.73 | 0.117 | 0.041 | 2.642 | 4.5 | 99.6 | 8.8 | 1.0 | 0.194 | C2 | |
| 1.85 | 0.109 | 0.041 | 0.568 | 5.9 | 99.9 | 9.0 | 0.3 | 0.040 | P1211 | |
| 2.06 | 0.111 | 0.041 | 1.652 | 6.4 | 99.9 | 9.7 | 1.1 | 0.439 | P212121 | |
| 1.74 | 0.100 | 0.041 | 0.630 | 6.1 | 99.3 | 10.1 | 0.3 | 0.138 | P1211 | |
| 2.06 | 0.097 | 0.041 | 1.668 | 6.5 | 100.0 | 10.9 | 1.1 | 0.404 | P212121 | |
| 3.20 | 0.098 | 0.041 | 5.483 | 12.4 | 99.9 | 13.4 | 0.7 | 0.243 | P422 | |
| 2.93 | 0.113 | 0.041 | 1.964 | 12.0 | 99.8 | 13.9 | 1.2 | 0.815 | P22121 | |
| 2.29 | 0.119 | 0.041 | 4.986 | 25.7 | 99.9 | 17.1 | 0.8 | 0.563 | P222 | |
| 1.45 | 0.089 | 0.041 | 3.242 | 27.3 | 96.1 | 25.3 | 0.7 | 0.177 | P622 | |
| 1.47 | 0.088 | 0.041 | 2.407 | 27.3 | 96.2 | 26.1 | 0.5 | 0.198 | P6122 | |
| 3.80 | 0.224 | 0.042 | 1.353 | 1.4 | 22.7 | 2.5 | 0.6 | 0.352 | C222 | |
| 3.80 | 0.224 | 0.042 | 1.353 | 1.4 | 22.7 | 2.5 | 0.6 | 0.352 | C222 | |
| 3.10 | 0.167 | 0.042 | 1.272 | 1.3 | 13.5 | 3.2 | 0.5 | 0.065 | C2 | |
| 3.10 | 0.167 | 0.042 | 1.272 | 1.3 | 13.5 | 3.2 | 0.5 | 0.065 | C2 | |
| 1.74 | 0.229 | 0.042 | 2.210 | 3.7 | 99.6 | 6.0 | 0.7 | 0.241 | P2 | |
| 1.74 | 0.229 | 0.042 | 2.210 | 3.7 | 99.6 | 6.0 | 0.7 | 0.241 | P2 | |
| 2.09 | 0.174 | 0.042 | 3.497 | 6.4 | 99.6 | 6.3 | 0.6 | 0.177 | P222 | |
| 4.19 | 0.218 | 0.042 | 4.970 | 6.7 | 99.8 | 6.4 | 0.5 | 0.166 | P321 | |
| 2.20 | 0.144 | 0.042 | 5.277 | 6.7 | 98.0 | 9.2 | 2.2 | 0.534 | C2 | |
| 2.69 | 0.132 | 0.042 | 4.801 | 12.2 | 99.7 | 10.4 | 0.5 | 0.447 | P222 | |
| 2.08 | 0.091 | 0.042 | 1.190 | 7.2 | 100.0 | 10.9 | 1.2 | 0.633 | P212121 | |
| 1.80 | 0.122 | 0.042 | 4.194 | 8.8 | 99.9 | 11.3 | 0.7 | 0.225 | P222 | |
| 2.16 | 0.142 | 0.042 | 15.904 | | 8.1 | 99.3 | 12.7 | 2.7 | 0.658 | C2 |
| 1.69 | 0.078 | 0.042 | 1.293 | 6.5 | 95.9 | 12.8 | 1.0 | 0.639 | P1211 | |
| 1.69 | 0.078 | 0.042 | 1.293 | 6.5 | 95.9 | 12.8 | 1.0 | 0.639 | P1211 | |
| 3.54 | 0.092 | 0.042 | 4.301 | 12.4 | 99.9 | 13.3 | 0.6 | 0.194 | P41212 | |
| 2.63 | 0.108 | 0.043 | 0.319 | 1.1 | 17.3 | 3.4 | 0.8 | 0.765 | P2 | |
| 3.12 | 0.190 | 0.043 | 1.675 | 1.7 | 10.9 | 3.7 | 0.6 | 0.326 | C2 | |
| 3.12 | 0.190 | 0.043 | 1.675 | 1.7 | 10.9 | 3.7 | 0.6 | 0.326 | C2 | |
| 2.82 | 0.137 | 0.043 | 0.731 | 1.4 | 14.0 | 3.8 | 0.7 | 0.432 | C2 | |
| 2.82 | 0.137 | 0.043 | 0.731 | 1.4 | 14.0 | 3.8 | 0.7 | 0.432 | C2 | |
| 2.61 | 0.058 | 0.043 | 0.368 | 1.6 | 44.2 | 4.8 | 1.2 | 0.088 | P1 | |
| 2.20 | 0.146 | 0.043 | 1.760 | 3.5 | 94.7 | 5.5 | 0.7 | 0.167 | P2 | |
| 2.63 | 0.125 | 0.043 | 6.992 | 5.6 | 99.1 | 7.1 | 0.3 | 0.080 | R3 | |
| 2.18 | 0.150 | 0.043 | 2.650 | 6.4 | 99.8 | 8.0 | 0.7 | 0.192 | P212121 | |
| 1.89 | 0.117 | 0.043 | 5.025 | 6.5 | 99.9 | 8.1 | 0.5 | 0.150 | P222 | |
| 1.93 | 0.122 | 0.043 | 3.695 | 6.6 | 99.9 | 8.2 | 0.5 | 0.129 | P222 | |
| 2.76 | 0.159 | 0.043 | 2.062 | 6.7 | 99.9 | 8.3 | 1.0 | 0.586 | F222 | |
| 1.28 | 0.061 | 0.043 | 0.884 | 3.2 | 84.6 | 9.3 | 0.6 | 0.359 | P2 | |
| 2.39 | 0.288 | 0.043 | 5.367 | 12.2 | 100.0 | 10.7 | 0.6 | 0.530 | C222 | |
| 1.43 | 0.188 | 0.043 | 1.437 | 20.0 | 100.0 | 14.5 | 2.5 | 0.297 | P3121 | |
| 2.39 | 0.154 | 0.043 | 5.341 | 25.5 | 100.0 | 15.7 | 0.7 | 0.369 | P222 | |
| 2.27 | 0.091 | 0.043 | 2.846 | 14.5 | 97.9 | 16.3 | 0.6 | 0.250 | C222 | |
| 2.20 | 0.095 | 0.043 | 3.340 | 26.3 | 98.5 | 21.3 | 1.0 | 0.630 | P21221 | |
| 3.11 | 0.230 | 0.044 | 8.050 | 1.5 | 12.7 | 2.7 | 0.1 | 0.061 | C2 | |
| 3.11 | 0.230 | 0.044 | 8.050 | 1.5 | 12.7 | 2.7 | 0.1 | 0.061 | C2 | |
| 2.85 | 0.321 | 0.044 | 2.339 | 1.8 | 97.3 | 3.8 | 0.6 | 0.109 | P1 | |
| 2.86 | 0.130 | 0.044 | 1.757 | 1.2 | 15.8 | 4.4 | 0.9 | 0.025 | C2 | |
| 2.86 | 0.130 | 0.044 | 1.757 | 1.2 | 15.8 | 4.4 | 0.9 | 0.025 | C2 | |
| 2.63 | 0.147 | 0.044 | 2.950 | 3.4 | 98.1 | 5.5 | 0.4 | 0.166 | P2 | |
| 2.47 | 0.202 | 0.044 | 4.629 | 6.7 | 99.9 | 5.6 | 0.4 | 0.277 | F222 | |
| 2.16 | 0.230 | 0.044 | 4.105 | 6.3 | 99.8 | 6.1 | 0.6 | 0.158 | P222 | |
| 1.91 | 0.137 | 0.044 | 0.708 | 6.2 | 98.9 | 6.6 | 0.1 | 0.052 | P1211 | |

| | | | | | | | | | |
|---|---|---|---|---|---|---|---|---|---|
| 2.76 | 0.121 | 0.044 | 1.663 | 3.4 | 98.1 | 6.6 | 0.7 | 0.210 | P1211 |
| 1.93 | 0.105 | 0.044 | 2.433 | 7.2 | 100.0 | 8.2 | 0.7 | 0.337 | P222 |
| 3.15 | 0.166 | 0.044 | 0.552 | 8.4 | 100.0 | 8.3 | 0.9 | 0.274 | H32 |
| 1.80 | 0.103 | 0.044 | 0.576 | 6.3 | 98.9 | 9.0 | 0.2 | 0.011 | P1211 |
| 2.00 | 0.096 | 0.044 | 2.245 | 7.5 | 100.0 | 11.4 | 0.9 | 0.358 | P212121 |
| 2.70 | 0.131 | 0.044 | 5.663 | 12.5 | 99.8 | 11.6 | 0.6 | 0.363 | P222 |
| 2.78 | 0.088 | 0.044 | 5.799 | 12.8 | 100.0 | 13.4 | 0.6 | 0.159 | P422 |
| 2.80 | 0.122 | 0.044 | 4.321 | 12.5 | 99.9 | 13.7 | 0.7 | 0.493 | P22121 |
| 2.22 | 0.122 | 0.044 | 5.955 | 25.7 | 100.0 | 16.6 | 0.7 | 0.402 | P222 |
| 2.48 | 0.145 | 0.044 | 4.380 | 25.4 | 100.0 | 18.6 | 0.9 | 0.505 | P21221 |
| 2.13 | 0.098 | 0.044 | 4.221 | 26.2 | 98.0 | 18.7 | 0.7 | 0.467 | P222 |
| 2.40 | 0.087 | 0.044 | 1.685 | 15.2 | 99.5 | 18.9 | 1.0 | 0.541 | C2221 |
| 2.29 | 0.117 | 0.044 | 4.113 | 25.6 | 100.0 | 19.2 | 0.9 | 0.546 | P21221 |
| 2.31 | 0.270 | 0.044 | 5.390 | 50.6 | 100.0 | 20.6 | 1.2 | 0.527 | P41212 |
| 3.01 | 0.082 | 0.044 | 3.638 | 12.1 | 99.9 | 23.1 | 0.7 | 0.449 | P41212 |
| 1.88 | 0.138 | 0.045 | 5.289 | 6.5 | 99.9 | 6.9 | 0.4 | 0.146 | P222 |
| 3.03 | 0.125 | 0.045 | 2.425 | 6.3 | 99.7 | 7.2 | 0.7 | 0.281 | C222 |
| 1.84 | 0.129 | 0.045 | 0.630 | 6.1 | 98.7 | 8.5 | 0.3 | 0.116 | P1211 |
| 1.92 | 0.170 | 0.045 | 1.403 | 8.1 | 99.4 | 9.3 | 1.3 | 0.325 | I121 |
| 2.01 | 0.109 | 0.045 | 2.515 | 6.6 | 100.0 | 9.8 | 0.6 | 0.224 | P212121 |
| 2.17 | 0.141 | 0.045 | 2.129 | 6.5 | 99.9 | 9.9 | 0.8 | 0.336 | P212121 |
| 2.06 | 0.105 | 0.045 | 1.883 | 6.3 | 99.6 | 10.3 | 0.8 | 0.366 | P212121 |
| 2.30 | 0.128 | 0.045 | 3.905 | 13.4 | 99.9 | 11.7 | 0.6 | 0.354 | P2 |
| 1.59 | 0.085 | 0.045 | 1.224 | 7.9 | 100.0 | 12.1 | 1.2 | 0.723 | I222 |
| 1.44 | 0.078 | 0.045 | 1.359 | 7.3 | 99.7 | 13.2 | 1.2 | 0.410 | I222 |
| 2.14 | 0.104 | 0.045 | 3.175 | 13.6 | 100.0 | 15.2 | 0.9 | 0.458 | P2 |
| 3.01 | 0.080 | 0.045 | 2.206 | 12.6 | 100.0 | 17.3 | 1.2 | 0.506 | P41212 |
| 2.37 | 0.122 | 0.045 | 3.615 | 25.4 | 100.0 | 19.1 | 1.0 | 0.549 | P21221 |
| 2.13 | 0.105 | 0.045 | 3.972 | 25.3 | 99.9 | 21.4 | 0.7 | 0.391 | P21221 |
| 3.13 | 0.209 | 0.046 | 1.437 | 1.5 | 12.2 | 3.1 | 0.7 | 0.286 | C2 |
| 3.13 | 0.209 | 0.046 | 1.437 | 1.5 | 12.2 | 3.1 | 0.7 | 0.286 | C2 |
| 1.93 | 0.131 | 0.046 | 3.666 | 7.1 | 99.4 | 7.3 | 0.5 | 0.237 | P222 |
| 2.78 | 0.105 | 0.046 | 3.475 | 6.3 | 99.7 | 7.3 | 0.6 | 0.344 | C222 |
| 1.80 | 0.110 | 0.046 | 0.629 | 5.9 | 100.0 | 9.2 | 0.3 | 0.298 | P1211 |
| 1.49 | 0.093 | 0.046 | 2.507 | 8.0 | 99.8 | 9.7 | 0.8 | 0.395 | I222 |
| 1.93 | 0.107 | 0.046 | 3.068 | 7.5 | 99.9 | 10.1 | 0.7 | 0.230 | P222 |
| 3.14 | 0.116 | 0.046 | 10.049 | 13.4 | 99.8 | 10.4 | 0.2 | 0.094 | P2 |
| 1.65 | 0.098 | 0.046 | 1.525 | 6.5 | 94.4 | 12.2 | 1.0 | 0.592 | P2 |
| 1.65 | 0.098 | 0.046 | 1.525 | 6.5 | 94.4 | 12.2 | 1.0 | 0.592 | P2 |
| 3.22 | 0.157 | 0.046 | 6.982 | 19.3 | 100.0 | 12.6 | 0.5 | 0.098 | P321 |
| 1.44 | 0.084 | 0.046 | 1.624 | 8.6 | 99.9 | 13.8 | 1.1 | 0.406 | P212121 |
| 2.80 | 0.140 | 0.046 | 3.391 | 12.5 | 99.9 | 14.1 | 1.0 | 0.532 | P22121 |
| 2.29 | 0.129 | 0.046 | 3.951 | 25.5 | 100.0 | 16.4 | 0.9 | 0.497 | P222 |
| 2.01 | 0.098 | 0.046 | 0.883 | 10.6 | 99.7 | 16.8 | 1.4 | 0.677 | P41212 |
| 2.01 | 0.098 | 0.046 | 0.883 | 10.6 | 99.7 | 16.8 | 1.4 | 0.677 | P41212 |
| 3.05 | 0.212 | 0.047 | 1.489 | 1.4 | 13.4 | 2.9 | 0.3 | 0.199 | C2 |
| 3.05 | 0.212 | 0.047 | 1.489 | 1.4 | 13.4 | 2.9 | 0.3 | 0.199 | C2 |
| 2.47 | 0.097 | 0.047 | 1.099 | 1.2 | 16.2 | 3.2 | 0.9 | 0.071 | C2 |
| 2.47 | 0.097 | 0.047 | 1.099 | 1.2 | 16.2 | 3.2 | 0.9 | 0.071 | C2 |
| 3.04 | 0.087 | 0.047 | 3.223 | 3.4 | 98.7 | 6.4 | 0.5 | 0.166 | C2 |
| 1.74 | 0.288 | 0.047 | 1.903 | 3.7 | 99.5 | 7.0 | 1.0 | 0.293 | P2 |
| 1.74 | 0.288 | 0.047 | 1.903 | 3.7 | 99.5 | 7.0 | 1.0 | 0.293 | P2 |
| 2.26 | 0.196 | 0.047 | 2.548 | 6.3 | 99.8 | 7.4 | 0.9 | 0.256 | P212121 |
| 2.86 | 0.171 | 0.047 | 1.050 | 3.3 | 99.1 | 9.6 | 2.7 | 0.286 | P1211 |

| | | | | | | | | | |
|---|---|---|---|---|---|---|---|---|---|
| 2.70 | 0.149 | 0.047 | 3.811 | 12.5 | 99.8 | 12.3 | 1.1 | 0.651 | P222 |
| 1.53 | 0.112 | 0.047 | 3.542 | 29.3 | 99.7 | 22.7 | 1.5 | 0.433 | P622 |
| 2.33 | 0.117 | 0.048 | 0.914 | 1.1 | 16.5 | 3.6 | 0.6 | 0.239 | C2 |
| 2.33 | 0.117 | 0.048 | 0.914 | 1.1 | 16.5 | 3.6 | 0.6 | 0.239 | C2 |
| 2.47 | 0.096 | 0.048 | 0.647 | 1.1 | 16.6 | 4.1 | 0.8 | 0.473 | C2 |
| 2.47 | 0.096 | 0.048 | 0.647 | 1.1 | 16.6 | 4.1 | 0.8 | 0.473 | C2 |
| 2.82 | 0.109 | 0.048 | 1.183 | 1.1 | 16.4 | 4.2 | 0.6 | 0.173 | C2 |
| 2.82 | 0.109 | 0.048 | 1.183 | 1.1 | 16.4 | 4.2 | 0.6 | 0.173 | C2 |
| 2.33 | 0.112 | 0.048 | 1.232 | 1.5 | 12.7 | 4.2 | 0.7 | 0.379 | C2 |
| 2.33 | 0.112 | 0.048 | 1.232 | 1.5 | 12.7 | 4.2 | 0.7 | 0.379 | C2 |
| 3.07 | 0.236 | 0.048 | 1.248 | 3.8 | 99.3 | 6.2 | 1.1 | 0.333 | C121 |
| 2.11 | 0.065 | 0.048 | 0.395 | 5.7 | 96.7 | 6.6 | 2.1 | 0.224 | I222 |
| 2.08 | 0.167 | 0.048 | 3.303 | 6.5 | 99.9 | 7.9 | 0.5 | 0.173 | P222 |
| 1.66 | 0.358 | 0.048 | 2.026 | 19.9 | 100.0 | 9.7 | 1.7 | 0.280 | P321 |
| 2.48 | 0.205 | 0.048 | 3.760 | 12.3 | 99.9 | 9.8 | 0.8 | 0.644 | P222 |
| 1.24 | 0.233 | 0.048 | 1.983 | 17.1 | 97.8 | 10.0 | 1.0 | 0.066 | P321 |
| 2.55 | 0.196 | 0.048 | 2.917 | 12.3 | 100.0 | 10.8 | 0.7 | 0.703 | P21221 |
| 1.94 | 0.103 | 0.048 | 2.633 | 12.2 | 100.0 | 11.6 | 0.8 | 0.371 | P422 |
| 1.94 | 0.103 | 0.048 | 2.633 | 12.2 | 100.0 | 11.6 | 0.8 | 0.371 | P422 |
| 1.59 | 0.169 | 0.048 | 3.005 | 12.7 | 98.5 | 12.2 | 0.8 | 0.271 | P212121 |
| 1.85 | 0.052 | 0.048 | 0.679 | 10.3 | 100.0 | 13.3 | 0.1 | 0.186 | P41212 |
| 1.89 | 0.112 | 0.048 | 4.828 | 11.0 | 95.7 | 13.5 | 1.5 | 0.736 | P212121 |
| 3.35 | 0.157 | 0.048 | 4.159 | 19.2 | 99.7 | 15.3 | 0.8 | 0.305 | P3121 |
| 2.88 | 0.519 | 0.048 | 94.183 | 47.3 | 93.5 | 15.9 | 2.2 | 0.523 | P422 |
| 1.53 | 0.109 | 0.048 | 2.573 | 28.7 | 99.7 | 23.0 | 1.0 | 0.355 | P6122 |
| 1.85 | 0.327 | 0.049 | 1.365 | 6.8 | 99.8 | 4.2 | 1.3 | 0.318 | I121 |
| 2.00 | 0.116 | 0.049 | 1.290 | 3.4 | 99.7 | 5.8 | 0.9 | 0.579 | C121 |
| 2.05 | 0.132 | 0.049 | 2.630 | 7.2 | 99.9 | 7.8 | 0.7 | 0.286 | P222 |
| 1.60 | 0.424 | 0.049 | 3.815 | 19.9 | 99.7 | 8.4 | 1.7 | 0.313 | P321 |
| 1.86 | 0.219 | 0.049 | 1.176 | 4.3 | 99.5 | 8.6 | 1.7 | 0.345 | I121 |
| 1.53 | 0.198 | 0.049 | 3.416 | 12.8 | 96.1 | 11.9 | 1.2 | 0.645 | P212121 |
| 2.01 | 0.100 | 0.049 | 1.873 | 12.3 | 100.0 | 13.0 | 1.0 | 0.574 | P41212 |
| 2.01 | 0.100 | 0.049 | 1.873 | 12.3 | 100.0 | 13.0 | 1.0 | 0.574 | P41212 |
| 2.33 | 0.136 | 0.050 | 1.623 | 1.3 | 14.7 | 3.9 | 0.6 | 0.117 | C2 |
| 2.33 | 0.136 | 0.050 | 1.623 | 1.3 | 14.7 | 3.9 | 0.6 | 0.117 | C2 |
| 2.46 | 0.102 | 0.050 | 1.054 | 1.2 | 16.3 | 4.0 | 0.8 | 0.139 | C2 |
| 2.46 | 0.102 | 0.050 | 1.054 | 1.2 | 16.3 | 4.0 | 0.8 | 0.139 | C2 |
| 2.62 | 0.104 | 0.050 | 0.864 | 1.1 | 16.8 | 4.3 | 0.7 | 0.342 | C2 |
| 2.62 | 0.104 | 0.050 | 0.864 | 1.1 | 16.8 | 4.3 | 0.7 | 0.342 | C2 |
| 2.33 | 0.095 | 0.050 | 0.887 | 1.5 | 12.6 | 4.9 | 0.8 | 0.347 | C2 |
| 2.33 | 0.095 | 0.050 | 0.887 | 1.5 | 12.6 | 4.9 | 0.8 | 0.347 | C2 |
| 3.12 | 0.670 | 0.050 | 6.766 | 12.9 | 95.9 | 6.7 | 0.4 | 0.069 | P422 |
| 3.12 | 0.670 | 0.050 | 6.766 | 12.9 | 95.9 | 6.7 | 0.4 | 0.069 | P422 |
| 3.33 | 0.624 | 0.050 | 5.070 | 12.8 | 99.8 | 8.5 | 0.7 | 0.157 | P42212 |
| 3.33 | 0.624 | 0.050 | 5.070 | 12.8 | 99.8 | 8.5 | 0.7 | 0.157 | P42212 |
| 2.76 | 0.110 | 0.050 | 3.707 | 5.6 | 99.0 | 8.9 | 0.5 | 0.069 | H3 |
| 2.25 | 0.111 | 0.050 | 1.310 | 6.5 | 99.9 | 9.6 | 1.1 | 0.456 | P212121 |
| 2.01 | 0.191 | 0.050 | 3.105 | 7.5 | 99.7 | 9.7 | 0.6 | 0.265 | P222 |
| 1.59 | 0.124 | 0.050 | 3.298 | 7.3 | 99.8 | 10.1 | 0.7 | 0.228 | I222 |
| 2.19 | 0.237 | 0.050 | 35.149 | 12.8 | 99.4 | 10.9 | 0.4 | 0.715 | P21212 |
| 2.76 | 0.218 | 0.050 | 5.230 | 13.0 | 99.8 | 11.9 | 0.8 | 0.568 | P41212 |
| 2.76 | 0.218 | 0.050 | 5.230 | 13.0 | 99.8 | 11.9 | 0.8 | 0.568 | P41212 |
| 2.88 | 0.107 | 0.050 | 2.148 | 12.2 | 99.8 | 13.5 | 0.9 | 0.502 | P41212 |
| 1.86 | 0.097 | 0.050 | 4.432 | 13.6 | 100.0 | 13.6 | 0.7 | 0.216 | I222 |

| | | | | | | | | | |
|---|---|---|---|---|---|---|---|---|---|
| 1.64 | 0.187 | 0.050 | 0.844 | 11.6 | 99.0 | 13.9 | 1.8 | 0.390 | P212121 |
| 2.39 | 0.152 | 0.050 | 4.908 | 25.5 | 100.0 | 15.2 | 0.7 | 0.290 | P222 |
| 1.92 | 0.092 | 0.050 | 3.046 | 13.5 | 100.0 | 15.4 | 0.9 | 0.360 | I222 |
| 2.47 | 0.146 | 0.050 | 4.504 | 25.4 | 100.0 | 17.3 | 0.8 | 0.476 | P2221 |
| 3.12 | 0.146 | 0.051 | 0.768 | 8.3 | 100.0 | 3.4 | 0.5 | 0.135 | P63 |
| 2.86 | 0.177 | 0.051 | 1.045 | 1.4 | 13.5 | 3.5 | 0.6 | 0.266 | C2 |
| 2.86 | 0.177 | 0.051 | 1.045 | 1.4 | 13.5 | 3.5 | 0.6 | 0.266 | C2 |
| 2.84 | 0.175 | 0.051 | 4.728 | 1.2 | 16.2 | 3.6 | 0.4 | 0.403 | C2 |
| 2.84 | 0.175 | 0.051 | 4.728 | 1.2 | 16.2 | 3.6 | 0.4 | 0.403 | C2 |
| 2.47 | 0.098 | 0.051 | 0.618 | 1.1 | 16.8 | 4.2 | 0.9 | 0.491 | C2 |
| 2.47 | 0.098 | 0.051 | 0.618 | 1.1 | 16.8 | 4.2 | 0.9 | 0.491 | C2 |
| 2.93 | 0.290 | 0.051 | 2.328 | 3.7 | 99.4 | 5.2 | 0.8 | 0.181 | C2 |
| 2.05 | 0.132 | 0.051 | 3.361 | 6.5 | 99.9 | 6.8 | 0.6 | 0.167 | P222 |
| 2.09 | 0.137 | 0.051 | 3.280 | 6.5 | 99.9 | 8.0 | 0.6 | 0.172 | P222 |
| 2.18 | 0.124 | 0.051 | 2.165 | 6.6 | 99.9 | 9.3 | 0.8 | 0.306 | P212121 |
| 2.02 | 0.126 | 0.051 | 6.189 | 13.4 | 100.0 | 11.5 | 0.5 | 0.183 | I222 |
| 2.02 | 0.126 | 0.051 | 6.189 | 13.4 | 100.0 | 11.5 | 0.5 | 0.183 | I222 |
| 1.71 | 0.108 | 0.051 | 1.741 | 7.3 | 100.0 | 12.5 | 1.0 | 0.399 | I222 |
| 1.56 | 0.205 | 0.051 | 1.194 | 11.1 | 95.4 | 12.6 | 0.9 | 0.260 | P222 |
| 2.09 | 0.119 | 0.051 | 4.543 | 13.3 | 100.0 | 13.2 | 0.6 | 0.224 | I222 |
| 2.09 | 0.119 | 0.051 | 4.543 | 13.3 | 100.0 | 13.2 | 0.6 | 0.224 | I222 |
| 1.35 | 0.146 | 0.051 | 0.989 | 11.3 | 97.2 | 13.4 | 1.5 | 0.339 | P42212 |
| 2.39 | 0.163 | 0.051 | 5.438 | 26.1 | 97.8 | 14.5 | 0.6 | 0.351 | P222 |
| 2.47 | 0.154 | 0.051 | 4.404 | 26.0 | 98.4 | 16.9 | 0.8 | 0.503 | P222 |
| 2.77 | 0.090 | 0.051 | 3.517 | 11.9 | 100.0 | 23.4 | 0.8 | 0.366 | P41212 |
| 2.88 | 0.277 | 0.052 | 3.528 | 3.2 | 95.3 | 4.6 | 0.5 | 0.178 | P2 |
| 2.09 | 0.347 | 0.052 | 2.827 | 4.5 | 99.7 | 5.3 | 1.4 | 0.181 | P2 |
| 1.80 | 0.115 | 0.052 | 2.203 | 3.3 | 96.0 | 6.7 | 0.8 | 0.452 | P2 |
| 2.42 | 0.239 | 0.052 | 1.778 | 6.8 | 98.6 | 7.8 | 1.1 | 0.430 | C121 |
| 2.42 | 0.239 | 0.052 | 1.778 | 6.8 | 98.6 | 7.8 | 1.1 | 0.430 | C121 |
| 1.81 | 0.291 | 0.052 | 2.285 | 4.5 | 99.3 | 8.1 | 1.7 | 0.254 | C2 |
| 1.25 | 0.065 | 0.052 | 0.502 | 4.0 | 90.7 | 11.0 | 1.6 | 0.443 | P1211 |
| 1.64 | 0.082 | 0.052 | 0.498 | 10.9 | 99.9 | 14.0 | 1.2 | 0.799 | P4212 |
| 1.64 | 0.082 | 0.052 | 0.498 | 10.9 | 99.9 | 14.0 | 1.2 | 0.799 | P4212 |
| 2.20 | 0.359 | 0.052 | 2.873 | 12.0 | 99.8 | 17.5 | 2.8 | 0.685 | P212121 |
| 1.56 | 0.439 | 0.053 | 0.910 | 6.1 | 87.4 | 3.1 | 1.0 | 0.227 | C2 |
| 3.20 | 0.166 | 0.053 | 2.665 | 1.7 | 10.4 | 3.7 | 0.4 | 0.124 | C2 |
| 3.20 | 0.166 | 0.053 | 2.665 | 1.7 | 10.4 | 3.7 | 0.4 | 0.124 | C2 |
| 2.04 | 0.142 | 0.053 | 3.397 | 6.3 | 99.7 | 6.1 | 0.5 | 0.137 | P222 |
| 2.20 | 0.186 | 0.053 | 2.326 | 6.9 | 99.6 | 6.5 | 1.0 | 0.264 | P222 |
| 2.29 | 0.159 | 0.053 | 1.736 | 6.9 | 99.7 | 8.5 | 1.1 | 0.334 | P212121 |
| 2.08 | 0.120 | 0.053 | 1.647 | 7.1 | 99.6 | 8.8 | 1.0 | 0.546 | P212121 |
| 2.05 | 0.110 | 0.053 | 2.006 | 6.5 | 100.0 | 9.5 | 0.8 | 0.284 | P212121 |
| 2.13 | 0.121 | 0.053 | 1.886 | 7.2 | 99.9 | 9.6 | 0.9 | 0.403 | P212121 |
| 2.18 | 0.215 | 0.053 | 1.944 | 7.1 | 100.0 | 10.2 | 1.0 | 0.502 | P212121 |
| 2.08 | 0.167 | 0.053 | 2.313 | 7.3 | 99.8 | 10.9 | 0.8 | 0.358 | P212121 |
| 1.80 | 0.177 | 0.053 | 1.893 | 8.5 | 99.5 | 11.0 | 1.6 | 0.283 | P22121 |
| 1.56 | 0.086 | 0.053 | 0.546 | 10.2 | 97.4 | 12.4 | 0.8 | 0.613 | P422 |
| 1.56 | 0.086 | 0.053 | 0.546 | 10.2 | 97.4 | 12.4 | 0.8 | 0.613 | P422 |
| 1.80 | 0.140 | 0.053 | 3.213 | 12.3 | 95.3 | 14.9 | 1.1 | 0.678 | P1211 |
| 1.49 | 0.294 | 0.053 | 3.708 | 23.8 | 98.3 | 15.9 | 1.4 | 0.711 | P212121 |
| 3.66 | 0.142 | 0.054 | 1.847 | 1.7 | 10.8 | 3.3 | 0.7 | 0.394 | C2 |
| 3.66 | 0.142 | 0.054 | 1.847 | 1.7 | 10.8 | 3.3 | 0.7 | 0.394 | C2 |
| 2.45 | 0.103 | 0.054 | 0.988 | 1.2 | 16.3 | 4.0 | 0.6 | 0.146 | C2 |

| | | | | | | | | | |
|---|---|---|---|---|---|---|---|---|---|
| 2.45 | 0.103 | 0.054 | 0.988 | 1.2 | 16.3 | 4.0 | 0.6 | 0.146 | C2 |
| 1.97 | 0.122 | 0.054 | 3.058 | 6.5 | 99.9 | 8.1 | 0.6 | 0.192 | P222 |
| 2.29 | 0.177 | 0.054 | 1.852 | 6.9 | 99.9 | 10.6 | 1.1 | 0.471 | P212121 |
| 3.12 | 0.408 | 0.054 | 5.331 | 45.8 | 98.5 | 18.5 | 1.6 | 0.764 | P41212 |
| 2.88 | 0.091 | 0.054 | 3.039 | 11.6 | 100.0 | 23.5 | 0.8 | 0.465 | P41212 |
| 3.07 | 0.233 | 0.055 | 2.401 | 1.7 | 11.0 | 2.5 | 0.4 | 0.127 | C2 |
| 3.07 | 0.233 | 0.055 | 2.401 | 1.7 | 11.0 | 2.5 | 0.4 | 0.127 | C2 |
| 3.08 | 0.151 | 0.055 | 0.784 | 3.0 | 99.5 | 2.6 | 0.1 | 0.489 | C121 |
| 2.20 | 0.329 | 0.055 | 3.803 | 6.9 | 98.4 | 5.7 | 0.6 | 0.240 | C2 |
| 2.20 | 0.329 | 0.055 | 3.803 | 6.9 | 98.4 | 5.7 | 0.6 | 0.240 | C2 |
| 2.65 | 0.485 | 0.055 | 5.413 | 6.5 | 93.3 | 5.9 | 1.9 | 0.411 | C2 |
| 2.21 | 0.139 | 0.055 | 4.885 | 4.4 | 98.5 | 6.5 | 0.8 | 0.297 | P2 |
| 2.13 | 0.124 | 0.055 | 2.115 | 6.3 | 99.8 | 8.0 | 0.7 | 0.323 | P212121 |
| 3.23 | 0.382 | 0.056 | 10.579 | | 4.7 | 98.4 | 3.3 | 0.1 | 0.016 C2 |
| 2.45 | 0.110 | 0.056 | 1.073 | 1.1 | 16.7 | 4.2 | 0.5 | 0.184 | C2 |
| 2.45 | 0.110 | 0.056 | 1.073 | 1.1 | 16.7 | 4.2 | 0.5 | 0.184 | C2 |
| 2.84 | 0.110 | 0.056 | 1.059 | 1.2 | 16.1 | 4.3 | 0.7 | 0.417 | C2 |
| 2.84 | 0.110 | 0.056 | 1.059 | 1.2 | 16.1 | 4.3 | 0.7 | 0.417 | C2 |
| 3.04 | 0.099 | 0.056 | 2.313 | 3.2 | 96.0 | 5.3 | 0.5 | 0.128 | C2 |
| 2.20 | 0.282 | 0.056 | 2.426 | 6.8 | 99.8 | 5.7 | 0.8 | 0.387 | I121 |
| 2.17 | 0.170 | 0.056 | 2.285 | 6.5 | 98.2 | 8.3 | 0.7 | 0.204 | P212121 |
| 2.24 | 0.110 | 0.056 | 2.345 | 5.6 | 99.7 | 9.3 | 1.1 | 0.554 | H3 |
| 2.32 | 0.100 | 0.056 | 0.956 | 5.7 | 99.6 | 10.3 | 2.3 | 0.213 | H3 |
| 2.83 | 0.295 | 0.056 | 4.910 | 12.5 | 98.4 | 11.1 | 1.1 | 0.514 | P222 |
| 2.87 | 0.116 | 0.056 | 3.081 | 12.1 | 100.0 | 12.3 | 0.8 | 0.538 | P41212 |
| 3.15 | 0.142 | 0.056 | 2.027 | 12.0 | 99.8 | 13.1 | 1.1 | 0.528 | P41212 |
| 2.21 | 0.114 | 0.056 | 3.881 | 25.8 | 99.9 | 16.7 | 1.0 | 0.530 | P222 |
| 2.28 | 0.111 | 0.056 | 3.187 | 25.7 | 100.0 | 18.7 | 1.2 | 0.617 | P21221 |
| 2.75 | 0.096 | 0.056 | 3.196 | 11.8 | 100.0 | 19.4 | 0.7 | 0.298 | P41212 |
| 1.94 | 0.136 | 0.057 | 1.710 | 3.4 | 99.5 | 5.4 | 0.8 | 0.537 | C2 |
| 2.45 | 0.225 | 0.057 | 3.578 | 7.0 | 99.5 | 6.3 | 0.6 | 0.209 | P222 |
| 2.30 | 0.119 | 0.057 | 2.128 | 4.4 | 99.0 | 7.4 | 1.0 | 0.208 | P1211 |
| 1.95 | 0.105 | 0.057 | 2.504 | 6.6 | 100.0 | 8.5 | 0.7 | 0.297 | P212121 |
| 1.80 | 0.113 | 0.057 | 0.614 | 6.1 | 98.8 | 8.6 | 0.2 | 0.194 | P1211 |
| 2.05 | 0.104 | 0.057 | 2.377 | 6.5 | 99.9 | 9.3 | 0.7 | 0.216 | P212121 |
| 2.11 | 0.150 | 0.057 | 5.748 | 13.3 | 100.0 | 10.2 | 0.5 | 0.134 | I222 |
| 1.65 | 0.178 | 0.057 | 1.683 | 8.5 | 99.9 | 10.3 | 1.4 | 0.286 | P212121 |
| 1.23 | 0.073 | 0.057 | 0.589 | 4.0 | 90.4 | 11.2 | 1.9 | 0.591 | P2 |
| 1.68 | 0.097 | 0.057 | 2.164 | 11.5 | 99.3 | 11.3 | 0.8 | 0.601 | P222 |
| 1.68 | 0.097 | 0.057 | 2.164 | 11.5 | 99.3 | 11.3 | 0.8 | 0.601 | P222 |
| 1.72 | 0.096 | 0.057 | 1.583 | 11.6 | 99.9 | 12.5 | 0.8 | 0.679 | P21221 |
| 1.72 | 0.096 | 0.057 | 1.583 | 11.6 | 99.9 | 12.5 | 0.8 | 0.679 | P21221 |
| 2.46 | 0.168 | 0.058 | 2.501 | 3.3 | 98.0 | 4.4 | 0.5 | 0.480 | P2 |
| 2.21 | 0.335 | 0.058 | 3.782 | 6.8 | 99.0 | 5.8 | 1.3 | 0.415 | C2 |
| 2.16 | 0.146 | 0.058 | 3.966 | 6.5 | 99.9 | 7.0 | 0.5 | 0.115 | P222 |
| 1.97 | 0.100 | 0.058 | 3.102 | 6.7 | 98.5 | 9.9 | 0.6 | 0.218 | P222 |
| 1.73 | 0.163 | 0.058 | 1.424 | 8.6 | 100.0 | 11.1 | 1.7 | 0.355 | P212121 |
| 1.87 | 0.130 | 0.058 | 3.990 | 24.7 | 100.0 | 12.7 | 1.0 | 0.601 | P422 |
| 1.87 | 0.130 | 0.058 | 3.990 | 24.7 | 100.0 | 12.7 | 1.0 | 0.601 | P422 |
| 1.42 | 0.181 | 0.058 | 1.275 | 61.3 | 96.8 | 26.9 | 1.4 | 0.388 | F432 |
| 3.11 | 0.113 | 0.059 | 2.449 | 1.2 | 15.0 | 3.0 | 0.6 | 0.083 | C2 |
| 3.11 | 0.113 | 0.059 | 2.449 | 1.2 | 15.0 | 3.0 | 0.6 | 0.083 | C2 |
| 2.33 | 0.097 | 0.059 | 0.764 | 1.1 | 16.1 | 3.0 | 0.7 | 0.522 | C2 |
| 2.33 | 0.097 | 0.059 | 0.764 | 1.1 | 16.1 | 3.0 | 0.7 | 0.522 | C2 |

| | | | | | | | | | |
|---|---|---|---|---|---|---|---|---|---|
| 2.64 | 0.139 | 0.059 | 1.111 | 1.2 | 16.5 | 3.6 | 0.7 | 0.198 | C2 |
| 2.64 | 0.139 | 0.059 | 1.111 | 1.2 | 16.5 | 3.6 | 0.7 | 0.198 | C2 |
| 3.49 | 0.268 | 0.059 | 5.561 | 4.6 | 98.8 | 4.1 | 0.3 | 0.008 | C121 |
| 2.57 | 0.147 | 0.059 | 1.395 | 3.3 | 98.3 | 5.3 | 0.6 | 0.506 | P1211 |
| 2.08 | 0.151 | 0.059 | 3.034 | 6.1 | 99.6 | 6.2 | 0.6 | 0.136 | P222 |
| 1.88 | 0.115 | 0.059 | 3.939 | 6.6 | 99.9 | 7.5 | 0.6 | 0.186 | P222 |
| 1.81 | 0.143 | 0.059 | 0.573 | 6.0 | 98.9 | 9.1 | 2.4 | 0.868 | P1211 |
| 2.80 | 0.278 | 0.059 | 4.537 | 12.3 | 99.1 | 10.8 | 0.8 | 0.383 | P222 |
| 1.53 | 0.087 | 0.059 | 3.276 | 8.1 | 99.8 | 11.1 | 0.8 | 0.338 | I222 |
| 1.76 | 0.160 | 0.059 | 0.886 | 7.7 | 92.8 | 11.4 | 1.8 | 0.364 | I121 |
| 1.58 | 0.082 | 0.059 | 2.076 | 8.1 | 100.0 | 12.1 | 0.8 | 0.442 | I222 |
| 1.74 | 0.135 | 0.059 | 1.708 | 12.8 | 99.6 | 12.1 | 1.1 | 0.362 | P212121 |
| 1.74 | 0.135 | 0.059 | 1.708 | 12.8 | 99.6 | 12.1 | 1.1 | 0.362 | P212121 |
| 2.83 | 0.147 | 0.060 | 0.839 | 1.4 | 14.0 | 3.5 | 0.8 | 0.362 | C2 |
| 2.83 | 0.147 | 0.060 | 0.839 | 1.4 | 14.0 | 3.5 | 0.8 | 0.362 | C2 |
| 1.95 | 0.092 | 0.060 | 0.711 | 3.2 | 94.5 | 3.7 | 0.1 | 0.118 | I121 |
| 3.48 | 0.450 | 0.060 | 2.446 | 3.7 | 99.4 | 4.4 | 1.1 | 0.201 | C2 |
| 2.29 | 0.271 | 0.060 | 1.234 | 2.8 | 95.0 | 4.4 | 1.3 | 0.097 | C121 |
| 1.79 | 0.124 | 0.060 | 1.188 | 4.6 | 99.6 | 5.5 | 0.9 | 0.517 | C121 |
| 2.64 | 0.468 | 0.060 | 2.411 | 3.4 | 98.9 | 6.1 | 1.0 | 0.192 | P2 |
| 1.53 | 0.113 | 0.060 | 1.753 | 4.4 | 99.1 | 6.4 | 0.9 | 0.268 | P2 |
| 2.55 | 0.202 | 0.060 | 2.567 | 6.9 | 99.8 | 7.8 | 0.7 | 0.245 | P212121 |
| 1.97 | 0.113 | 0.060 | 3.693 | 6.5 | 99.9 | 8.2 | 0.5 | 0.146 | P222 |
| 2.40 | 0.121 | 0.060 | 1.484 | 6.6 | 100.0 | 9.5 | 1.2 | 0.459 | P212121 |
| 2.05 | 0.092 | 0.060 | 2.069 | 6.7 | 98.8 | 10.6 | 0.9 | 0.313 | P22121 |
| 2.06 | 0.214 | 0.060 | 1.173 | 5.7 | 99.8 | 12.7 | 1.7 | 0.231 | P212121 |
| 2.38 | 0.135 | 0.060 | 3.803 | 25.7 | 100.0 | 19.0 | 1.0 | 0.577 | P21221 |
| 2.46 | 0.551 | 0.061 | 2.659 | 19.2 | 99.7 | 9.1 | 1.3 | 0.373 | P3121 |
| 1.65 | 0.142 | 0.061 | 2.954 | 12.2 | 98.2 | 10.6 | 0.7 | 0.184 | P222 |
| 1.65 | 0.142 | 0.061 | 2.954 | 12.2 | 98.2 | 10.6 | 0.7 | 0.184 | P222 |
| 1.68 | 0.176 | 0.061 | 1.090 | 7.5 | 87.8 | 11.0 | 1.2 | 0.177 | C2 |
| 1.92 | 0.131 | 0.061 | 2.951 | 24.7 | 100.0 | 13.6 | 1.1 | 0.622 | P41212 |
| 1.92 | 0.131 | 0.061 | 2.951 | 24.7 | 100.0 | 13.6 | 1.1 | 0.622 | P41212 |
| 2.30 | 0.131 | 0.061 | 2.480 | 13.1 | 100.0 | 13.8 | 1.0 | 0.402 | I222 |
| 3.15 | 0.139 | 0.062 | 4.584 | 1.7 | 11.0 | 3.5 | 0.3 | 0.158 | C2 |
| 3.15 | 0.139 | 0.062 | 4.584 | 1.7 | 11.0 | 3.5 | 0.3 | 0.158 | C2 |
| 2.11 | 0.130 | 0.062 | 0.560 | 2.7 | 93.4 | 4.0 | 0.1 | 0.182 | C121 |
| 1.93 | 0.202 | 0.062 | 6.734 | 7.2 | 99.9 | 5.8 | 0.6 | 0.195 | P222 |
| 2.15 | 0.612 | 0.062 | 2.174 | 19.8 | 100.0 | 6.7 | 2.2 | 0.209 | P3121 |
| 1.59 | 0.150 | 0.062 | 2.563 | 13.2 | 100.0 | 12.0 | 1.0 | 0.367 | P212121 |
| 3.62 | 0.173 | 0.063 | 0.852 | 1.4 | 12.8 | 2.5 | 1.2 | 0.328 | C2 |
| 3.62 | 0.173 | 0.063 | 0.852 | 1.4 | 12.8 | 2.5 | 1.2 | 0.328 | C2 |
| 2.64 | 0.158 | 0.063 | 1.233 | 1.4 | 13.7 | 3.2 | 0.6 | 0.274 | C2 |
| 2.64 | 0.158 | 0.063 | 1.233 | 1.4 | 13.7 | 3.2 | 0.6 | 0.274 | C2 |
| 2.64 | 0.444 | 0.063 | 4.779 | 6.8 | 99.9 | 5.5 | 0.4 | 0.152 | F222 |
| 1.96 | 0.302 | 0.063 | 2.227 | 4.4 | 99.5 | 5.7 | 2.1 | 0.301 | C2 |
| 1.96 | 0.302 | 0.063 | 2.227 | 4.4 | 99.5 | 5.7 | 2.1 | 0.301 | C2 |
| 2.77 | 0.354 | 0.063 | 4.494 | 6.7 | 99.9 | 6.6 | 0.5 | 0.285 | F222 |
| 1.62 | 0.600 | 0.063 | 3.365 | 6.2 | 96.6 | 6.8 | 0.7 | 0.277 | P2 |
| 2.26 | 0.591 | 0.063 | 5.102 | 19.6 | 99.8 | 7.1 | 2.8 | 0.522 | P321 |
| 2.16 | 0.111 | 0.063 | 2.123 | 6.2 | 99.8 | 8.1 | 0.8 | 0.281 | P222 |
| 1.26 | 0.333 | 0.063 | 0.805 | 17.1 | 97.1 | 8.5 | 1.4 | 0.301 | P321 |
| 1.53 | 0.112 | 0.063 | 2.084 | 8.9 | 98.8 | 11.2 | 1.1 | 0.314 | P212121 |
| 2.39 | 0.132 | 0.063 | 3.330 | 15.2 | 99.6 | 11.3 | 0.7 | 0.271 | C222 |

| | | | | | | | | | | |
|---|---|---|---|---|---|---|---|---|---|---|
| 1.30 | 0.044 | 0.063 | 1.726 | 3.2 | 88.0 | 13.4 | 0.8 | 0.490 | I121 | |
| 2.54 | 0.125 | 0.063 | 1.813 | 15.8 | 100.0 | 13.7 | 1.0 | 0.609 | C2221 | |
| 2.02 | 0.162 | 0.063 | 4.825 | 12.6 | 99.9 | 14.6 | 2.3 | 0.871 | P212121 | |
| 1.74 | 0.144 | 0.064 | 1.853 | 4.5 | 99.3 | 5.1 | 0.8 | 0.384 | C2 | |
| 1.78 | 0.182 | 0.064 | 1.917 | 4.5 | 97.9 | 5.8 | 1.0 | 0.353 | I121 | |
| 1.78 | 0.182 | 0.064 | 1.917 | 4.5 | 97.9 | 5.8 | 1.0 | 0.353 | I121 | |
| 2.20 | 0.272 | 0.064 | 1.954 | 4.4 | 99.8 | 5.9 | 1.4 | 0.262 | P1211 | |
| 2.09 | 0.180 | 0.064 | 1.873 | 7.0 | 99.9 | 6.7 | 1.0 | 0.483 | P212121 | |
| 2.17 | 0.137 | 0.064 | 2.031 | 6.2 | 99.7 | 7.6 | 0.7 | 0.240 | P21221 | |
| 3.10 | 0.559 | 0.064 | 5.230 | 13.0 | 99.9 | 7.7 | 1.0 | 0.358 | P222 | |
| 2.34 | 0.286 | 0.064 | 3.965 | 13.1 | 99.9 | 8.0 | 1.0 | 0.490 | P222 | |
| 1.70 | 0.110 | 0.064 | 1.731 | 9.7 | 99.5 | 9.4 | 0.6 | 0.309 | P422 | |
| 1.49 | 0.168 | 0.064 | 4.726 | 13.3 | 100.0 | 9.6 | 0.6 | 0.206 | P222 | |
| 1.85 | 0.356 | 0.064 | 6.499 | 20.0 | 100.0 | 10.7 | 2.3 | 0.302 | P321 | |
| 1.80 | 0.107 | 0.064 | 1.115 | 10.3 | 100.0 | 10.9 | 0.9 | 0.633 | P41212 | |
| 1.57 | 0.141 | 0.064 | 2.765 | 8.5 | 98.0 | 12.1 | 1.3 | 0.222 | P222 | |
| 1.57 | 0.141 | 0.064 | 2.765 | 8.5 | 98.0 | 12.1 | 1.3 | 0.222 | P222 | |
| 1.62 | 0.129 | 0.064 | 1.526 | 8.4 | 99.1 | 12.8 | 1.4 | 0.350 | P212121 | |
| 1.62 | 0.129 | 0.064 | 1.526 | 8.4 | 99.1 | 12.8 | 1.4 | 0.350 | P212121 | |
| 2.00 | 0.210 | 0.064 | 34.319 | | 11.5 | 100.0 | 13.2 | 1.4 | 0.755 | P41212 |
| 3.02 | 0.117 | 0.064 | 2.014 | 12.1 | 100.0 | 16.7 | 1.2 | 0.558 | P42212 | |
| 2.46 | 0.136 | 0.065 | 1.057 | 1.3 | 14.8 | 4.0 | 0.8 | 0.351 | C2 | |
| 2.46 | 0.136 | 0.065 | 1.057 | 1.3 | 14.8 | 4.0 | 0.8 | 0.351 | C2 | |
| 2.66 | 0.261 | 0.065 | 10.961 | | 13.4 | 100.0 | 7.4 | 0.2 | 0.053 | I222 |
| 2.66 | 0.261 | 0.065 | 10.961 | | 13.4 | 100.0 | 7.4 | 0.2 | 0.053 | I222 |
| 1.24 | 0.399 | 0.065 | 1.314 | 17.1 | 96.8 | 8.3 | 1.5 | 0.282 | P321 | |
| 1.52 | 0.119 | 0.065 | 2.208 | 8.3 | 99.4 | 9.1 | 0.8 | 0.339 | P212121 | |
| 2.01 | 0.308 | 0.065 | 6.387 | 12.4 | 99.2 | 10.6 | 1.3 | 0.813 | P212121 | |
| 1.52 | 0.167 | 0.065 | 4.317 | 70.0 | 99.7 | 32.7 | 1.6 | 0.817 | F432 | |
| 2.64 | 0.160 | 0.066 | 1.099 | 1.3 | 14.1 | 3.0 | 0.6 | 0.043 | C2 | |
| 2.64 | 0.160 | 0.066 | 1.099 | 1.3 | 14.1 | 3.0 | 0.6 | 0.043 | C2 | |
| 3.11 | 0.154 | 0.066 | 1.071 | 1.4 | 13.7 | 3.2 | 0.5 | 0.303 | C2 | |
| 3.11 | 0.154 | 0.066 | 1.071 | 1.4 | 13.7 | 3.2 | 0.5 | 0.303 | C2 | |
| 3.13 | 0.181 | 0.066 | 1.243 | 1.4 | 13.6 | 3.7 | 0.8 | 0.002 | C2 | |
| 3.13 | 0.181 | 0.066 | 1.243 | 1.4 | 13.6 | 3.7 | 0.8 | 0.002 | C2 | |
| 2.64 | 0.327 | 0.066 | 1.100 | 3.0 | 94.5 | 4.1 | 2.5 | 0.233 | C2 | |
| 1.73 | 0.221 | 0.066 | 4.145 | 4.6 | 97.2 | 5.5 | 1.0 | 0.265 | C2 | |
| 1.73 | 0.221 | 0.066 | 4.145 | 4.6 | 97.2 | 5.5 | 1.0 | 0.265 | C2 | |
| 1.95 | 0.104 | 0.066 | 2.155 | 6.6 | 100.0 | 9.0 | 0.8 | 0.342 | P212121 | |
| 2.07 | 0.383 | 0.066 | 4.221 | 12.6 | 99.9 | 10.3 | 1.9 | 0.807 | P212121 | |
| 1.72 | 0.919 | 0.067 | 2.213 | 6.8 | 98.4 | 2.8 | 1.0 | 0.322 | C121 | |
| 2.01 | 0.170 | 0.067 | 2.309 | 3.3 | 98.5 | 3.8 | 0.9 | 0.242 | P2 | |
| 1.98 | 0.155 | 0.067 | 2.712 | 6.5 | 99.9 | 6.7 | 0.7 | 0.211 | P222 | |
| 2.06 | 0.140 | 0.067 | 1.841 | 6.5 | 100.0 | 8.0 | 0.9 | 0.327 | P212121 | |
| 1.88 | 0.112 | 0.067 | 3.229 | 6.6 | 100.0 | 8.3 | 0.6 | 0.237 | P222 | |
| 1.79 | 0.128 | 0.067 | 2.273 | 6.2 | 98.7 | 9.4 | 0.9 | 0.605 | C2221 | |
| 1.71 | 0.152 | 0.067 | 7.158 | 12.4 | 99.8 | 14.9 | 1.0 | 0.805 | P41212 | |
| 3.29 | 0.152 | 0.067 | 7.575 | 25.2 | 99.2 | 16.8 | 0.5 | 0.254 | P21221 | |
| 2.32 | 0.174 | 0.068 | 1.295 | 3.7 | 96.5 | 4.5 | 1.2 | 0.283 | P2 | |
| 2.58 | 0.313 | 0.068 | 4.213 | 13.3 | 100.0 | 9.8 | 0.7 | 0.235 | P22121 | |
| 2.58 | 0.313 | 0.068 | 4.213 | 13.3 | 100.0 | 9.8 | 0.7 | 0.235 | P22121 | |
| 3.44 | 0.131 | 0.068 | 3.170 | 16.2 | 99.9 | 9.9 | 0.8 | 0.749 | P422 | |
| 2.43 | 0.228 | 0.068 | 2.990 | 9.5 | 99.9 | 11.0 | 0.9 | 0.303 | P3121 | |
| 2.28 | 0.358 | 0.068 | 10.264 | | 11.4 | 99.3 | 13.4 | 2.1 | 0.792 | P1211 |

| | | | | | | | | | | |
|---|---|---|---|---|---|---|---|---|---|---|
| 2.28 | 0.269 | 0.068 | 4.198 | 11.9 | 98.4 | 13.8 | 1.4 | 0.682 | P212121 | |
| 3.11 | 0.447 | 0.068 | 22.112 | | 46.6 | 99.9 | 15.4 | 0.6 | 0.792 | P4212 |
| 1.67 | 0.149 | 0.068 | 1.077 | 11.6 | 85.0 | 15.5 | 0.7 | 0.325 | P1211 | |
| 1.42 | 0.237 | 0.068 | 1.368 | 58.0 | 93.9 | 23.5 | 1.2 | 0.418 | F432 | |
| 2.85 | 0.183 | 0.069 | 0.634 | 1.8 | 97.7 | 1.9 | 0.8 | 0.581 | P1 | |
| 2.87 | 0.184 | 0.069 | 2.800 | 1.4 | 13.3 | 2.8 | 0.4 | 0.131 | C2 | |
| 2.87 | 0.184 | 0.069 | 2.800 | 1.4 | 13.3 | 2.8 | 0.4 | 0.131 | C2 | |
| 2.18 | 0.134 | 0.069 | 1.199 | 3.3 | 98.5 | 4.8 | 1.1 | 0.534 | P1211 | |
| 3.77 | 0.295 | 0.069 | 1.172 | 3.6 | 99.4 | 5.4 | 1.7 | 0.339 | C121 | |
| 1.81 | 0.153 | 0.069 | 2.555 | 6.4 | 99.6 | 6.0 | 0.9 | 0.332 | P222 | |
| 2.47 | 0.341 | 0.069 | 5.362 | 13.3 | 100.0 | 7.9 | 0.4 | 0.136 | P222 | |
| 2.47 | 0.341 | 0.069 | 5.362 | 13.3 | 100.0 | 7.9 | 0.4 | 0.136 | P222 | |
| 2.45 | 0.498 | 0.069 | 9.489 | 18.4 | 98.9 | 8.9 | 2.3 | 0.703 | P321 | |
| 1.65 | 0.171 | 0.069 | 1.582 | 11.9 | 84.1 | 16.2 | 0.9 | 0.299 | P2 | |
| 2.64 | 0.144 | 0.070 | 1.207 | 1.4 | 14.0 | 3.1 | 0.7 | 0.187 | C2 | |
| 2.64 | 0.144 | 0.070 | 1.207 | 1.4 | 14.0 | 3.1 | 0.7 | 0.187 | C2 | |
| 1.82 | 0.335 | 0.070 | 1.703 | 19.9 | 100.0 | 10.7 | 2.4 | 0.227 | P3121 | |
| 3.15 | 0.194 | 0.071 | 2.833 | 1.5 | 12.0 | 3.0 | 0.3 | 0.254 | C2 | |
| 3.15 | 0.194 | 0.071 | 2.833 | 1.5 | 12.0 | 3.0 | 0.3 | 0.254 | C2 | |
| 1.86 | 0.346 | 0.071 | 1.920 | 3.4 | 98.7 | 3.5 | 0.9 | 0.239 | C2 | |
| 2.47 | 0.154 | 0.071 | 2.397 | 3.8 | 95.9 | 3.8 | 0.6 | 0.407 | P2 | |
| 2.26 | 0.104 | 0.071 | 1.467 | 6.3 | 99.9 | 9.5 | 1.0 | 0.369 | P21221 | |
| 3.10 | 0.110 | 0.072 | 0.666 | 1.1 | 72.3 | 1.1 | 0.4 | 0.040 | P121 | |
| 2.84 | 0.141 | 0.072 | 0.890 | 1.4 | 13.7 | 3.4 | 0.6 | 0.437 | C2 | |
| 2.84 | 0.141 | 0.072 | 0.890 | 1.4 | 13.7 | 3.4 | 0.6 | 0.437 | C2 | |
| 1.91 | 0.232 | 0.072 | 4.484 | 6.7 | 99.8 | 5.4 | 0.8 | 0.149 | C2 | |
| 2.43 | 0.263 | 0.072 | 2.622 | 13.1 | 100.0 | 9.1 | 1.1 | 0.518 | P212121 | |
| 3.55 | 0.136 | 0.072 | 2.372 | 16.2 | 99.6 | 11.1 | 0.9 | 0.824 | P41212 | |
| 3.15 | 0.143 | 0.072 | 1.943 | 11.3 | 99.4 | 11.7 | 1.1 | 0.517 | P41212 | |
| 4.07 | 0.286 | 0.073 | 0.590 | 5.8 | 100.0 | 1.3 | 0.7 | 0.008 | P212121 | |
| 2.86 | 0.227 | 0.073 | 3.903 | 1.4 | 13.6 | 2.9 | 0.3 | 0.098 | C2 | |
| 2.86 | 0.227 | 0.073 | 3.903 | 1.4 | 13.6 | 2.9 | 0.3 | 0.098 | C2 | |
| 2.09 | 0.173 | 0.073 | 3.668 | 8.5 | 99.9 | 6.6 | 0.9 | 0.261 | P222 | |
| 1.69 | 0.349 | 0.073 | 1.692 | 12.2 | 99.1 | 8.9 | 1.5 | 0.376 | P21221 | |
| 2.01 | 0.098 | 0.073 | 8.711 | 8.2 | 99.9 | 13.6 | 1.0 | 0.828 | P1211 | |
| 3.17 | 0.245 | 0.074 | 2.564 | 1.4 | 12.9 | 2.0 | 0.2 | 0.005 | C2 | |
| 3.17 | 0.245 | 0.074 | 2.564 | 1.4 | 12.9 | 2.0 | 0.2 | 0.005 | C2 | |
| 2.99 | 0.152 | 0.074 | 0.265 | 2.3 | 96.8 | 2.2 | 1.0 | 0.530 | P1 | |
| 3.30 | 0.187 | 0.074 | 5.721 | 32.5 | 99.9 | 8.8 | 0.8 | 0.882 | P622 | |
| 2.80 | 0.242 | 0.074 | 6.498 | 13.3 | 100.0 | 9.0 | 0.5 | 0.206 | I222 | |
| 2.80 | 0.242 | 0.074 | 6.498 | 13.3 | 100.0 | 9.0 | 0.5 | 0.206 | I222 | |
| 1.70 | 0.104 | 0.074 | 0.594 | 12.0 | 87.7 | 18.4 | 2.4 | 0.702 | P1211 | |
| 3.11 | 0.434 | 0.075 | 5.015 | 6.8 | 99.8 | 3.6 | 0.4 | 0.188 | C2 | |
| 3.11 | 0.434 | 0.075 | 5.015 | 6.8 | 99.8 | 3.6 | 0.4 | 0.188 | C2 | |
| 1.83 | 0.521 | 0.075 | 0.364 | 3.4 | 73.1 | 4.2 | 2.4 | 0.729 | P1 | |
| 1.67 | 0.708 | 0.075 | 1.677 | 13.7 | 95.0 | 4.3 | 1.1 | 0.091 | C222 | |
| 2.13 | 0.139 | 0.075 | 1.396 | 6.1 | 99.8 | 7.9 | 1.0 | 0.474 | P212121 | |
| 2.77 | 0.464 | 0.075 | 1.564 | 15.7 | 100.0 | 8.7 | 1.9 | 0.167 | C2221 | |
| 3.16 | 0.148 | 0.075 | 2.307 | 12.0 | 99.5 | 13.2 | 1.1 | 0.571 | P41212 | |
| 1.67 | 0.108 | 0.075 | 0.996 | 12.1 | 87.7 | 18.6 | 2.8 | 0.697 | P2 | |
| 3.05 | 0.328 | 0.076 | 3.320 | 1.4 | 13.6 | 1.9 | 0.3 | 0.161 | C2 | |
| 3.05 | 0.328 | 0.076 | 3.320 | 1.4 | 13.6 | 1.9 | 0.3 | 0.161 | C2 | |
| 2.31 | 0.323 | 0.076 | 5.870 | 7.0 | 99.8 | 5.2 | 0.5 | 0.169 | P222 | |
| 1.93 | 0.262 | 0.076 | 1.404 | 4.4 | 99.7 | 5.3 | 1.4 | 0.263 | I121 | |

| | | | | | | | | | |
|---|---|---|---|---|---|---|---|---|---|
| 1.93 | 0.262 | 0.076 | 1.404 | 4.4 | 99.7 | 5.3 | 1.4 | 0.263 | I121 |
| 2.53 | 0.532 | 0.076 | 4.533 | 15.0 | 96.5 | 6.0 | 1.4 | 0.470 | P222 |
| 2.43 | 0.122 | 0.077 | 0.683 | 6.6 | 97.8 | 3.2 | 0.6 | 0.035 | P4212 |
| 2.33 | 0.303 | 0.077 | 3.992 | 6.8 | 99.9 | 4.8 | 0.6 | 0.157 | C2 |
| 2.33 | 0.303 | 0.077 | 3.992 | 6.8 | 99.9 | 4.8 | 0.6 | 0.157 | C2 |
| 2.49 | 0.307 | 0.077 | 17.298 | | 5.7 | 98.1 | 5.8 | 0.5 | 0.618 P222 |
| 2.01 | 0.356 | 0.077 | 1.973 | 7.1 | 99.8 | 7.2 | 1.0 | 0.313 | P212121 |
| 2.01 | 0.356 | 0.077 | 1.973 | 7.1 | 99.8 | 7.2 | 1.0 | 0.313 | P212121 |
| 2.99 | 0.536 | 0.077 | 5.355 | 13.1 | 99.9 | 7.3 | 0.6 | 0.224 | P212121 |
| 2.48 | 0.206 | 0.077 | 2.549 | 19.5 | 100.0 | 15.0 | 2.1 | 0.832 | P6322 |
| 3.00 | 0.130 | 0.077 | 2.839 | 11.8 | 100.0 | 15.3 | 1.1 | 0.586 | P41212 |
| 3.52 | 0.307 | 0.077 | 6.045 | 43.8 | 97.2 | 23.6 | 1.8 | 0.796 | P41212 |
| 5.89 | 0.307 | 0.078 | 0.786 | 2.0 | 95.4 | 1.8 | 0.9 | 0.268 | P1 |
| 2.65 | 0.290 | 0.078 | 2.864 | 6.7 | 99.6 | 5.0 | 0.9 | 0.280 | P222 |
| 2.16 | 0.162 | 0.078 | 2.342 | 8.5 | 100.0 | 7.3 | 1.0 | 0.380 | P212121 |
| 1.30 | 0.063 | 0.078 | 0.612 | 3.4 | 82.4 | 10.1 | 1.3 | 0.761 | I121 |
| 3.11 | 0.087 | 0.079 | 0.141 | 1.3 | 6.2 | 1.7 | 1.1 | 0.935 | C2 |
| 1.30 | 0.119 | 0.079 | 2.257 | 12.4 | 99.6 | 10.8 | 0.7 | 0.253 | P222 |
| 1.38 | 0.115 | 0.079 | 1.650 | 13.1 | 100.0 | 12.4 | 1.1 | 0.537 | P212121 |

-------------------------------------------------------------------------

<<< EOF Supplementary_1 815 830 >>>
