## supplementary2 for "I/σI *vs* {Rmerg, Rmeas, Rpim, CC1/2} for Crystal Diffraction Data Quality Evaluation"

**Supplementary 2**: Reduction Statistics of Data Sets Used.
Article Title: **Effective and Reliable Metrics to Select Crystal Diffraction Data**

**trypA**. Data processing statistics by resolution shells.

| Reso | Rmerg | Rmeas | Rpim | CC1/2 | I/σI | Compl% | Multi |
|---|---|---|---|---|---|---|---|
| 8.76 | 0.045 | 0.054 | 0.029 | 0.998 | 24.8 | 88.9 | 4.7 |
| 5.06 | 0.051 | 0.060 | 0.033 | 0.998 | 23.5 | 91.7 | 5.0 |
| 3.92 | 0.048 | 0.059 | 0.033 | 0.997 | 23.7 | 91.5 | 4.5 |
| 3.31 | 0.060 | 0.074 | 0.043 | 0.996 | 19.6 | 92.1 | 4.5 |
| 2.92 | 0.084 | 0.103 | 0.058 | 0.993 | 14.7 | 93.3 | 4.8 |
| 2.64 | 0.107 | 0.130 | 0.073 | 0.987 | 12.2 | 94.3 | 4.8 |
| 2.43 | 0.128 | 0.156 | 0.087 | 0.982 | 10.4 | 95.5 | 4.9 |
| 2.26 | 0.139 | 0.170 | 0.096 | 0.978 | 9.6 | 96.4 | 4.9 |
| 2.13 | 0.151 | 0.185 | 0.104 | 0.975 | 8.6 | 96.8 | 4.8 |
| 2.01 | 0.168 | 0.213 | 0.128 | 0.964 | 6.6 | 96.8 | 4.1 |
| 1.91 | 0.205 | 0.255 | 0.150 | 0.959 | 5.6 | 96.6 | 4.4 |
| 1.83 | 0.263 | 0.328 | 0.192 | 0.924 | 4.3 | 97.1 | 4.5 |
| 1.75 | 0.304 | 0.377 | 0.220 | 0.912 | 3.8 | 97.3 | 4.6 |
| 1.69 | 0.363 | 0.451 | 0.263 | 0.878 | 3.1 | 98.0 | 4.6 |
| 1.63 | 0.404 | 0.504 | 0.295 | 0.851 | 2.7 | 98.3 | 4.6 |
| 1.57 | 0.466 | 0.583 | 0.343 | 0.827 | 2.2 | 99.0 | 4.6 |
| 1.53 | 0.529 | 0.660 | 0.388 | 0.786 | 2.0 | 98.8 | 4.6 |
| 1.48 | 0.563 | 0.721 | 0.443 | 0.735 | 1.6 | 99.2 | 4.0 |
| 1.44 | 0.637 | 0.807 | 0.487 | 0.722 | 1.3 | 99.1 | 4.2 |
| 1.40 | 0.767 | 0.973 | 0.588 | 0.657 | 1.1 | 98.9 | 4.3 |
| 1.37 | 0.853 | 1.082 | 0.654 | 0.622 | 0.9 | 99.2 | 4.3 |
| 1.34 | 0.970 | 1.225 | 0.734 | 0.591 | 0.8 | 99.4 | 4.4 |
| 1.31 | 1.113 | 1.410 | 0.850 | 0.588 | 0.7 | 99.7 | 4.4 |
| 1.28 | 1.180 | 1.496 | 0.903 | 0.464 | 0.5 | 99.5 | 4.3 |
| 1.25 | 1.217 | 1.553 | 0.948 | 0.464 | 0.5 | 97.5 | 4.1 |
| 1.23 | 1.212 | 1.604 | 1.039 | 0.348 | 0.4 | 95.8 | 3.1 |
| 1.20 | 1.245 | 1.701 | 1.152 | 0.360 | 0.3 | 91.8 | 2.6 |
| 1.18 | 1.353 | 1.897 | 1.328 | 0.291 | 0.3 | 83.9 | 2.3 |
| 1.16 | 1.357 | 1.919 | 1.357 | 0.253 | 0.2 | 75.5 | 2.0 |
| 1.14 | 2.078 | 2.938 | 2.078 | 0.202 | 0.2 | 42.1 | 1.5 |

**trypADLS**. DIALS data processing statistics by resolution shells of trypA.

| Reso | Rmerg | Rmeas | Rpim | CC1/2 | I/σI | Compl% | Multi |
|---|---|---|---|---|---|---|---|
| 3.47 | 0.057 | 0.064 | 0.027 | 0.993 | 31.7 | 89.3 | 4.76 |
| 2.76 | 0.085 | 0.094 | 0.040 | 0.994 | 21.7 | 92.1 | 4.78 |
| 2.41 | 0.126 | 0.139 | 0.058 | 0.986 | 15.4 | 94.1 | 4.93 |
| 2.19 | 0.153 | 0.169 | 0.071 | 0.936 | 13.2 | 95.6 | 4.95 |

| | | | | | | | |
|---|---|---|---|---|---|---|---|
| 2.03 | 0.167 | 0.186 | 0.078 | 0.973 | 11.3 | 96.3 | 4.84 |
| 1.91 | 0.199 | 0.226 | 0.104 | 0.955 | 8.6 | 96.0 | 4.13 |
| 1.82 | 0.246 | 0.275 | 0.120 | 0.945 | 7.0 | 96.4 | 4.55 |
| 1.74 | 0.299 | 0.334 | 0.145 | 0.910 | 5.8 | 96.9 | 4.64 |
| 1.67 | 0.346 | 0.386 | 0.168 | 0.889 | 4.9 | 97.7 | 4.65 |
| 1.61 | 0.381 | 0.425 | 0.184 | 0.867 | 4.2 | 97.7 | 4.68 |
| 1.56 | 0.417 | 0.465 | 0.201 | 0.853 | 3.7 | 98.7 | 4.69 |
| 1.52 | 0.473 | 0.528 | 0.229 | 0.801 | 3.2 | 98.9 | 4.68 |
| 1.48 | 0.501 | 0.564 | 0.253 | 0.764 | 2.9 | 98.8 | 4.35 |
| 1.44 | 0.528 | 0.602 | 0.281 | 0.770 | 2.4 | 98.4 | 4.06 |
| 1.41 | 0.586 | 0.660 | 0.296 | 0.710 | 2.2 | 99.0 | 4.39 |
| 1.38 | 0.643 | 0.725 | 0.327 | 0.691 | 1.9 | 98.6 | 4.41 |
| 1.35 | 0.672 | 0.758 | 0.341 | 0.673 | 1.7 | 99.1 | 4.46 |
| 1.33 | 0.705 | 0.792 | 0.353 | 0.611 | 1.7 | 99.2 | 4.52 |
| 1.30 | 0.764 | 0.858 | 0.382 | 0.636 | 1.5 | 99.5 | 4.52 |
| 1.28 | 0.806 | 0.906 | 0.404 | 0.544 | 1.3 | 99.3 | 4.48 |

**trypB30**. Data processing statistics by resolution shells.

| Reso | Rmerg | Rmeas | Rpim | CC1/2 | I/σI | Compl% | Multi |
|---|---|---|---|---|---|---|---|
| 8.38 | 0.011 | 0.015 | 0.009 | 1.000 | 72.3 | 45.5 | 2.4 |
| 4.84 | 0.011 | 0.014 | 0.009 | 0.999 | 67.3 | 48.4 | 2.1 |
| 3.75 | 0.011 | 0.014 | 0.009 | 1.000 | 66.9 | 51.4 | 2.2 |
| 3.17 | 0.012 | 0.015 | 0.009 | 1.000 | 60.0 | 53.4 | 2.2 |
| 2.79 | 0.014 | 0.019 | 0.012 | 0.999 | 49.5 | 54.2 | 2.2 |
| 2.53 | 0.017 | 0.023 | 0.015 | 0.999 | 42.3 | 54.8 | 1.9 |
| 2.33 | 0.019 | 0.025 | 0.017 | 0.999 | 40.8 | 55.1 | 2.0 |
| 2.16 | 0.019 | 0.026 | 0.017 | 0.998 | 36.8 | 56.0 | 2.1 |
| 2.03 | 0.022 | 0.030 | 0.020 | 0.998 | 33.5 | 56.0 | 2.1 |
| 1.92 | 0.023 | 0.032 | 0.022 | 0.998 | 29.5 | 56.3 | 2.1 |
| 1.83 | 0.027 | 0.037 | 0.026 | 0.997 | 22.8 | 55.1 | 1.9 |
| 1.75 | 0.035 | 0.048 | 0.034 | 0.996 | 20.4 | 56.6 | 1.9 |
| 1.68 | 0.040 | 0.057 | 0.039 | 0.996 | 17.7 | 56.2 | 2.0 |
| 1.61 | 0.042 | 0.059 | 0.041 | 0.994 | 15.3 | 56.7 | 2.0 |
| 1.56 | 0.052 | 0.073 | 0.051 | 0.992 | 13.1 | 55.8 | 2.0 |
| 1.51 | 0.054 | 0.077 | 0.054 | 0.991 | 11.7 | 53.8 | 2.1 |
| 1.46 | 0.057 | 0.080 | 0.057 | 0.988 | 9.3 | 45.5 | 1.8 |
| 1.42 | 0.062 | 0.088 | 0.062 | 0.980 | 8.1 | 36.0 | 1.6 |
| 1.38 | 0.079 | 0.112 | 0.079 | 0.980 | 6.6 | 29.3 | 1.6 |
| 1.34 | 0.062 | 0.087 | 0.062 | 0.977 | 6.3 | 17.2 | 1.5 |

**trypB720**. Data processing statistics by resolution shells.

| Reso | Rmerg | Rmeas | Rpim | CC1/2 | I/σI | Compl% | Multi |
|---|---|---|---|---|---|---|---|

| | | | | | | | |
|---|---|---|---|---|---|---|---|
| 8.38 | 0.018 | 0.019 | 0.005 | 1.000 | 174.6 | 99.9 | 24.4 |
| 4.84 | 0.016 | 0.017 | 0.005 | 1.000 | 172.7 | 99.1 | 24.3 |
| 3.75 | 0.017 | 0.017 | 0.005 | 1.000 | 179.2 | 99.8 | 25.8 |
| 3.17 | 0.020 | 0.021 | 0.005 | 1.000 | 161.5 | 100.0 | 27.2 |
| 2.79 | 0.024 | 0.025 | 0.006 | 1.000 | 135.9 | 99.9 | 27.8 |
| 2.53 | 0.026 | 0.027 | 0.007 | 1.000 | 115.9 | 99.4 | 24.3 |
| 2.32 | 0.029 | 0.030 | 0.008 | 1.000 | 113.2 | 99.4 | 26.3 |
| 2.16 | 0.032 | 0.033 | 0.009 | 1.000 | 103.5 | 99.9 | 27.1 |
| 2.03 | 0.035 | 0.036 | 0.010 | 1.000 | 92.7 | 100.0 | 27.4 |
| 1.92 | 0.041 | 0.042 | 0.011 | 1.000 | 80.1 | 100.0 | 27.8 |
| 1.83 | 0.049 | 0.051 | 0.014 | 1.000 | 61.6 | 99.1 | 25.4 |
| 1.75 | 0.058 | 0.060 | 0.016 | 0.999 | 54.3 | 99.6 | 26.0 |
| 1.68 | 0.066 | 0.069 | 0.018 | 0.999 | 47.4 | 99.8 | 26.7 |
| 1.61 | 0.077 | 0.080 | 0.021 | 0.999 | 40.9 | 99.9 | 27.1 |
| 1.56 | 0.092 | 0.096 | 0.025 | 0.999 | 35.1 | 100.0 | 27.1 |
| 1.50 | 0.106 | 0.110 | 0.030 | 0.998 | 29.6 | 99.7 | 26.2 |
| 1.46 | 0.121 | 0.127 | 0.038 | 0.997 | 22.5 | 90.7 | 20.7 |
| 1.42 | 0.139 | 0.147 | 0.047 | 0.995 | 18.2 | 78.3 | 17.7 |
| 1.38 | 0.157 | 0.167 | 0.057 | 0.993 | 14.8 | 67.3 | 15.9 |
| 1.34 | 0.167 | 0.180 | 0.065 | 0.989 | 11.7 | 47.4 | 12.7 |

**Data6OPM**. Data processing statistics by resolution shells for data collected on a crystal entitled "Casposase Bound to Integration Product", (PDB ID: 6OPM). The data were integrated by XDS and scaled by XSCALE. XSCALE doesn't evaluate Rpim. Rpim was evaluated after XSCALE scaling.

| Reso | Rmerg | Rmeas | Rpim | CC1/2 | I/σI | Compl% | Multi |
|---|---|---|---|---|---|---|---|
| 13.86 | 0.045 | 0.047 | 0.014 | 1.000 | 38.1 | 87.9 | 11.3 |
| 9.80 | 0.044 | 0.045 | 0.013 | 0.999 | 39.0 | 100.0 | 12.7 |
| 8.00 | 0.054 | 0.056 | 0.016 | 0.999 | 34.3 | 100.0 | 12.1 |
| 6.93 | 0.067 | 0.070 | 0.019 | 0.998 | 29.7 | 100.0 | 13.7 |
| 6.20 | 0.083 | 0.086 | 0.023 | 0.998 | 25.2 | 100.0 | 14.5 |
| 5.66 | 0.092 | 0.095 | 0.025 | 0.998 | 23.0 | 100.0 | 15.1 |
| 5.24 | 0.093 | 0.096 | 0.025 | 0.998 | 22.8 | 100.0 | 15.1 |
| 4.90 | 0.099 | 0.103 | 0.028 | 0.998 | 19.7 | 100.0 | 13.3 |
| 4.62 | 0.105 | 0.109 | 0.030 | 0.998 | 18.9 | 100.0 | 13.3 |
| 4.38 | 0.123 | 0.127 | 0.034 | 0.998 | 17.5 | 100.0 | 14.2 |
| 4.18 | 0.142 | 0.147 | 0.038 | 0.998 | 15.4 | 100.0 | 14.7 |
| 4.00 | 0.178 | 0.185 | 0.048 | 0.998 | 12.8 | 100.0 | 15.0 |
| 3.85 | 0.248 | 0.257 | 0.066 | 0.997 | 10.7 | 100.0 | 15.3 |
| 3.71 | 0.363 | 0.375 | 0.095 | 0.994 | 8.6 | 99.9 | 15.5 |
| 3.58 | 0.495 | 0.512 | 0.129 | 0.991 | 7.3 | 99.9 | 15.7 |
| 3.47 | 0.642 | 0.664 | 0.169 | 0.988 | 5.9 | 99.9 | 15.4 |
| 3.36 | 0.812 | 0.842 | 0.223 | 0.973 | 4.8 | 99.9 | 14.2 |
| 3.27 | 1.201 | 1.246 | 0.332 | 0.947 | 3.5 | 99.9 | 14.1 |
| 3.18 | 1.732 | 1.794 | 0.466 | 0.910 | 2.7 | 99.9 | 14.8 |

| | | | | | | | |
|---|---|---|---|---|---|---|---|
| 3.10 | 2.391 | 2.475 | 0.641 | 0.819 | 2.0 | 100.0 | 14.9 |

**Data7MRQ**. Data processing statistics by resolution shells for data collected on a crystal entitled "Vertebrate Contactin and Amyloid Precursor Protein", (PDB ID: 7MRQ). The data were processed by HKL2000.

| Reso | Rmerg | Rmeas | Rpim | CC1/2 | I/σI | Compl% | Multi |
|---|---|---|---|---|---|---|---|
| 7.65 | 0.064 | 0.070 | 0.028 | 0.991 | 18.3 | 99.2 | 6.5 |
| 6.08 | 0.079 | 0.086 | 0.032 | 0.994 | 18.3 | 100.0 | 7.0 |
| 5.31 | 0.098 | 0.106 | 0.040 | 0.993 | 17.9 | 100.0 | 7.2 |
| 4.82 | 0.107 | 0.116 | 0.043 | 0.991 | 17.6 | 100.0 | 7.2 |
| 4.48 | 0.126 | 0.136 | 0.050 | 0.991 | 15.6 | 100.0 | 7.2 |
| 4.21 | 0.147 | 0.159 | 0.059 | 0.989 | 14.1 | 100.0 | 7.2 |
| 4.00 | 0.185 | 0.199 | 0.074 | 0.984 | 12.0 | 100.0 | 7.2 |
| 3.83 | 0.236 | 0.255 | 0.094 | 0.981 | 9.5 | 100.0 | 7.0 |
| 3.68 | 0.300 | 0.324 | 0.119 | 0.979 | 6.8 | 99.7 | 6.9 |
| 3.55 | 0.365 | 0.394 | 0.146 | 0.965 | 5.4 | 98.5 | 6.7 |
| 3.44 | 0.413 | 0.446 | 0.166 | 0.952 | 4.6 | 94.6 | 6.5 |
| 3.34 | 0.503 | 0.544 | 0.204 | 0.935 | 3.3 | 90.7 | 6.3 |
| 3.26 | 0.614 | 0.665 | 0.252 | 0.910 | 2.3 | 86.0 | 5.9 |
| 3.18 | 0.740 | 0.806 | 0.313 | 0.845 | 2.0 | 81.7 | 5.6 |
| 3.10 | 0.870 | 0.950 | 0.374 | 0.843 | 1.5 | 73.6 | 5.4 |
| 3.04 | 1.125 | 1.236 | 0.501 | 0.646 | 1.1 | 63.9 | 5.2 |
| 2.98 | 1.181 | 1.309 | 0.549 | 0.479 | 0.9 | 60.7 | 4.8 |
| 2.92 | 1.577 | 1.766 | 0.772 | 0.321 | 0.7 | 51.9 | 4.3 |
| 2.87 | 1.596 | 1.807 | 0.823 | 0.189 | 0.6 | 46.5 | 3.8 |
| 2.82 | 1.652 | 1.902 | 0.909 | 0.25 | 0.6 | 36.8 | 3.5 |

**Data8D7K**. Data processing statistics by resolution shells for data collected on a crystal entitled "Bifunctional Inhibition of Neutrophil Elastase and Cathepsin G by Eap2 from S. aureus" (PDB ID: 8D7K).

| Reso | Rmerg | Rmeas | Rpim | CC1/2 | I/σI | Compl% | Multi |
|---|---|---|---|---|---|---|---|
| 18.02 | 0.045 | 0.056 | 0.033 | 0.996 | 26.6 | 93.8 | 4.3 |
| 10.41 | 0.045 | 0.056 | 0.034 | 0.995 | 28.8 | 88.3 | 4.7 |
| 8.06 | 0.057 | 0.073 | 0.045 | 0.986 | 20.5 | 88.7 | 4.0 |
| 6.81 | 0.072 | 0.091 | 0.056 | 0.978 | 16.7 | 89.2 | 4.2 |
| 6.01 | 0.092 | 0.117 | 0.071 | 0.965 | 13.8 | 93.3 | 4.3 |
| 5.43 | 0.111 | 0.141 | 0.086 | 0.955 | 13.1 | 93.1 | 4.5 |
| 5.00 | 0.097 | 0.124 | 0.075 | 0.934 | 14.0 | 93.4 | 4.5 |
| 4.65 | 0.110 | 0.140 | 0.085 | 0.913 | 13.2 | 94.5 | 4.5 |
| 4.37 | 0.125 | 0.159 | 0.097 | 0.945 | 11.7 | 96.2 | 4.5 |
| 4.14 | 0.168 | 0.216 | 0.132 | 0.928 | 9.3 | 96.3 | 4.4 |
| 3.93 | 0.240 | 0.310 | 0.194 | 0.868 | 6.7 | 95.6 | 4.0 |
| 3.76 | 0.328 | 0.419 | 0.257 | 0.861 | 5.8 | 98.0 | 4.5 |

| | | | | | | | |
|---|---|---|---|---|---|---|---|
| 3.60 | 0.425 | 0.539 | 0.327 | 0.849 | 4.8 | 98.4 | 4.6 |
| 3.47 | 0.576 | 0.730 | 0.442 | 0.754 | 3.7 | 99.3 | 4.8 |
| 3.35 | 0.697 | 0.878 | 0.527 | 0.673 | 2.9 | 99.5 | 4.9 |
| 3.24 | 0.971 | 1.225 | 0.736 | 0.539 | 2.0 | 99.8 | 4.9 |
| 3.14 | 1.281 | 1.613 | 0.965 | 0.426 | 1.5 | 99.9 | 5.0 |
| 3.05 | 1.463 | 1.844 | 1.105 | 0.367 | 1.2 | 99.8 | 5.0 |
| 2.96 | 1.731 | 2.179 | 1.304 | 0.250 | 1.0 | 99.8 | 5.1 |
| 2.89 | 2.399 | 3.022 | 1.812 | 0.197 | 0.7 | 99.8 | 5.1 |

**Data9ATU**. Data processing statistics by resolution shells for data collected on a crystal entitled "Bifunctional Inhibition of Neutrophil Elastase by Eap4 from S. aureus" (PDB ID: 9ATU).

| Reso | Rmerg | Rmeas | Rpim | CC1/2 | I/$\sigma$I | Compl% | Multi |
|---|---|---|---|---|---|---|---|
| 4.11 | 0.078 | 0.081 | 0.022 | 0.986 | 23.0 | 99.4 | 13.2 |
| 3.27 | 0.077 | 0.080 | 0.021 | 0.997 | 26.1 | 99.3 | 13.3 |
| 2.85 | 0.116 | 0.121 | 0.033 | 0.996 | 20.6 | 99.9 | 13.4 |
| 2.59 | 0.202 | 0.209 | 0.055 | 0.991 | 12.8 | 99.9 | 14.0 |
| 2.41 | 0.438 | 0.454 | 0.119 | 0.981 | 8.2 | 100.0 | 14.1 |
| 2.26 | 0.564 | 0.588 | 0.166 | 0.936 | 5.0 | 99.3 | 12.3 |
| 2.15 | 0.881 | 0.917 | 0.253 | 0.887 | 3.0 | 100.0 | 12.8 |
| 2.03 | 1.250 | 1.312 | 0.389 | 0.476 | 1.4 | 99.5 | 10.5 |
| 1.95 | 1.587 | 1.698 | 0.571 | 0.275 | 0.7 | 93.4 | 6.7 |
| 1.91 | 1.817 | 1.976 | 0.728 | 0.190 | 0.5 | 74.2 | 4.3 |

**ANSdd125**. Data processing statistics by AIMLESS

Analysis of anisotropy of data
===============================
Principal axes:
d1: 0.75 a* - 0.66 c* ~= hkl direction 1 0 -1
d2: k axis
d3: 0.32 a* + 0.95 c* ~= hkl direction 1 0 3
Eigenvalues of [B](orth) along principal axes : -11.173 7.362 3.811
Difference between maximum and minimum anisotropic B (= 8 pi^2 U) 18.5
Estimated maximum resolution limits, d1: 1.95, d2: 1.72, d3: 1.72

| Reso | Rmerg | Rmeas | Rpim | CC1/2 | I/$\sigma$I | Compl% | Multi |
|---|---|---|---|---|---|---|---|
| 10.88 | 0.032 | 0.038 | 0.020 | 0.999 | 46.0 | 99.1 | 6.3 |
| 6.28 | 0.039 | 0.046 | 0.024 | 0.998 | 42.8 | 100.0 | 6.7 |
| 4.86 | 0.034 | 0.042 | 0.023 | 0.999 | 41.6 | 99.0 | 5.9 |
| 4.11 | 0.035 | 0.042 | 0.023 | 0.999 | 41.2 | 99.6 | 6.1 |
| 3.63 | 0.040 | 0.048 | 0.026 | 0.998 | 39.8 | 99.8 | 6.4 |
| 3.28 | 0.044 | 0.052 | 0.028 | 0.998 | 36.4 | 99.8 | 6.7 |
| 3.02 | 0.055 | 0.065 | 0.034 | 0.998 | 30.6 | 99.9 | 6.8 |
| 2.81 | 0.066 | 0.077 | 0.041 | 0.997 | 25.8 | 99.9 | 7.1 |

| | | | | | | | |
|---|---|---|---|---|---|---|---|
| 2.64 | 0.081 | 0.096 | 0.051 | 0.996 | 21.2 | 100.0 | 6.8 |
| 2.50 | 0.095 | 0.115 | 0.063 | 0.995 | 16.7 | 100.0 | 6.4 |
| 2.37 | 0.117 | 0.140 | 0.076 | 0.995 | 14.1 | 100.0 | 6.5 |
| 2.27 | 0.144 | 0.171 | 0.091 | 0.994 | 11.9 | 100.0 | 6.9 |
| 2.18 | 0.166 | 0.196 | 0.103 | 0.994 | 10.5 | 100.0 | 7.0 |
| 2.09 | 0.217 | 0.256 | 0.135 | 0.988 | 8.3 | 100.0 | 7.0 |
| 2.02 | 0.260 | 0.306 | 0.160 | 0.985 | 6.8 | 100.0 | 7.1 |
| 1.95 | 0.375 | 0.440 | 0.229 | 0.967 | 4.8 | 100.0 | 7.1 |
| 1.89 | 0.514 | 0.604 | 0.314 | 0.938 | 3.3 | 100.0 | 7.1 |
| 1.84 | 0.616 | 0.730 | 0.389 | 0.903 | 2.5 | 99.2 | 6.7 |
| 1.79 | 0.801 | 0.981 | 0.557 | 0.799 | 1.6 | 96.4 | 5.8 |
| 1.74 | 0.982 | 1.237 | 0.738 | 0.699 | 1.2 | 87.4 | 4.9 |

**ANSdd227**. Data processing statistics by AIMLESS

Analysis of anisotropy of data
==============================
Principal axes:
d1: 0.97 a* - 0.24 c* ~= hkl direction 4 0 -1
d2: k axis
d3: 0.08 a* + 1.00 c* ~= hkl direction 1 0 12
Eigenvalues of [B](orth) along principal axes : 1.572 7.488 -9.061
Difference between maximum and minimum anisotropic B (= 8 pi^2 U) 16.5
Estimated maximum resolution limits, d1: 1.76, d2: 2.21, d3: 1.96

| Reso | Rmerg | Rmeas | Rpim | CC1/2 | I/σI | Compl% | Multi |
|---|---|---|---|---|---|---|---|
| 11.13 | 0.064 | 0.081 | 0.048 | 0.994 | 17.5 | 98.9 | 4.3 |
| 6.43 | 0.069 | 0.088 | 0.054 | 0.992 | 16.8 | 99.5 | 4.7 |
| 4.98 | 0.072 | 0.091 | 0.056 | 0.991 | 16.9 | 98.9 | 4.6 |
| 4.21 | 0.078 | 0.102 | 0.065 | 0.987 | 14.6 | 98.6 | 3.8 |
| 3.71 | 0.085 | 0.111 | 0.070 | 0.987 | 13.6 | 98.7 | 4.0 |
| 3.36 | 0.098 | 0.128 | 0.081 | 0.987 | 12.4 | 98.6 | 4.4 |
| 3.09 | 0.135 | 0.177 | 0.113 | 0.978 | 10.2 | 98.7 | 4.5 |
| 2.87 | 0.171 | 0.226 | 0.146 | 0.971 | 8.8 | 98.7 | 4.5 |
| 2.70 | 0.247 | 0.329 | 0.215 | 0.940 | 7.5 | 98.6 | 4.6 |
| 2.55 | 0.333 | 0.445 | 0.292 | 0.898 | 6.3 | 98.4 | 4.7 |
| 2.43 | 0.411 | 0.547 | 0.358 | 0.887 | 5.4 | 98.3 | 4.7 |
| 2.32 | 0.525 | 0.701 | 0.459 | 0.750 | 4.3 | 98.1 | 4.3 |
| 2.23 | 0.639 | 0.853 | 0.559 | 0.641 | 3.8 | 97.8 | 4.1 |
| 2.14 | 0.708 | 0.938 | 0.608 | 0.641 | 3.4 | 97.9 | 4.4 |
| 2.07 | 0.840 | 1.107 | 0.711 | 0.624 | 3.0 | 97.6 | 4.5 |
| 2.00 | 1.033 | 1.346 | 0.851 | 0.541 | 2.4 | 97.4 | 4.6 |
| 1.94 | 1.146 | 1.480 | 0.924 | 0.574 | 2.1 | 97.4 | 4.7 |
| 1.88 | 1.399 | 1.794 | 1.106 | 0.477 | 1.6 | 96.9 | 4.8 |
| 1.83 | 1.624 | 2.071 | 1.268 | 0.498 | 1.3 | 96.9 | 4.8 |
| 1.78 | 1.917 | 2.448 | 1.502 | 0.353 | 1.0 | 96.7 | 4.9 |

**ANSdd313**. Data processing statistics by AIMLESS

Analysis of anisotropy of data
=============================

Principal axes:
d1: 0.95 a* + 0.30 c* ~= hkl direction 3 0 1
d2: k axis
d3: -0.56 a* + 0.83 c* ~= hkl direction -1 0 1
Eigenvalues of [B](orth) along principal axes : 4.972 -14.578 9.606
Difference between maximum and minimum anisotropic B (= 8 pi^2 U) 24.2
Estimated maximum resolution limits, d1: 1.97, d2: 2.70, d3: 1.97

| Reso | Rmerg | Rmeas | Rpim | CC1/2 | I/σI | Compl% | Multi |
|---|---|---|---|---|---|---|---|
| 12.46 | 0.049 | 0.069 | 0.049 | 0.993 | 18.0 | 97.7 | 3.0 |
| 7.19 | 0.054 | 0.076 | 0.053 | 0.994 | 17.2 | 98.5 | 3.4 |
| 5.57 | 0.059 | 0.082 | 0.057 | 0.992 | 16.1 | 98.5 | 3.4 |
| 4.71 | 0.054 | 0.075 | 0.052 | 0.993 | 15.8 | 98.9 | 2.9 |
| 4.15 | 0.057 | 0.079 | 0.055 | 0.993 | 15.3 | 99.3 | 3.1 |
| 3.76 | 0.066 | 0.092 | 0.063 | 0.992 | 13.9 | 99.1 | 3.3 |
| 3.46 | 0.080 | 0.110 | 0.075 | 0.991 | 11.9 | 99.5 | 3.4 |
| 3.22 | 0.098 | 0.135 | 0.092 | 0.990 | 10.0 | 99.7 | 3.4 |
| 3.02 | 0.129 | 0.177 | 0.120 | 0.986 | 7.9 | 99.9 | 3.5 |
| 2.86 | 0.174 | 0.238 | 0.161 | 0.974 | 6.2 | 99.8 | 3.5 |
| 2.72 | 0.216 | 0.295 | 0.199 | 0.966 | 5.1 | 100.0 | 3.6 |
| 2.60 | 0.257 | 0.351 | 0.238 | 0.947 | 4.1 | 99.9 | 3.3 |
| 2.49 | 0.300 | 0.410 | 0.279 | 0.924 | 3.4 | 100.0 | 3.2 |
| 2.40 | 0.371 | 0.505 | 0.340 | 0.893 | 2.8 | 99.9 | 3.1 |
| 2.31 | 0.441 | 0.600 | 0.405 | 0.887 | 2.5 | 99.9 | 3.4 |
| 2.24 | 0.536 | 0.727 | 0.489 | 0.854 | 2.1 | 99.9 | 3.4 |
| 2.17 | 0.645 | 0.877 | 0.591 | 0.819 | 1.7 | 99.8 | 3.5 |
| 2.11 | 0.800 | 1.087 | 0.731 | 0.704 | 1.4 | 99.9 | 3.5 |
| 2.05 | 1.121 | 1.521 | 1.021 | 0.601 | 1.0 | 99.9 | 3.5 |
| 2.00 | 1.290 | 1.749 | 1.174 | 0.579 | 0.9 | 99.8 | 3.5 |

**ANSdd374**. Data processing statistics by AIMLESS

Analysis of anisotropy of data
=============================

Principal axes:
Principal axes:
d1: 0.96 a* - 0.27 c* ~= hkl direction 4 0 -1
d2: 1.00 b* + 0.00 c* ~= hkl direction 0 1 0
d3: 0.51 a* + 0.86 c* ~= hkl direction 1 0 2
Eigenvalues of [B](orth) along principal axes : -8.650 1.080 7.570

Difference between maximum and minimum anisotropic B (= 8 pi^2 U) 16.2
Estimated maximum resolution limits, d1: 4.00, d2: 2.74, d3: 3.01

| Reso | Rmerg | Rmeas | Rpim | CC1/2 | I/σI | Compl% | Multi |
|---|---|---|---|---|---|---|---|
| 17.33 | 0.125 | 0.177 | 0.125 | 0.971 | 20.3 | 99.6 | 3.0 |
| 10.01 | 0.143 | 0.202 | 0.142 | 0.941 | 21.9 | 99.9 | 3.5 |
| 7.75 | 0.169 | 0.238 | 0.168 | 0.932 | 15.8 | 100.0 | 3.5 |
| 6.55 | 0.219 | 0.305 | 0.212 | 0.915 | 11.8 | 100.0 | 3.6 |
| 5.78 | 0.300 | 0.416 | 0.287 | 0.917 | 9.9 | 100.0 | 3.6 |
| 5.23 | 0.297 | 0.414 | 0.287 | 0.903 | 10.0 | 99.8 | 3.3 |
| 4.81 | 0.301 | 0.419 | 0.290 | 0.893 | 10.3 | 99.8 | 3.1 |
| 4.47 | 0.468 | 0.651 | 0.452 | 0.710 | 8.4 | 99.7 | 2.9 |
| 4.20 | 0.493 | 0.686 | 0.476 | 0.692 | 8.3 | 99.8 | 3.2 |
| 3.98 | 0.562 | 0.785 | 0.547 | 0.631 | 7.3 | 100.0 | 3.3 |
| 3.78 | 0.802 | 1.114 | 0.771 | 0.431 | 5.6 | 99.8 | 3.4 |
| 3.61 | 0.933 | 1.293 | 0.892 | 0.334 | 4.8 | 99.9 | 3.4 |
| 3.47 | 0.999 | 1.381 | 0.949 | 0.475 | 3.8 | 99.8 | 3.5 |
| 3.34 | 1.166 | 1.604 | 1.097 | 0.556 | 2.7 | 99.9 | 3.5 |
| 3.22 | 1.360 | 1.869 | 1.275 | 0.426 | 2.1 | 100.0 | 3.5 |
| 3.11 | 1.421 | 1.952 | 1.331 | 0.444 | 1.7 | 100.0 | 3.6 |
| 3.02 | 1.652 | 2.266 | 1.543 | 0.466 | 1.4 | 100.0 | 3.6 |
| 2.93 | 2.586 | 3.544 | 2.412 | 0.221 | 0.9 | 100.0 | 3.6 |
| 2.85 | 2.504 | 3.434 | 2.339 | 0.209 | 0.9 | 100.0 | 3.6 |
| 2.77 | 2.896 | 3.968 | 2.698 | 0.188 | 0.7 | 100.0 | 3.6 |

**MERG10**. Data processing statistics by XIA2.MULTIPLEX

| Reso | Rmerg | Rmeas | Rpim | CC1/2 | I/σI | Compl% | Multi |
|---|---|---|---|---|---|---|---|
| 2.86 | 0.058 | 0.063 | 0.023 | 0.997 | 39.9 | 99.23 | 6.79 |
| 2.27 | 0.077 | 0.083 | 0.030 | 0.995 | 32.7 | 99.52 | 7.43 |
| 1.98 | 0.092 | 0.100 | 0.036 | 0.992 | 27.2 | 99.52 | 7.10 |
| 1.80 | 0.113 | 0.122 | 0.044 | 0.990 | 21.6 | 99.17 | 7.13 |
| 1.67 | 0.140 | 0.150 | 0.053 | 0.986 | 17.6 | 99.31 | 7.38 |
| 1.58 | 0.160 | 0.172 | 0.062 | 0.979 | 14.5 | 99.39 | 7.22 |
| 1.50 | 0.189 | 0.204 | 0.075 | 0.974 | 11.9 | 99.14 | 6.88 |
| 1.43 | 0.212 | 0.231 | 0.088 | 0.964 | 10.1 | 99.03 | 6.38 |
| 1.38 | 0.243 | 0.263 | 0.098 | 0.956 | 8.8 | 99.55 | 6.63 |
| 1.33 | 0.278 | 0.300 | 0.111 | 0.948 | 7.8 | 99.48 | 6.76 |
| 1.29 | 0.306 | 0.330 | 0.122 | 0.942 | 7.0 | 99.63 | 6.73 |
| 1.25 | 0.333 | 0.361 | 0.136 | 0.933 | 6.0 | 99.04 | 6.30 |
| 1.22 | 0.344 | 0.383 | 0.162 | 0.920 | 4.7 | 95.58 | 4.83 |
| 1.19 | 0.363 | 0.411 | 0.189 | 0.870 | 3.9 | 88.45 | 3.79 |
| 1.16 | 0.382 | 0.439 | 0.212 | 0.860 | 3.3 | 77.07 | 3.21 |

<<< EOF>>
